# Multimodal sensory control of motor performance by glycinergic interneurons of the spinal cord deep dorsal horn

**DOI:** 10.1101/2022.05.21.492933

**Authors:** Mark A Gradwell, Nofar Ozeri-Engelhard, Jaclyn T Eisdorfer, Olivier D Laflamme, Melissa Gonzalez, Aman Upadhyay, Adin Aoki, Tara Shrier, Melissa Gandhi, Gloria Abbas-Zadeh, Olisemaka Oputa, Joshua K Thackray, Matthew Ricci, Nusrath Yusuf, Jessica Keating, Manon Bohic, Zarghona Imtiaz, Simona A Alomary, Jordan Katz, Michael Haas, Yurdiana Hernandez, Turgay Akay, Victoria Abraira

## Abstract

To achieve smooth motor performance in a changing sensory environment, motor outputs must be constantly updated in response to sensory feedback. Inhibitory interneurons in the spinal cord play an essential role in shaping motor activity by gating the transmission of sensory information and setting the pattern and rhythm of motor neurons. Here, we identify the medial deep dorsal horn of the spinal cord as a “hot zone” of convergent proprioceptive and cutaneous information from the hindlimb, where inhibitory neurons show increased responsiveness to sensory input and are preferentially recruited during locomotion in comparison to excitatory neurons. We identify a novel population of glycinergic inhibitory neurons within the deep dorsal horn that express parvalbumin (dPV) and receive convergent proprioceptive and cutaneous input from the paw. We show that dPVs possess intrinsic properties that support spontaneous discharge, even in the absence of synaptic input. However, a drug cocktail mimicking descending input (5-HT, dopamine, NMDA) amplifies dPV output, while cutaneous and proprioceptive inputs shape the temporal dynamics of dPV activity. These findings suggest dPV-mediated inhibition is modulated by behavioral state and can be fine-tuned by sensory input. Using intersectional genetic strategies, we selectively target spinal cord dPVs and demonstrate their capacity to provide divergent ipsilateral inhibition to both pre-motor and motor networks of the ventral horn, thereby controlling the timing and magnitude of cutaneous-evoked muscle activity. Manipulating the activity of dPVs during treadmill locomotion results in altered limb kinematics at the transition of stance to swing and altered step cycle timing at increased speeds. To investigate the effects of manipulating dPV activity on broader sets of motor behaviors, we used depth vision and machine learning to quantify and scale naturalistic behavior. We find that although sub-movements remain stable, the transitions between sub-movements are reduced, suggesting a role in movement switching. In sum, our study reveals a new model by which sensory convergence and inhibitory divergence produce a surprisingly flexible influence on motor networks to increase the diversity of mechanisms by which sensory input facilitates smooth movement and context-appropriate transitions.

**Highlights:**

- Inhibitory deep dorsal horn interneurons integrate convergent proprioceptive and cutaneous sensory inputs from the paw and are preferentially recruited during locomotion.
- Deep dorsal horn parvalbumin+ interneurons (dPVs) represent a population of glycinergic interneurons that can provide sustained inhibitory control.
- Sensory input engages dPVs to facilitate inhibition with high temporal precision and reduced variability.
- dPVs contribute to the ipsilateral inhibitory control of motor and premotor networks of the ventral horn, thereby gating the magnitude and timing of cutaneous-evoked flexor and extensor muscle activity.
- *In vivo*, dPVs modulate gait dynamics in a state- and phase-dependent manner, to ensure smooth movement transitions between step-cycle phases and naturalistic sub-movements.

## INTRODUCTION

Intrinsic spinal networks generate complex movements by setting the pattern and rhythm of muscle activity, defining the activation profile of muscles, and governing parameters of locomotor activity such as the timing of stance to swing transitions. This rhythmic, patterned activity can be coordinated within limb’s joints (intralimb) and across limbs (interlimb); the interplay of which determines locomotion gait. To adapt to changes in the environment, such as varying surfaces and speeds, it is essential for gait parameters (rhythm, pattern, and coordination) and limb kinematics (angles of joints) to be adjustable. The foundations of our understanding of locomotion networks can be attributed to the work of Charles Sherrington and Graham Brown in the early 20th century^1,2^. Sherrington posited that locomotion operates through a series of reflexive “chains”, and Brown built upon this work to suggest the existence of intrinsic spinal networks that can generate “the act of progression” in the absence of sensory input. Over time, our comprehension has evolved, revealing functionally organized networks intrinsic to the spinal cord that drive the rhythm and pattern of locomotion^3–8^. Although sensory afferents are not essential for locomotor production, they play a crucial role in shaping both gait and limb kinematics to achieve smooth and adaptable movements. However, the physiological properties of sensorimotor circuits that allow for crosstalk between sensory signals and motor pools, thereby enabling flexible modulation of pre-set motor programs remains an open question. To approximate a holistic understanding of the spinal networks of locomotion, we need a functional roadmap that connects multimodal sensory information from individual limbs to the motor pathways capable of coordinating complex movements.

When walking, the spinal cord receives constant feedback information from two primary sensory modalities. The first, proprioception, provides information on muscle state (tension, speed, and location) and is detected by receptors located in muscles, joints, and tendons. The second, touch, relays information about skin indentation, vibration, and slip, and is detected by cutaneous afferents, low threshold mechanoreceptors (LTMRs), innervating specialized mechanosensitive organs embedded in skin, bones, and joints such as Merkel cell, Ruffini endings, Meissner and Pacinian corpuscles^9^. Proprioceptive and cutaneous input work in concert to shape the rhythm of locomotion by adjusting the duration of each phase of the step cycle^10–12^. For example, during the swing phase, activation of mechanoreceptors on the dorsal paw elicits a flexor response^13,14^. In contrast, applying the same stimuli during the stance phase promotes extension^15^. Such findings beg the question of whether proprioceptive and cutaneous input to the spinal cord use parallel or segregated pathways to modulate motor output. Studies have shown that activation of cutaneous inputs can facilitate proprioceptive reflexes, suggesting a convergence of proprioceptive and cutaneous pathways^16,17^. This facilitation exceeds a simple summation of each input and is thought to occur through convergence onto common pathways^18^.

Anatomical and physiological studies indicate that processing of convergent proprioceptive and cutaneous input occurs in the medial part of the deep dorsal horn of the spinal cord^19–22^. This region is also home to a large population of premotor interneurons^23–28^,“reflex encoders’’ that participate in cutaneous withdrawal reflexes^29,30^, and “motor synergy encoders’’ that coordinate motor response across multiple muscle groups ^31^. Consistent with this, coordinated motor output can most efficiently be elicited by electrical stimulation of the medial deep dorsal horn^32^. Together, these studies indicate that interneurons located in the medial deep dorsal horn possess the ability to integrate multimodal sensory input and generate motor output, thereby engaging in multimodal sensorimotor transformations. Indeed, both excitatory and inhibitory deep dorsal horn interneurons have been implicated in multimodal sensorimotor processing. For instance, the excitatory dl3 pre-motor interneurons integrate proprioceptive and LTMR input to control grasping^33^. Additionally, inhibitory interneurons expressing the Satb2 transcription factor integrate proprioceptive and noxious input to regulate the nociceptive withdrawal reflex^34^. Inhibitory interneurons that transiently express RORβ provide presynaptic inhibition of proprioceptive afferents to ensure a fluid gait during locomotion^35^. Collectively, these studies show that deep dorsal horn excitatory and inhibitory interneurons are key players in sensorimotor responses required for coordinated motor activity. However, what are the input-output circuit maps underlying their role in sensorimotor control for locomotion?

Here we corroborate that the medial portion of lamina IV-V in the mouse spinal cord (i.e., the deep dorsal horn) is a site of high convergence for proprioceptive and cutaneous input. We find that inhibitory interneurons, due to their increased responsiveness to sensory input, are preferentially recruited during locomotion. We next identify a large, previously uncharacterized, population of inhibitory interneurons in the deep dorsal horn that express the calcium-binding protein parvalbumin (dPVs). We show that dPVs integrate both cutaneous and proprioceptive sensory input and provide divergent output to premotor and motor neurons. This circuit map supports a role in multimodal sensorimotor processing that regulates motor activity in a flexible, context-dependent manner. Utilizing intersectional genetics, we ablate dPVs in adult mice to show that loss of dPV-mediated inhibition 1) reduces cutaneous-evoked inhibition of muscle response latency and amplitude, 2) impacts hindlimb kinematics and step cycle dynamics, particularly during the transition from stance to swing phase during forced locomotion, and 3) reduces transition probabilities between sub-movements during naturalistic behavior. Taken together, these findings identify dPV neurons as key players in sensory-evoked inhibition of ventral horn motor networks to orchestrate the smooth transition between both step-cycle phases and naturalistic sub-movements.

## RESULTS

### The medial deep dorsal horn is defined by convergent proprioceptive and cutaneous input

Building on previous work indicating that sensory inputs from both muscle and skin are involved in shaping motor behavior^36^, we sought to identify the specific region of the spinal cord where proprioceptive and cutaneous sensory information converge within the spinal cord. To this end, we used *PV^2aCreER^*;*Advillin^FlpO^*; *R26^LSL-FSF-TdTomato^ (Ai65)*; *R26^FSF-SynGFP^* mice, in which proprioceptive and cutaneous inputs are genetically labeled with different fluorophores. In this cross, proprioceptive input is labeled with both green and red fluorescent proteins (seen as yellow), while cutaneous (non-proprioceptive input) is labeled with only a red fluorescent protein (**Figure 1A**). We found that cutaneous inputs are concentrated in the superficial and deep dorsal horn (Lamina I-IV), whereas proprioceptive inputs span the deep dorsal and ventral horns (Lamina V-X). Notably, the convergence of proprioceptive and cutaneous input was largely restricted to the medial deep dorsal horn (mDDH, LIV medial-LV medial), highlighting the significance of this specific region of the spinal cord for multimodal sensory processing (**Figure 1B**).

**Figure 1.**
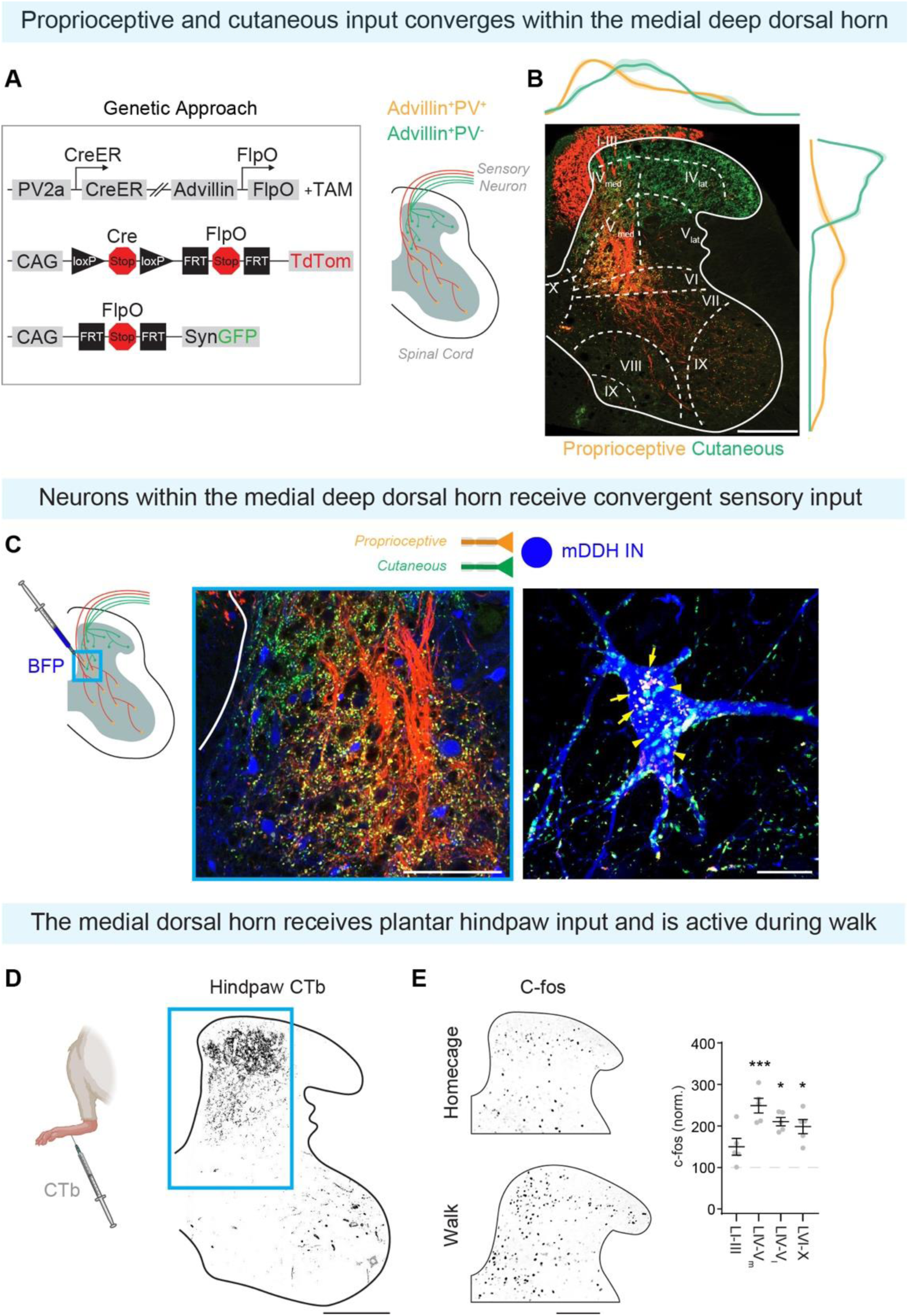
The medial deep dorsal horn is defined by convergent proprioceptive and cutaneous input and is recruited by walking. **(A)** Quadruple transgenic approach, *PV^2aCreER^;Advillin^FlpO^;Rosa^LSL-FSF-Tomato^/Rosa^FSF-SynGFP^,* to label cutaneous (green) and proprioceptive (red) afferent terminals. **(B)** Transverse section of lumbar spinal cord from *PV^2aCreER^;Advillin^FlpO^;Rosa^LSL-FSF-Tomato^/Rosa^FSF-SynGFP^* mouse, including medio-lateral and dorso-ventral density distributions. **(C)** Left: mDDH of BFP injected spinal cord from *PV^2aCreER^;Advillin^FlpO^;Rosa^LSL-FSF-Tomato^/Rosa^FSF-SynGFP^* mouse. Right: BFP labeled neuron receiving convergent proprioceptive (arrows) and cutaneous (arrowheads) afferent input. **(D)** Transverse section of lumbar spinal cord following CTb injection into the ventral paw. **(E)** Left: c-fos expression in lumbar spinal cord dorsal horn of mice from ‘homecage’ and ‘walk’ conditions. Right: Quantification of c-fos as percentage of homecage across lamina groups (LI-III, LIV-V_Medial_, LIV-V_Lateral_, and LVI-X). Scale bar: 200 µm, 20 µm for C right. For further details on genetic crosses and statistical tests see Methods.

To test whether individual interneurons within the mDDH receive convergent cutaneous and proprioceptive input, we employed a viral approach (AAV8-CAG2-eBFP2-wpre) to label individual interneurons in *PV^2aCreER^*;*Advillin^FlpO^*;*R26^LSL-FSF-TdTomato^*; *R26^FSF-SynGFP^* quadruple transgenic mice (**Figure 1C**). To distinguish inhibitory (Pax2^+^) and excitatory (Pax2^-^) mDDH interneurons, we performed Pax2 immunostaining (**Figure S1A)**. Our analysis of synaptic input from proprioceptive and cutaneous afferents revealed that both inhibitory and excitatory interneurons within the mDDH process convergent sensory input (**Figure S1A**). The medial position of this region suggests integration of inputs from the distal limb (i.e., paw^37–39)^. To address this, we injected the anterograde tracer CTb into the ventral hindpaw and found that paw innervating primary sensory afferents concentrated in the medial portion of the dorsal horn (**Figure 1D**). Together, these findings suggest that individual interneurons of the medial deep dorsal horn (mDDH) integrate convergent proprioceptive and cutaneous sensory inputs from the paw.

It is well understood that weight-bearing overground locomotion recruits both proprioceptive and cutaneous afferents^40,41^. Based on the convergence of multimodal sensory information from the paw, we expect that the mDDH would be more activated during walking. Thus, we examined the expression of the early immediate gene c-fos, a correlate of neural activity, during treadmill locomotion^42,43^. As expected, we observed an increase in c-fos positive neurons throughout the deep and ventral laminas, with a distinct locus in the medial region of Lamina IV-V (mDDH, **Figure 1E**). These data corroborate our histological observations that the medial deep dorsal horn is a unique region, activated by convergent proprioceptive and cutaneous inputs during weight-supported locomotion.

### Dorsal horn inhibitory neurons are preferentially recruited during walking

We next asked whether the increase in activity during overground walking was restricted to either excitatory or inhibitory interneurons of the deep dorsal horn. To do this we used *vGAT^iresCre^;Tau^LSL-F-nLacZ-F-DTR^ (Tau^ds-DTR^)* mice and quantified c-fos in inhibitory (c-fos^+^/LacZ+) and excitatory (c-fos^+^/LacZ^-^) neurons. Our findings demonstrate that overground walking primarily activates inhibitory neurons in the medial deep dorsal horn (**Figure 2A**). In line with preferential inhibitory neuron recruitment during walking, we found that optogenetic activation of sensory afferents was more likely to evoke action potentials (APs) in inhibitory neurons compared to excitatory neurons (**Figure 2B**), despite receiving similar amplitude inputs (**Figure S1B**). This observation suggests that inhibitory interneurons possess more excitable intrinsic properties than their excitatory counterparts, which could contribute to their greater involvement in processing sensory input during walking.

**Figure 2.**
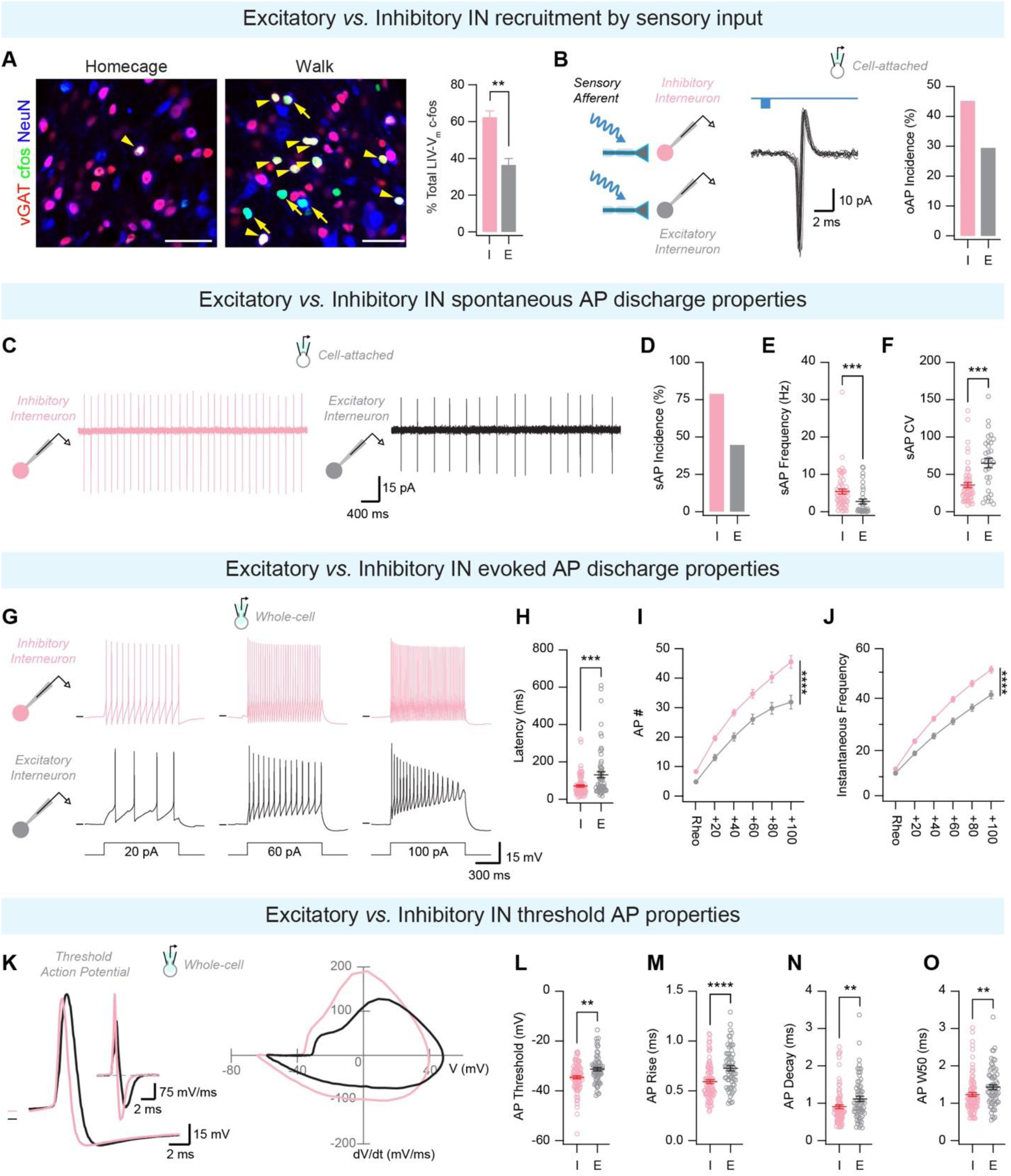
DDH inhibitory interneurons are preferentially recruited during walking due to their excitable electrophysiological properties. **(A)** Images of LIV_Medial_-V_Medial_ from ‘homecage’ (left) and ‘walk’ (right) mice. Neurons are in blue, c-fos labeling is in green, inhibitory neurons are in red. Arrowheads indicate activated inhibitory neurons and arrows indicate activated excitatory neurons. Scale bars: 40 µm. Right: quantification of inhibitory (I) and excitatory (E) c-fos expressing neurons in LIV_Medial_-V_Medial_. **(B)** Cell-attached voltage clamp recording from putative DDH inhibitory (GAD67+) and excitatory (GAD67-) neurons following optic stimulation of sensory input, showing 10 consecutive traces of optically evoked action potential (oAP) discharge (left) and percentage of inhibitory (I) and excitatory (E) neurons with sensory evoked AP discharge (right). **(C)** Cell-attached voltage clamp recording showing spontaneous AP discharge in inhibitory (pink) and excitatory (black) neurons. **(D)** Percentage of neurons exhibiting spontaneous AP discharge. **(E)** Quantification of sAP frequency. **(F)** Quantification of sAP coefficient of variation (CV). **(G)** Whole-cell current clamp recording from inhibitory (pink) and excitatory (black) DDH neurons in response to depolarizing current injection. **(H)** Quantification of latency to threshold AP. **(I)** Quantification of AP number. **(J)** Quantification of AP instantaneous frequency. **(K)** Threshold AP from inhibitory (pink) and excitatory (black) neurons elicited by depolarizing current injection (left) and plot of the first derivatives as a function of time (inset), and phase plot (right). **(L)** Quantification of AP threshold. **(M)** Quantification of AP rise time. **(N)** Quantification of AP decay time. **(O)** Quantification of AP half-width. For further details on genetic crosses and statistical tests see Methods.

To determine why inhibitory interneurons are preferential recruited during walking, we performed an electrophysiological characterization of randomly sampled inhibitory (vGAT^+^) and excitatory (vGluT2^+^) neurons using the *vGAT^iresCre^* or *vGluT2^iresCre^;R26^LSL-TdTomato^ (Ai14)* mouse lines. Cell-attached recordings showed that inhibitory neurons are more likely to exhibit spontaneous action potential (sAP) discharge than excitatory neurons (**Figure 2C, D**). Of the spontaneously active neurons, inhibitory neurons showed higher sAP frequency with a lower coefficient of variation (**Figure 2E, F**), indicating that inhibitory neurons are more active than their excitatory counterparts and are a likely source of sustained inhibition. To further examine the intrinsic excitability of inhibitory interneurons, we defined their responsiveness to depolarizing current injection. Using previously established AP discharge phenotypes^44^, we found that both excitatory and inhibitory neurons within the deep dorsal horn are most likely to exhibit tonic firing (TF) AP discharge patterns (**Figure 2G**). TF INs could be further classified into two subcategories: TF-non-accommodating (TF_Na_) and TF accommodating (TF_Ac_), with excitatory neurons more likely to exhibit TF_Ac_ and DF discharge patterns than inhibitory neurons. (**Figure S1C-E**). These discharge patterns are consistent with a lower latency to the first AP, an increased number of APs, and increased instantaneous AP frequency in inhibitory interneurons (**Figure 2H-J**). We also found significant differences in AP threshold and kinetics that promote increased excitability in inhibitory neurons (**Figure 2K-O, Figure S1F-H**). Finally, inhibitory neurons displayed a more depolarized resting membrane potential (RMP) than excitatory neurons, but no differences in other passive membrane properties (**Figure S1I-K**). Together, the intrinsic properties of inhibitory neurons in the deep dorsal horn are consistent with a role in the faithful encoding of stimulus intensity and duration.

These electrophysiological data indicate that inhibitory and excitatory neurons within the deep dorsal horn process sensory information differently. Inhibitory neurons exhibited more excitable intrinsic properties, which may contribute to their increased responsiveness to sensory input and their preferential recruitment during locomotor activity. These findings suggest that inhibitory interneurons play a particularly important role in modulating sensorimotor circuits.

### dPVs are a novel population of inhibitory interneurons confined to the medial deep dorsal horn and act as a source of highly reliable ipsilateral tonic inhibition

To further investigate inhibitory interneurons within the medial deep dorsal horn (mDDH), we next aimed to identify a genetic marker that would specifically label this population. One well-known marker of inhibitory interneurons in the brain and spinal cord is the calcium-binding protein parvalbumin (PV)^45–47^. Using the PV^TdTomato^ mouse line, we identified a population of PV^+^ interneurons confined to the mDDH (**Figure 3A**), which we previously highlight as the region receiving convergent sensory input (**Figure 1B**). We named these cells deep dorsal horn parvalbumin-expressing interneurons (dPVs) and found that they represent 18+5.45% of laminae IV-V (data not shown). PV^+^ interneurons in the superficial dorsal horn (Lamina IIi-III) are composed of both excitatory and inhibitory neurons^48,49^. Thus, we investigated the neurochemical phenotype of dPVs using excitatory and inhibitory markers. We found that dPVs are largely inhibitory (**Figure 3B**), with the majority being purely glycinergic (**Figure 3C**). Notably, dPVs were found to be distinct from two previously characterized inhibitory interneuron populations in the deep dorsal horn, Satb2^+^ and RORβ^+^ interneurons^34,35^ (**Figure S2A**). In addition, we find that dPV are present in rats and non-human primates (**Figure S2B**), suggesting a conserved role across mammalian species.

**Figure 3.**
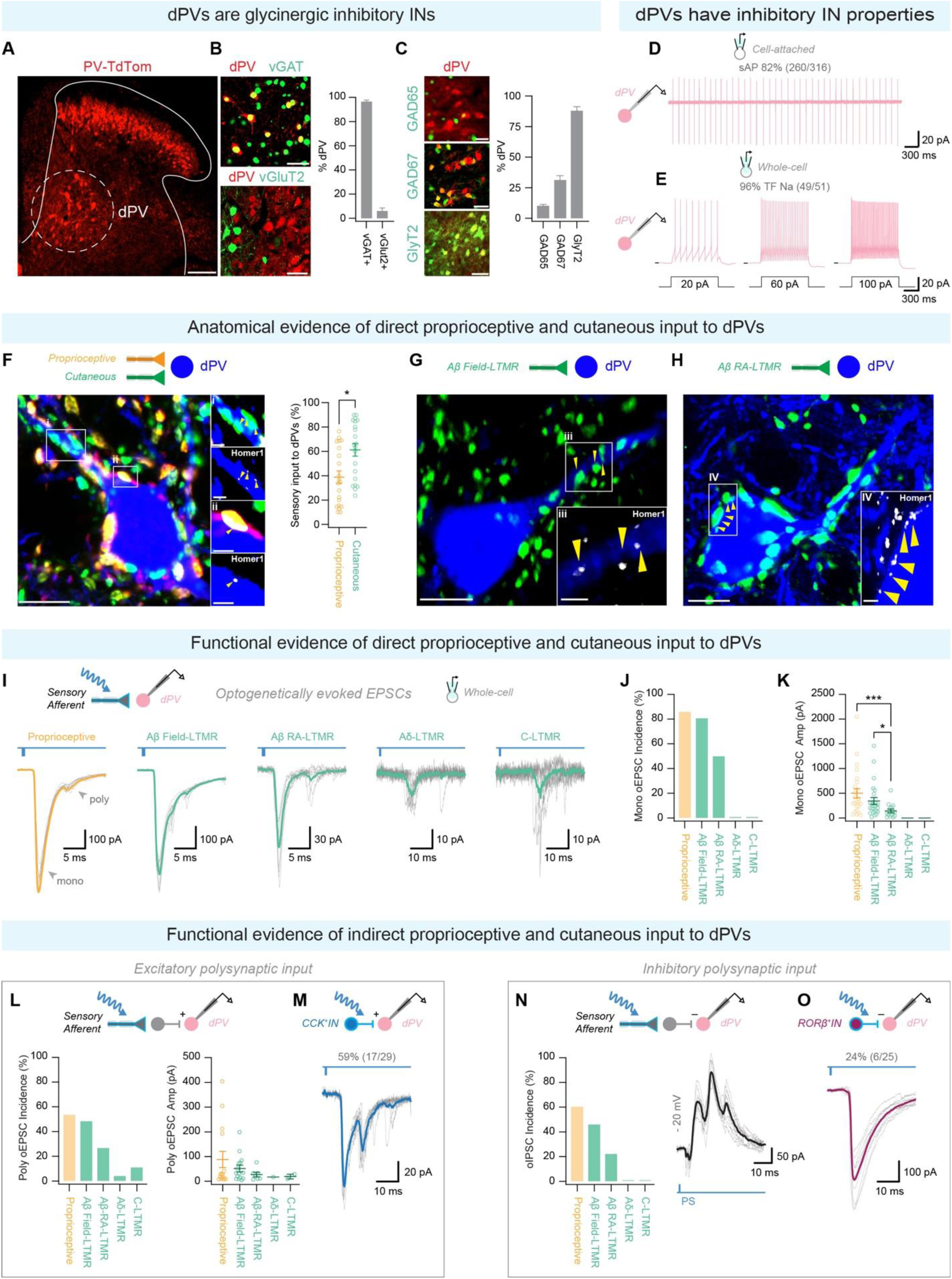
Inhibitory dPVs process convergent proprioceptive and cutaneous sensory inputs. **(A)** Transverse lumbar section of PV^TdTomato^ BAC transgenic highlighting PV-expressing interneurons in the deep dorsal horn (dPVs, circle). Scale bar: 100 µm. **(B)** Left: Images from mDDH showing colocalization of PV (red) with vGAT (top, green) and vGluT2 (bottom, green). Right: Percentage dPVs that coexpress vGAT and vGluT2+. Scale bar: 40 µm. **(C)** Left: Images from mDDH showing colocalization of PV (red) with GAD65 (top, green), GAD67 (middle, green), and GlyT2 (bottom, green). Right: Percentage dPVs that coexpress GAD65, GAD67, and GlyT2. Scale bar: 40 µm. **(D)** Cell-attached voltage clamp recording showing spontaneous AP discharge in dPVs. **(E)** Whole-cell current clamp recording from dPVs in response to depolarizing current injection. **(F)** Left: Image showing a dPV neuron (blue) in contact with cutaneous (green) and proprioceptive (yellow) puncta. Scale bar: 10µm. Inset: i) cutaneous and ii) proprioceptive puncta in contact with dPV and in apposition to the postsynaptic marker Homer1 (white, yellow arrowheads). Scale bars: 2µm. Right: Percentage of cutaneous (green) and proprioceptive (yellow) input to dPVs. **(G-H)** Images of dPV (blue) in contact with Aβ field-LTMR (G, green) and Aβ RA-LTMR input (H, green). Scale bars: 10 µm. Inset: Puncta apposition with Homer1 (white, yellow arrowheads). Scale bars: 2µm. **(I)** Whole-cell voltage clamp recordings of optically-evoked EPSC (oEPSC) from dPVs in response to photostimulation of sensory afferents (proprioceptors, Aβ Field-LTMRs, Aβ RA-LTMRs, Aδ-LTMRs, and C-LTMRs). 10 consecutive sweeps (gray) with average overlaid (bold color). **(J)** Percentage of dPV that receive monosynaptic oEPSCs following sensory afferent photostimulation. **(K)** Quantification of monosynaptic oEPSC amplitude following sensory afferent photostimulation. **(L)** Left: Percentage of dPV that receive polysynaptic oEPSCs following sensory afferent photostimulation. Right: Quantification of polysynaptic oEPSCs amplitude following sensory afferent photostimulation. **(M)** Whole-cell voltage clamp recording of oEPSCs from dPV in response to photostimulation of CCK+ interneurons. 10 consecutive sweeps (gray) with average overlaid (bold color). **(N)** Left: Percentage of dPV that receive polysynaptic oIPSCs following sensory afferent photostimulation. Right: Whole-cell voltage clamp recording (−20 mV) of oIPSC from dPV following sensory afferent photostimulation. **(O)** Whole-cell voltage clamp recording of oIPSCs from dPV in response to photostimulation of RORβ+ interneurons. 10 consecutive sweeps (gray) with average overlaid (bold color). For further details on genetic crosses and statistical tests see Methods.

To further characterize the dPV population and compare their properties to other inhibitory interneurons within the deep dorsal horn we performed a series of electrophysiological and morphological studies. Consistent with their inhibitory neurotransmitter phenotype, the majority of dPVs exhibited sAP discharge and were classified as TF_Na_ in response to depolarizing current injection (**Figure 3D, E**). These electrophysiological properties were similar to those of other inhibitory interneurons within the mDDH (**Figure S2C-I**), although dPVs exhibited higher sAP frequency and lower sAP coefficient of variation (**Figure S2E, F**). Inhibitory interneurons that receive input from proprioceptors are known to cross the midline, with dorsal horn inhibitory neurons activated by skin and group II muscle spindle afferents terminating in the contralateral intermediate zone and ventral horn^50–53^. In the developing spinal cord, Slit expression at the midline activates Robo receptor expression on commissural axons, repelling them out of the midline into the longitudinal tract on the contralateral spinal cord^54^. Thus, to determine if dPVs cross the midline, we analyzed the spinal cords of *PV^FlpO^;Robo3^iresCreER^*;*R26^LSL-FSF-Tomato^* and *PV^iresCre^*;*Robo3^iresFlpO^*;*R26^LSL-FSF-Tomato^* animals. We found that only a small portion (3.05% + 0.79) of dPVs are Robo3^+^ (**Figure S2J**), suggesting that dPV are largely ipsilaterally projecting.

In summary, our findings reveal a previously uncharacterized population of glycinergic interneurons in the mDDH - deep dorsal horn parvalbumin-expressing interneurons (dPVs). Our results suggest that dPVs act as a source of highly reliable ipsilateral inhibition across different species.

### Individual dPVs integrate convergent proprioceptive and cutaneous input from limbs

Since dPVs cell bodies are confined to the medial deep dorsal horn (**Figure 3A**), the region innervated by the distal limb, we next tested whether dPVs integrate sensory information from limbs. First, we examined whether dPV morphology changes along the rostrocaudal axis, as sensory input to the spinal cord differs between cervical, thoracic, and lumbar regions, with cervical and lumbar regions receiving input from forelimb and hindlimb respectively. We found that neuronal reconstructions of dPVs in cervical, thoracic, and lumbar segments showed heterogeneity throughout the rostrocaudal axis, with cervical and lumbar dPVs sharing common morphological features that are distinct from thoracic dPVs, most notably increased spine density, **(Figure S2K-N**). Additionally, anterograde tracers injected into the hindpaw revealed several direct excitatory contacts between paw-innervating sensory neurons and dPVs (**Figure S3A**). To assess the relative contribution of proprioceptive and cutaneous inputs onto dPVs’ cell bodies and dendrites, we used the same genetic strategies as in **Figure 1A** (*PV^2aCreER^*;*Advillin^FlpO^*; *R26^LSL-FSF-TdTomato^*; *R26^FSF-SynGFP^*). This approach revealed that both proprioceptors and cutaneous inputs converge onto single dPVs, with a bias toward cutaneous input (**Figure 3F**). Moreover, in line with their extensive dendritic arborizations (**Figure S2J**), cutaneous and proprioceptive input preferentially target dPVs dendrites (**Figure S3B**).

Given that dPVs are located within the deep dorsal horn, it is likely that they receive cutaneous input from low-threshold mechanoreceptors^49,55–57^ (LTMRs). To investigate this, we examined inputs from different classes of LTMR to the lumbar spinal cord^9,55,57,58^. We found that Aβ Field and Aβ Rapidly Adapting (Aβ RA) terminals form direct inputs onto dPVs, while C- and Aδ-LTMRs terminate superficially (**Figure 3G, H, Figure S3C**). This finding is in line with Aβ-LTMRs synapsing deeper in the dorsal horn compared to Aδ and C-LTMRs^49,55–57^. To validate these findings and examine their functional implications, we performed slice electrophysiology experiments to record the activity of dPVs in response to optogenetic stimulation of different sensory modalities (**Figure 3I**). Consistent with the histology, we found that most monosynaptic input to dPVs originates from proprioceptors, Aβ RA, and Aβ Field-LTMRs (**Figure 3J, Figure S3D-F**). The incidence and amplitude of optically evoked EPSCs (oEPSCs) were consistent with the topographical overlap between each afferent population’s central terminations and the dPV population (**Figure 3I-K, Figure S3C**).

In addition to the monosynaptic input, we observed polysynaptic input onto dPVs originating from proprioceptors, Aβ, Aδ, and C-LTMRs (**Figure 3I, L, N**). These polysynaptic inputs are likely mediated by interneurons located within the LTMR-recipient zone (LTMR-RZ, Lamina IIi-IV), which is heavily innervated by LTMRs^49^ . To further investigate these circuits, we targeted two large populations of interneurons within the LTMR-recipient zone: CCK^+^ (excitatory) and RORβ^+^ (inhibitory)^49,59,60^. We found that both CCK and RORβ^+^ interneurons provide direct input to dPVs (**Figure 3M, O**). To confirm neurotransmitter phenotype both mono- and poly oEPSC and oIPSCs were completely abolished by bath application of CNQX and Bicuculline+Strychnine, respectively (**Figure S3G, H**).

Coordinated movement requires precise sensory feedback^61,62^, which is modulated by descending cortical pathways and sensory afferents that converge on a common set of neurons^63–66^. Importantly, the majority of cortical connections with motor pathways are established through premotor networks, underscoring the integration of descending cortical command with sensory feedback^67–69^. Consistent with this concept, we show that in addition to diverse sensory input, a significant proportion of dPVs (26%) receive direct cortical input (**Figure S3I, J**).

Therefore, our findings suggest that dPVs integrate information from sensory afferents (primarily proprioceptors and Aβ LTMRs innervating glabrous skin), LTMR-recipient zone excitatory and inhibitory interneurons, and descending cortical projections. These data provide strong evidence against modality-specific labeled lines and instead suggest a diverse ensemble of inputs converge onto dPVs to inform motor output.

### dPVs are interconnected and form a diffuse inhibitory circuit throughout the deep dorsal and ventral spinal cord

Deep dorsal horn interneurons have been shown to make direct connections with motor and premotor networks of the ventral horn^23,31,70,71^. To investigate the output partners of dPV interneurons, we developed an intersectional genetic approach that would restrict recombination to the dPV interneurons of the spinal cord. We examined the overlap of dPVs with spinal cord genetic lineages and found that the Lbx1-lineage gives rise to the majority of dPVs (**Figure 4A**). An *Lbx1^Cre^*;*PV^FlpO^* intersection successfully restricts recombination to dPVs throughout the rostrocaudal axis of the spinal cord while avoiding PV^+^ neurons within the superficial dorsal horn, ventral horn, and DRG (**Figure 4B, Figure S4A-B**). Restriction of Cre/Flp recombination to dPVs is likely due to the combined action of *PV^FlpO^* eliciting more efficient recombination in dPVs (as opposed to superficial PVs) and the *Lbx1^Cre^* lineage not giving rise to the entire superficial PV+ interneuron population^72^ (**Figure S4A**).

**Figure 4.**
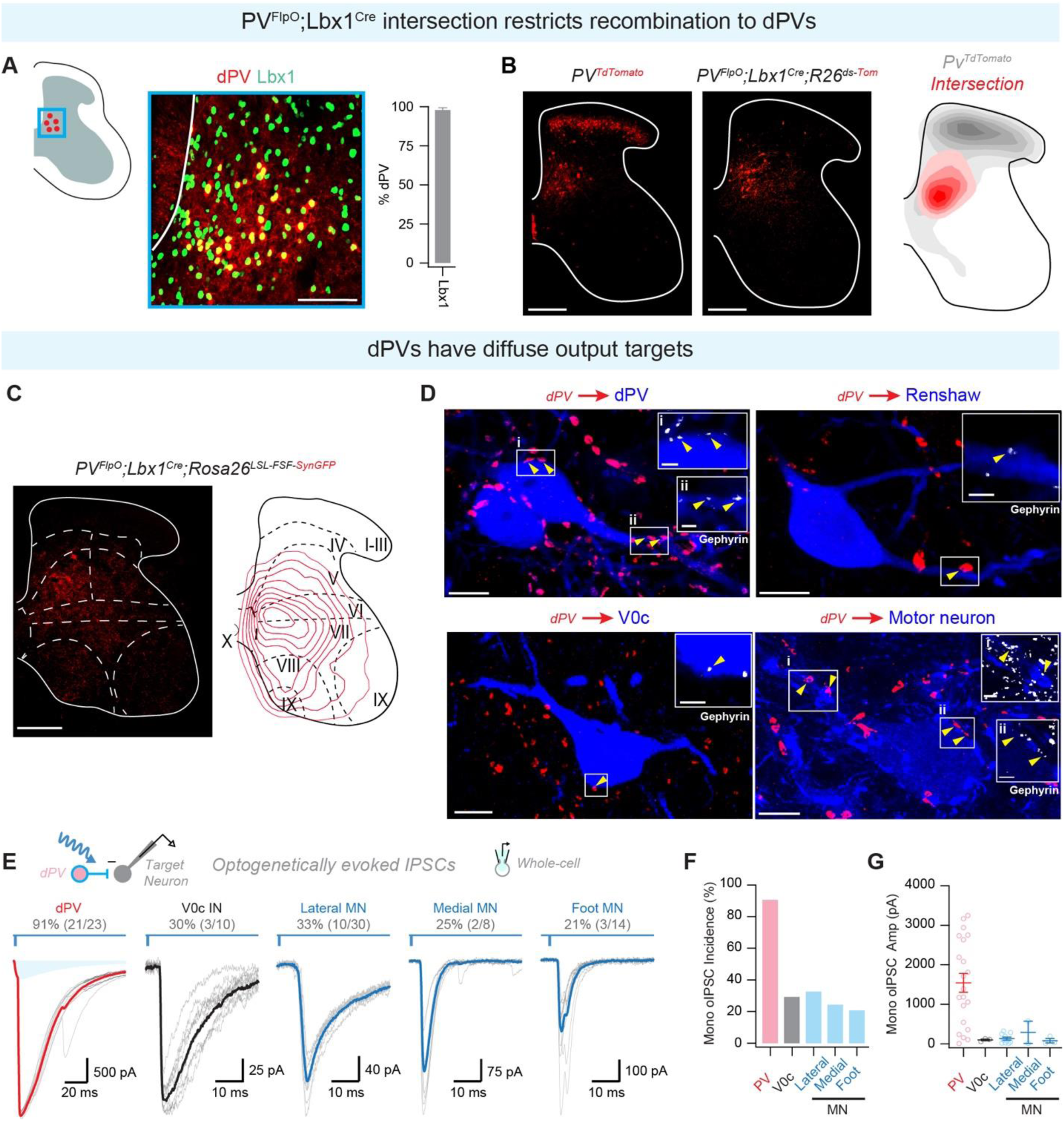
dPVs have diffuse postsynaptic targets throughout the deep dorsal and ventral horns. **(A)** Left: Colocalization of PV (red) and Lbx1 (green). Scale bar: 100 µm. Right: Percentage of dPVs that colocalize with Lbx1. **(B)** Left: Image of PV^TdTomato^ (red) expression, middle: *PvFlpO*;*Lbx1Cre* intersection (red), right: contour density plot highlighting the restriction of the *PvFlpO*;*Lbx1Cre* intersection (red) to the mDDH. Scale bars: 200 µm. **(C)** Left: Image of dPV synaptic terminals (red). Right: Density map depicting distribution of dPV axon terminals. Scale bar: 200 µm. **(D)** Images of direct contacts between dPV axon terminals (red) and other dPVs (top left, blue), renshaw cells (top right, blue), V0c interneurons (bottom left, blue), and motor neurons (bottom right, blue). Scale bar: 10 µm. Insets: Appositions to Gephyrin (white, arrowheads). Scale bar: 2 µm. **(E)** Whole-cell voltage clamp recordings of oIPSC from dPVs, V0c interneurons, lateral, medial, and foot motor neurons in response to photostimulation of dPVs. 10 consecutive sweeps (gray) with average overlaid (bold color). **(F)** Percentage of dPVs, V0c interneurons, lateral, medial, and foot motor neurons that receive monosynaptic oIPSCs following dPV photostimulation. **(G)** Quantification of monosynaptic oIPSCs amplitude following dPV photostimulation. For further details on genetic crosses and statistical tests see Methods.

We used this intersection to visualize dPVs synaptic output (*Lbx1^Cre^*;*PV^FlpO^;R26^LSL-FSF-synGFP^ (RC::FPSit)* which revealed a diffuse pattern of dPVs axon terminals through the deep dorsal, intermediate zone, and ventral horn (**Figure 4C**). To validate our findings, we also injected an AAV-Cre virus into the lumbar cord of *PV^FlpO^;R26^LSL-FSF-synGFP^* animals and observed similarly diffuse outputs (**Figure S4C**). Following unilateral injection, we did not observe any puncta on the contralateral side, further validating that the vast majority of dPVs do not cross the midline. Consistent with their neurotransmitter phenotype (**Figure 3C**), dPVs terminals as labeled by the *Lbx1^Cre^*;*PV^FlpO^;R26^LSL-FSF-synGFP^* intersection are largely glycinergic (**Figure S4D**).

Using pre- and postsynaptic histological markers we found that dPV synapses onto pre-motor neurons such as Renshaw cells (calbindin^73^), V0cs (ChAT^74^), and motor neurons (ChAT^75^) (**Figure 4D**). To further validate these findings we performed slice electrophysiology and recorded from potential postsynaptic targets while optically stimulating dPVs with channelrhodopsin in adult spinal cord slices (*Lbx1^Cre^*;*PV^FlpO^;R26^LSL-FSF-ChR^*^2^ *(Ai80)*). Consistent with our histological findings, we found monosynaptic inhibitory connections onto V0cs and motor neurons within the medial, lateral, and foot motor pools (**Figure 4E-G**). These inhibitory connections were predominantly glycinergic, in-line with a glycinergic neurotransmitter phenotype of dPVs (**Figure 3C, Figure S4E**).

Previous studies have reported synaptic coupling between PV^+^ interneurons in the superficial dorsal horn^48^, and other CNS regions^76,77^, where they can act as a precision clock for oscillatory networks^78^. This coupling is proposed to promote enhanced information coding and rhythmic gain modulation of incoming synaptic input^79,80^. Thus, we sought to determine whether PV^+^ interneurons within the deep dorsal horn share similar connections. Optical mapping of dPV outputs uncovered a surprisingly high incidence of dPV-dPV connectivity (**Figure 4E**), which was further validated by histology (**Figure 4D**). Interestingly, we found that dPV-mediated inhibition was most frequent and strongest onto other dPVs when compared to premotor and motor neurons (**Figure 4F-G**). However, the incidence and amplitude of dPV-mediated inhibition onto premotor and motor neurons was relatively equal (**Figure 4F-G**).

To investigate whether dPVs inputs onto motor neurons show any preferential bias for flexor or extensor motor neurons we employed a retrograde tracer strategy into hindlimb flexors (biceps femoris and tibialis anterior) and extensors (vastus lateralis and lateral gastrocnemius) of *Lbx1^Cre^*;*PV^FlpO^;R26^LSL-FSF-synGFP^* animals (**Figure S4F**). This approach revealed a bias toward extensor motor neurons. Lbx1-derived premotor interneurons innervating flexor and extensor motor neurons can be distinguished based on birth order. Specifically, flexor innervating neurons are born at E10.5 and extensor innervating neurons are born at E12.5^81^. Our EdU labeling of dPVs shows that 31.57+0.75% of dPVs are born at E10.5 while 43.57+0.04% are born at E12.5 (**Figure S4G)**. These findings indicate that dPVs innervate both flexors and extensors, with a bias toward extensor motor neurons.

Together, these data highlight dPVs as a source of divergent glycinergic inhibition throughout pre-motor and motor networks in the intermediate and ventral horns, with a bias towards extensor motor neurons.

### dPVs are active during walking and exhibit rhythmogenic-like activity

Based on our anatomical and functional connectivity findings, it appears that dPVs play a critical role as an integrative inhibitory node connecting multimodal sensory input and motor output pathways. To test whether dPVs are recruited during locomotion, we examined c-fos expression in dPVs after treadmill walking (**Figure 5A**). In line with our previous data showing enhanced activity in inhibitory neurons of the medial deep dorsal horn (**Figure 2A**), we found a 5.04+0.34 times increase in c-fos expressing dPVs following treadmill walking, which supports the idea that dPVs are actively recruited during locomotion (**Figure 5A**).

**Figure 5.**
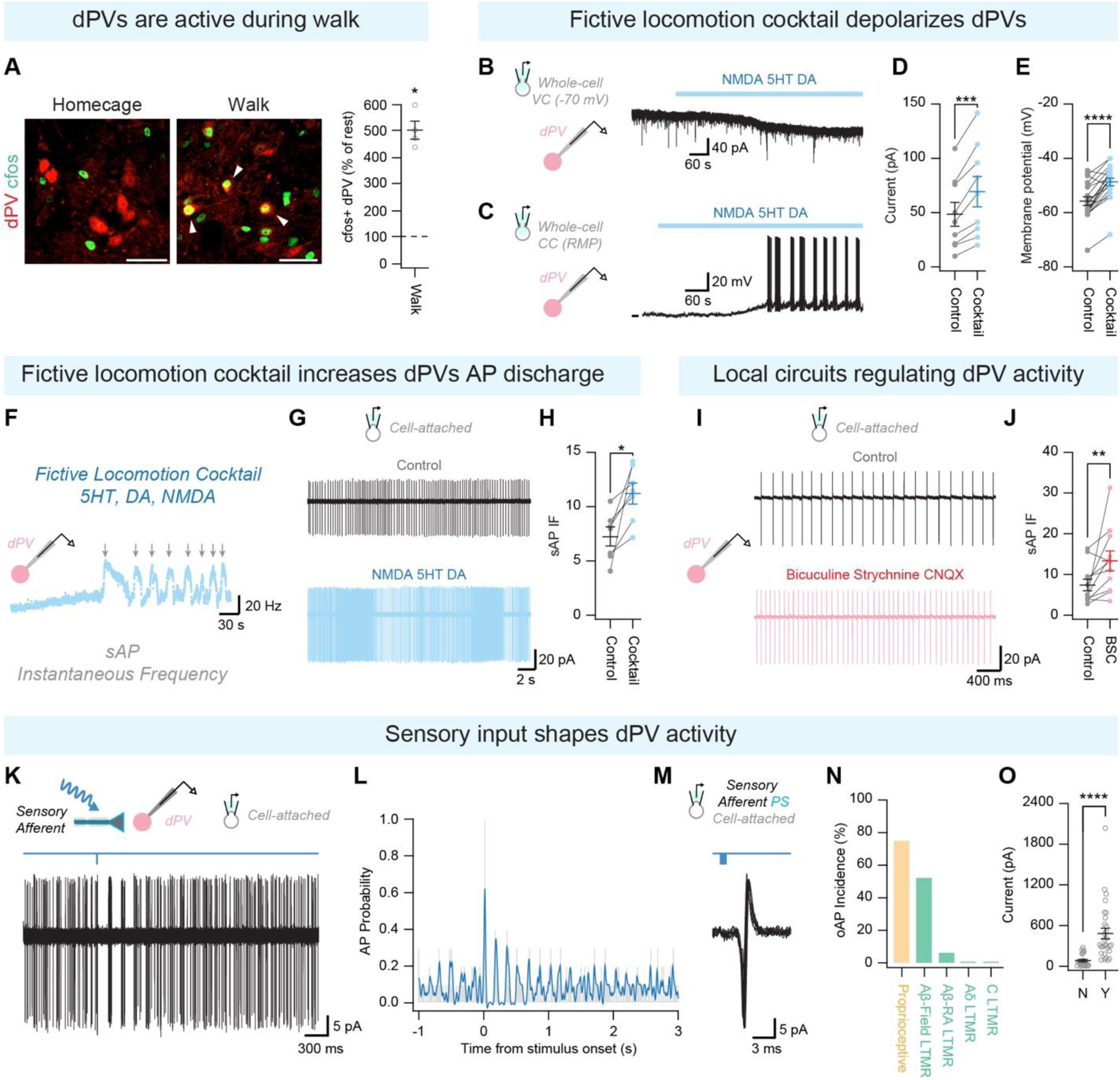
Neuromodulation and sensory input shapes dPVs activity dynamics. **(A)** Left: Images of LIV_Medial_-V_Medial_ from ‘homecage’ (left) and ‘walk’ (right) mice. c-fos labeling is in green, dPVs are in red. Arrowheads indicate activated dPVs. Scale bars: 40 µm. Right: quantification of dPV c-fos expressing neurons relative to homecage. **(B)** Whole-cell voltage clamp recording from dPV during bath application of fictive locomotion cocktail [NMDA (20 µM), 5HT (20 µM), and dopamine (50 µM)]. **(C)** Whole-cell current clamp recording from dPV during bath application of fictive locomotion cocktail. **(D)** Quantification of holding current before and after application of fictive locomotion cocktail. **(E)** Quantification of membrane potential before and after fictive locomotion cocktail. **(F)** Plot showing sAP instantaneous frequency of dPV following fictive locomotion cocktail application. **(G)** Cell-attached voltage clamp recording from dPV during bath application of fictive locomotion cocktail. **(H)** Quantification of sAP instantaneous frequency before and after fictive locomotion cocktail. **(I)** Cell-attached voltage clamp recording from dPV during bath application of bicuculline (10 µM), and strychnine (1 µM), and CNQX (10 µM). **(J)** Quantification of sAP instantaneous frequency before and after application of bicuculline, strychnine, and CNQX. **(K)** Cell-attached voltage clamp recording from dPV following photostimulation of sensory afferents. 10 consecutive sweeps overlaid. **(L)** Histogram of AP probability with an overlaid piecewise polynomial computed from p = 0.9 in MATLAB. Sensory afferent photostimulation at 0 s. **(M)** Cell-attached voltage clamp recording from dPV showing optically-evoked AP (oAP) following photostimulation of sensory afferents. 10 consecutive sweeps overlaid. **(N)** Percentage of dPV that exhibit oAPs following sensory afferent photostimulation. **(O)** Quantification of oEPSC amplitude in dPVs that do not (N) and do (Y) exhibit oAPs following sensory afferent photostimulation. For further details on genetic crosses and statistical tests see Methods.

Alternating and rhythmic outputs mimicking locomotion can be induced *in-vitro* by adding a neurotransmitter cocktail of serotonin, dopamine and NMDA^82–87^. We investigated whether dPVs exhibit conditional bursting properties that could contribute to locomotor rhythmogenesis by applying a neurotransmitter cocktail of serotonin (20 µM), dopamine (50 µM), and NMDA (20 µM) during patch-clamp recordings in adult spinal cord slices. This cocktail depolarized dPVs (**Figure 5B-E**), occasionally (3/7) evoked rhythmic bursts of AP discharge (**Figure 5F**), and increased sAP frequency (**Figure 5G, H**). Interestingly, blocking ionotropic synaptic transmission (CNQX (10 µM), bicuculline (10 µM), and strychnine (1 µM)) enhanced spontaneous AP frequency in dPVs (**Figure 5I, J**), indicating that their activity is predominantly regulated by inhibitory synaptic transmission.

Although alternating stepping can be generated in the absence of sensory input ^1^, the ability to produce smooth and coordinated movements in a dynamic environment requires the processing and integration of sensory information^61,88^. We investigated the influence of sensory input on dPV activity. Our results suggest that sensory input can ‘reset’ the timing of dPVs spontaneous AP discharge and reduce variability in output timing (**Figure 5K, L**). We found that the ability of a sensory afferent population to evoke AP discharge, and shape dPV activity, was consistent with the amplitude of their light-evoked postsynaptic currents (**Figure M-O**).

Overall, our findings suggest that dPVs are activated by overground walking and exhibit rhythmogenic like activity previously associated with locomotion. dPVs possess intrinsic properties that allow for spontaneous discharge independent of synaptic input, and they have the capacity to exhibit rhythmic bursting characteristic of left-right alternation during locomotion. Additionally, sensory input from both proprioceptors and Aβ LTMRs has the capacity to shape this activity to ensure appropriate output timing.

### dPVs modulate the timing and amplitude of cutaneous-evoked muscle activity

To investigate whether dPVs play a role in modulating motor output, we utilized a triple transgenic strategy with a dual recombinase diphtheria toxin receptor mouse line (*PV^FlpO^*;*Lbx1^Cre^*;*Tau^LSL-F-nLacZ-F-DTR^*) to specifically manipulate the activity of the dPV population *in-vivo*. We then performed intrathecal (I.T.) injection of diphtheria toxin (DTx) prepared in a hyperbaric solution to ablate dPVs^70,89^ (**Figure 6A and Figure S5A**). This strategy allowed us to generate two groups of mice: 1) dPV^abl^ (*PV^FlpO^*;*Lbx1^Cre^*;*Tau^LSL-F-nLacZ-F-DTR^* mice injected with DTx) with ablated dPVs, and 2) dPV^norm^ (*PV^FlpO^* or *Lbx1^Cre^*;*Tau^LSL-F-nLacZ-F-DTR^* injected with DTx) with non-ablated dPVs. Our approach also restricted killing of PV neurons to the deep dorsal horn (**Figure S5B**).

**Figure 6.**
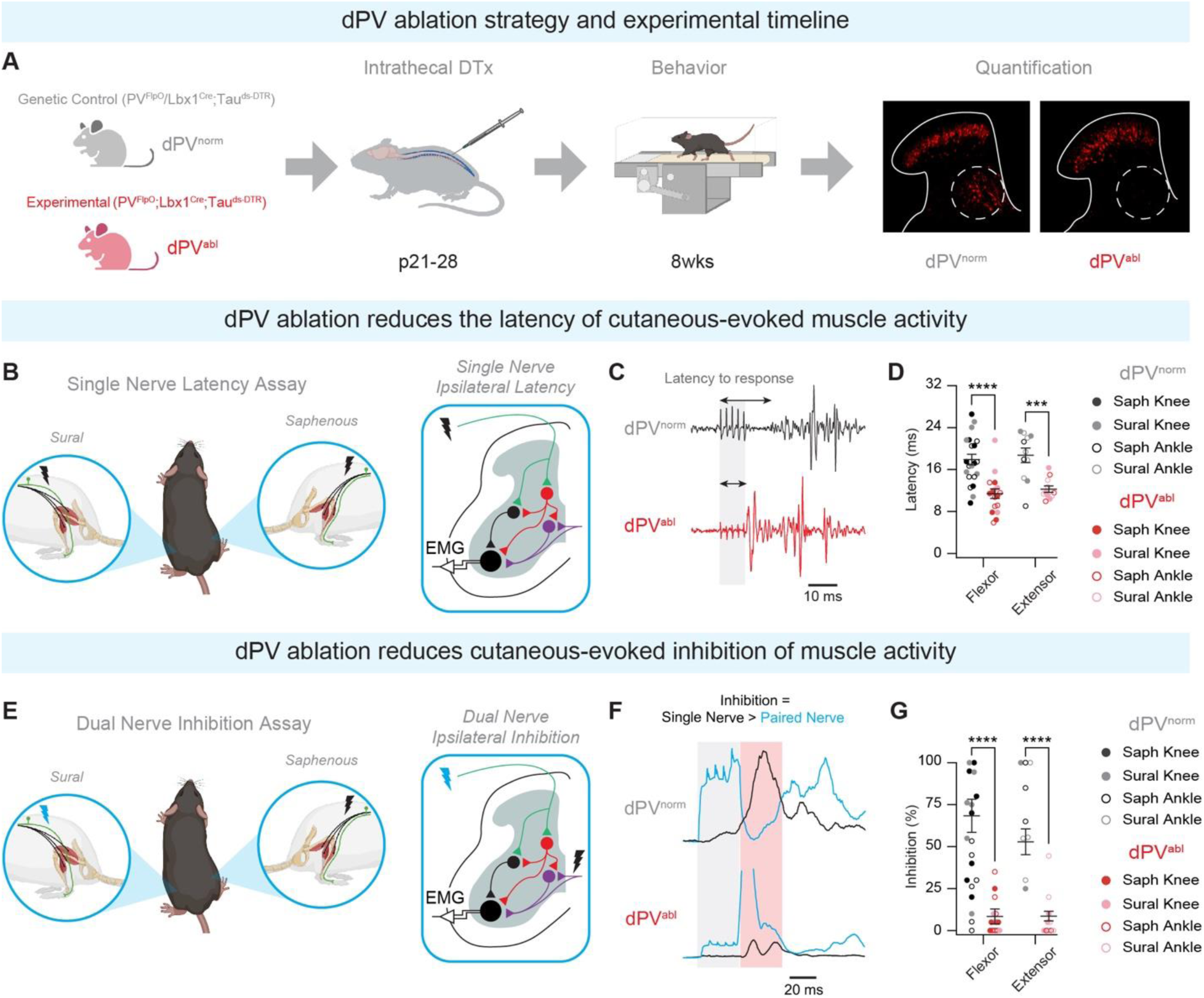
Ablation of dPVs reduces the latency of cutaneous-evoked muscle activity and the degree of cutaneous-evoked inhibition. **(A)** Experimental design for dPV ablation and behavioral studies. **(B)** Experimental setup designed to reveal the contribution of dPVs to cutaneous-evoked flexor/extensor motor activity. **(C)** EMG response to cutaneous nerve stimulation in dPVnorm (black) and dPVabl (red) animals. **(D)** Quantification of latency to EMG responses. **(E)** Experimental setup designed to reveal the contribution of dPVs to cutaneous-evoked inhibition of flexor/extensor motor activity. **(F)** EMG response to single nerve (black) and paired nerve (blue) stimulation in dPVnorm (black) and dPVabl (red) animals. Grey shading highlights the stimulus artifact, red shading highlights analysis window. **(G)** Percentage of trials exhibited cutaneous-evoked inhibition (Single nerve response > Paired nerve response). For further details on genetic crosses and statistical tests see Methods.

Sensory information plays a critical role establishing motor patterns and serves two primary functions 1) to control the timing of the transition from the stance to swing phase of locomotion, and 2) to regulate the magnitude of extensor muscle activity during the stance phase^90,91^. Our connectivity data (**Figure 3F-K** and **Figure 4D-G**) demonstrate that dPVs receive cutaneous and proprioceptive inputs and synapse onto premotor and motor networks, indicating that dPVs are ideally positioned to integrate sensory input and inform motor output. To test whether dPVs shape the ***timing*** of flexor/extensor activation in response to cutaneous input, we recorded electromyogram (EMG) activities from hindlimb flexors and extensors in response to sensory nerve stimulation in dPV^norm^ and dPV^abl^ mice (**Figure 6B**). Sural and saphenous nerves, which respectively carry cutaneous afferent fibers innervating the posterior and anterior distal hindlimb^14,92–95^, were stimulated while recording from knee and ankle flexors and extensors. Our results demonstrate that dPV ablation decreased muscle response latency in both flexors and extensors following ipsilateral stimulation of the saphenous and sural nerves (**Figure 6C-D**), suggesting that dPVs shape the timing of motor responses to cutaneous stimulation.

To test the ability of dPVs to modulate the ***magnitude*** of muscle responses to sensory input we designed a paired nerve assay rooted in the inhibitory crossed reflex pathway mediated by cutaneous feedback^94,96^. In this assay, stimulation of a contralateral cutaneous nerve evokes inhibitory pathways that suppress a motor response mediated by ipsilateral cutaneous nerve stimulation (**Figure 6E**). We started by stimulating the ipsilateral nerve and calculating the averaged expected response (black trace, **Figure 6F**). Next, both ipsi- and contralateral nerves were stimulated at varying intervals (see **Figure S5C**). The summed response (blue trace, **Figure 6F**) was compared to the expected response to reveal the magnitude of inhibition. Using this approach, we observed a decrease in the magnitute of cutaneous-induced ipsilateral inhibition to flexor and extensor muscles following dPV ablation (**Figure 6G**).

These data suggest that dPVs mediate ipsilateral cutaneous evoked-muscle inhibition to both flexors and extensors. We additionally performed the experiments to reveal the role of dPVs in mediating muscle responses to contralateral cutaneous nerve stimulation (**Figure S6**). Our results showed dPV ablation reduced the muscle response latency in both flexor and extensors (**Figure S6A**) and reduced nerve-evoked inhibition in flexors (**Figure S6B**) following contralateral cutaneous afferent stimulation. This role could be mediated by commissural dPVs, representing ∼3% of dPVs (**Figure S2J**), or by contralateral neurons, such as V0c’s targeted by dPVs (**Figure 4D-E**).

Together, these data suggest that by inhibiting the transmission of sensory information onto motor circuits, dPVs play an important role in modulating the timing and magnitude of flexor and extensor muscle activity in response to sensory input.

### dPVs do not contribute to corrective movements during skilled tasks

Corrective movements and adaptations to perturbations are integral to the execution of skilled tasks. The effective integration of sensory input, both cutaneous and proprioceptive, is crucial in facilitating this phenomenon^10,61,97–100^. Recent work suggests that interneurons and projection neurons in the deep dorsal horn and intermediate zone are also critical for corrective movements^70,101,102^. Since dPVs receive direct and indirect cutaneous/proprioceptive input (**Figure 3F-K**), we investigated whether dPVs shape corrective motor reflexes by allowing dPV^abl^ and dPV^norm^ mice to locomote across a beam with different shapes and sizes, a horizontal ladder with changing rung patterns, and a rotarod. We observed that dPV^abl^ and dPV^norm^ mice do not exhibit any major differences in performance on all three tasks **(Figure S7A-C**). Taken together, these data suggest that unlike other interneuron populations in the deep dorsal horn, dPVs do not influence corrective motor reflexes.

### dPVs ablation alters limb kinematics at the transition from stance to swing

Somatosensory feedback actively regulates the durations and transitions of locomotor phases by interacting with spinal circuits, dynamically modulating the duration and magnitude of muscle activity^12,36,103^. These motifs of circuit activation enable individual extensor and flexor muscles that control the hip, knee, and ankle joints to produce distinct and stereotypic patterns of recruitment, with marked oscillation in flexor-extensor muscle recruitment at the transition between step cycle phases (swing to stance and stance to swing)^90,104,105^. Given we observed a role for dPVs in gating the timing and magnitude of sensory-evoked muscle activity (**Figure 6**), we sought to determine if ablating dPVs affects limb kinematics in ways that are consistent with a role in processing convergent cutaneous/proprioceptive input. We subjected dPV^abl^ and dPV^norm^ mice to treadmill locomotion and used DeepLabCut^106^ for pose estimation of important landmarks along the right hindlimb throughout the step cycle (**Figure 7A-B**). Expectedly, we noticed the most profound differences in joint angles at the transition from the stance to swing phase. To quantitate hip, knee, and ankle joint angles at this transition of the step cycle, we defined this instance as the posterior extreme position (PEP) of the toe. For comparison, we also defined the anterior extreme position (AEP), occurring at the transition from the swing to stance phase. Our results demonstrate that dPV^abl^ mice exhibit significantly more flexed knee and ankle joints at the PEP, with no observed changes in joint angles at the AEP (**Figure 7C-E**). Altered kinematics across multiple joint angles suggests dPVs influence is not restricted to a specific motor pool and is consistent with our finding that dPVs make divergent outputs onto both pre-motor networks and multiple motor pools within the ventral horn (**Figure 4D-E**). Changes in limb kinematics specific to PEP may indicate that dPVs play a role in producing fluid locomotion by securing smooth switching from stance to swing phases, which may act to regulate the rate of stride propagation.

**Figure 7.**
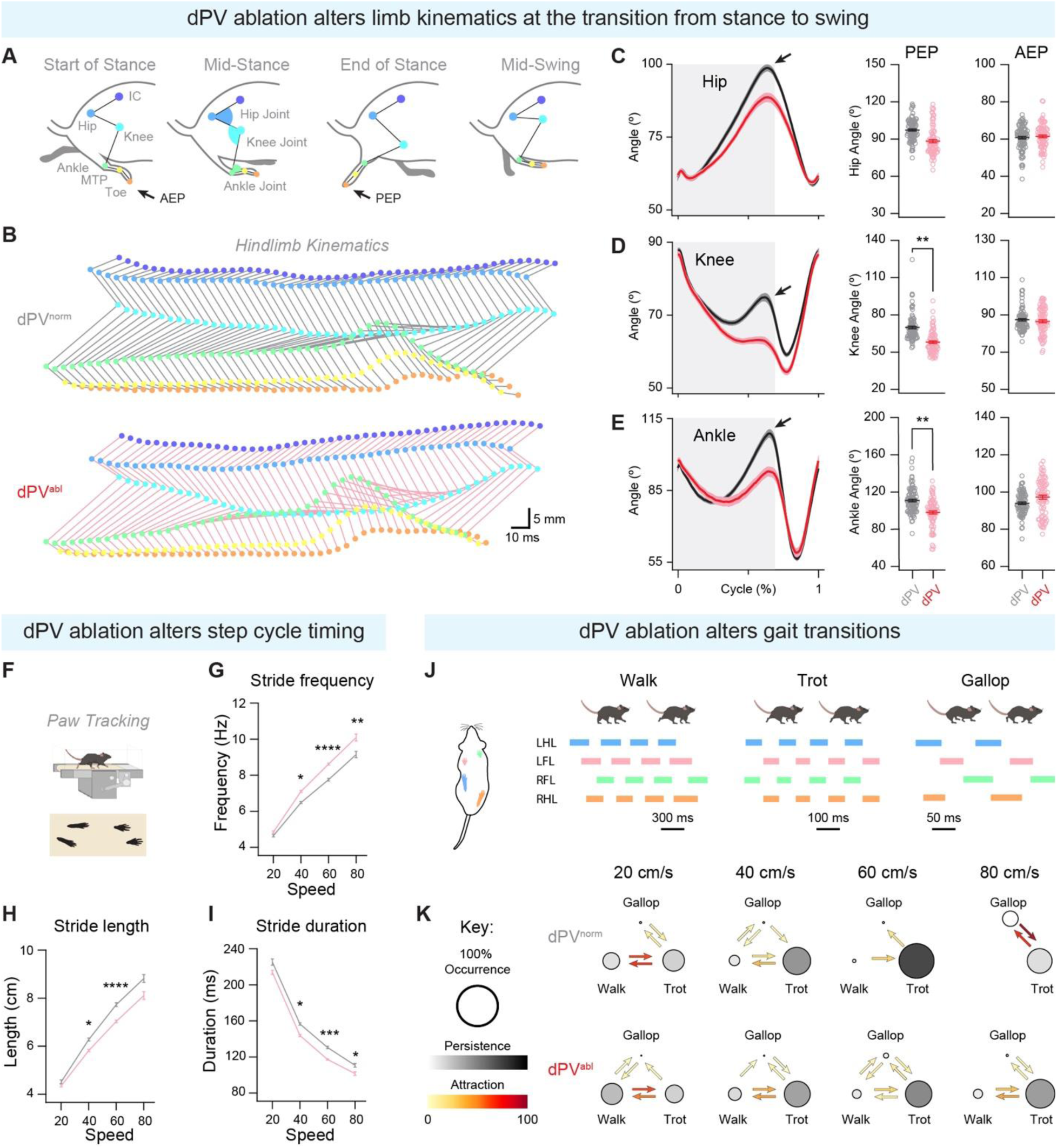
Reduced sensory-evoked inhibition following dPV ablation alters limb kinematics and step cycle timing. **(A)** Illustrative schematic of stride phases with labeled hindlimb landmarks (Iliac crest (IC), hip, knee, ankle, first metatarsal phalangeal (MTP), and toe) that were used to quantitate the hip, knee, and ankle joints. **(B)** Example stick diagrams of a step performed by dPV^norm^ (top, gray) and dPV^abl^ (bottom, pink) demonstrates changed joint angles and limb kinematics. **(C-E)** Left: Estimation of **(C)** hip, **(D)** knee, and **(E)** ankle joint angles during normalized step cycle (0 is the start of stance and 1 is the end of the swing phase). Arrows denote the posterior extreme position (PEP), gray shading denotes the stance phase. Right: Quantification of hip, knee, and ankle angles at PEP (left) and anterior extreme position (AEP, right) show that dPV^abl^ mice exhibit significantly more flexed knee and ankle angles during the transition from stance to swing (PEP). **(F)** Paw tracking used to quantify step cycle and gait during locomotion. **(G-I)** Quantification of **(G)** stride frequency; **(H)** stride length, and; **(I)** stride duration across four speeds: 20, 40, 60, and 80 cm/s). **(J)** Diagram depicting walk, trot, and gallop gaits during treadmill locomotion. **(K)** Gait transitions at four locomotor speeds of dPV^norm^ (top) and dPV^abl^ (bottom) mice show that dPV^abl^ animals bias a walking gait, especially at higher locomotor speeds (i.e., 60 and 80 cm/s). Circle size, circle color, and arrow color represent gait occurrence, persistence, and attractiveness, respectively (see Experimental procedures for definitions). For further details on genetic crosses and statistical tests see Methods.

### At increasing speeds, dPV ablation disrupts the timing of step cycle and gait transitions

In-turn, we assessed the rate of stride propagation in dPV^abl^ and dPV^norm^ mice during treadmill locomotion at a range of speeds with a high-speed camera located underneath a transparent belt (**Figure 7F**). We show that dPV^abl^ mice exhibit increased stride frequency, reduced stride length, and shorter stride duration compared to dPV^norm^ mice (**Figure 7G-I**), particularly at increased treadmill speeds (40 - 80 cm/s). Therefore, dPV^abl^ mice adopt a shorter, more frequent stride at higher speeds of locomotion, indicating an involvement in step cycle timing. Given these alterations in stride dynamics, we proceeded to investigate whether dPV ablation affects gait patterns.

To maintain stability at increasing speed, mice typically transition from walking to trotting, galloping, and bounding, adjusting their inter-limb and intra-limb stepping patterns^107^ (**Figure 7J**). To assess the impact of dPV ablation on gait transitions, we classified each step cycle as one of these four main gaits. While dPV^norm^ fully transitioned to trotting at 60 cm/s and alternated between trotting and galloping at 80 cm/s, dPV^abl^ retained the walking gait at higher speeds and refrained from galloping at 80 cm/s (**Figure 7K**). Gait transitions require appropriate adjustment in limb coordination^108^. To determine whether dPV^abl^ mice display altered limb coordination, we calculated phase coupling between two sets of limbs in dPV^abl^ and dPV^norm^ mice (**Figure S7D**). Our analysis revealed a shift in ipsilateral coupling in dPV^abl^ mice, toward an out-of-phase coordination pattern.

Taken together, our findings suggest that dPVs play a role in shaping limb kinematics and regulating step cycle timing, facilitating smooth transitions between different phases of locomotion. Further, the alterations in limb dynamics observed in dPV^abl^ mice promote the maintenance of a walking gait even at higher speeds of locomotion.

### dPV ablation disrupts movement transitions during naturalistic motor behaviors

Studies of intermediate spinal cord^31,109^ and ventral horn glycinergic networks^110,111^ propose that spinal cord interneurons with properties like dPVs influence unique sub-second movements (aka. ‘switching’) required for the structure of naturalistic motor behaviors (i.e., turning, rearing, etc.). Thus, while the activity in dPVs during forced locomotion is important for setting the timing and amplitude of flexor/extensor activities, their divergent action onto motor and premotor networks may facilitate movement switching during unconstrained behavior, contributing to the generation of variable and adaptable motor behaviors. Thus, deep dorsal horn glycinergic networks, including dPVs, by virtue of their multimodal sensory input (touch and proprioceptive, **Figure 3E-J**) and divergent output pathways (motor and premotor, **Figure 4D-G**) may serve to titrate excitatory drive onto ventral horn motor networks to sculpt movement in more nuanced ways.

To test this hypothesis, we used machine learning-assisted three-dimensional video analysis (Motion Sequencing, MoSeq^112–115)^. MoSeq combines depth imaging with unsupervised computational algorithms to fractionate mouse behavior into brief (∼350 ms) behavioral “syllables” such as turning, rearing, and crouching (**Figure 8A**). These syllables are stereotyped, reused, and concatenated into larger complex behaviors by specified “transition probabilities”. This approach can therefore phenotype mice by screening for deleterious or neomorphic behaviors that emerge because of spinal cord circuit dysfunction^115^, such as when dPVs are ablated. Importantly, MoSeq captures all behavioral features, including those that may be overlooked when focusing on traditional outcome measures like joint angles or speed.

**Figure 8.**
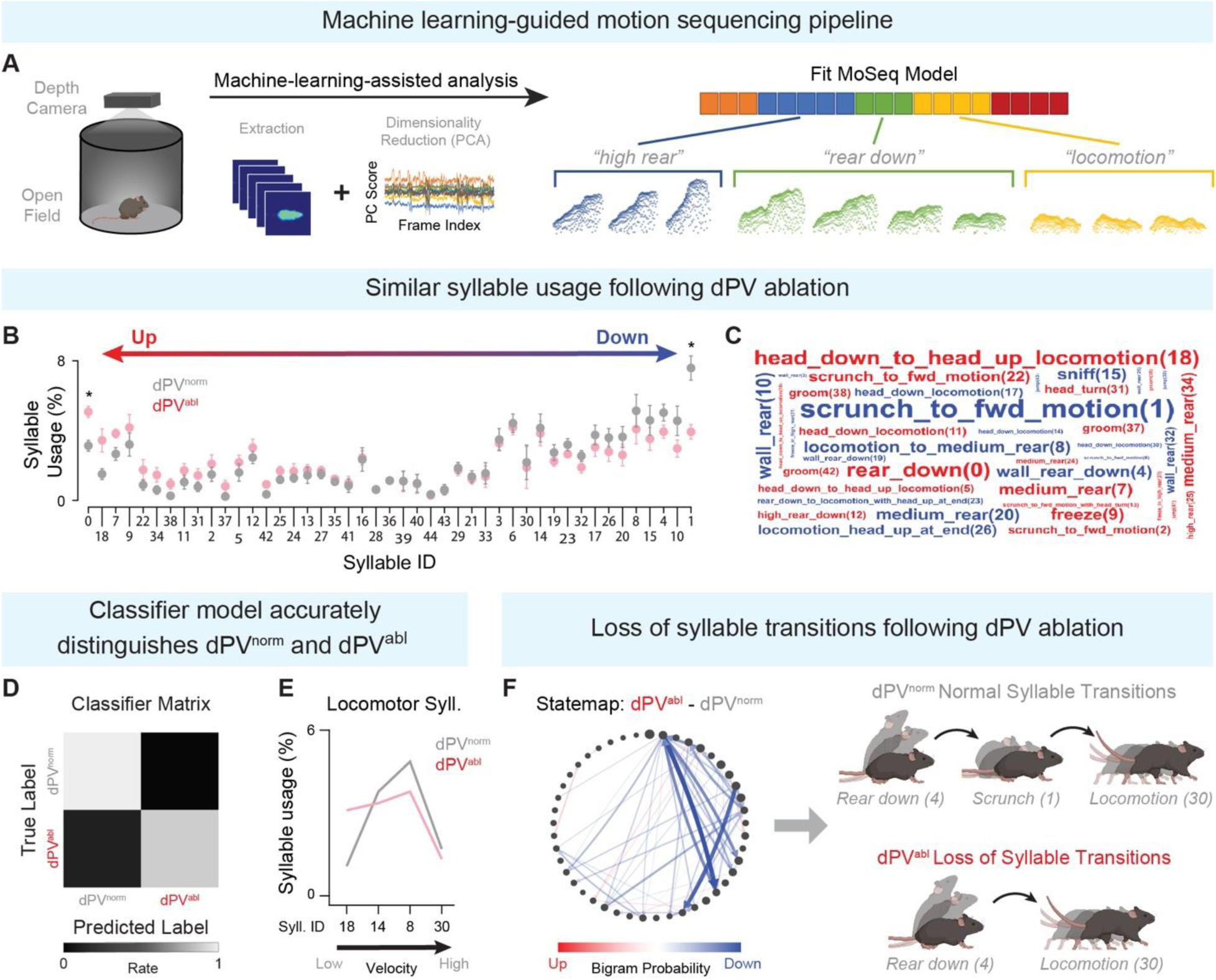
dPVs enable smooth movement transitions during naturalistic behavior. **(A)** Experimental paradigm of the machine learning-guided motion sequencing (MoSeq) pipeline: recording is performed using 3D imaging in an open arena (left) and unsupervised computational modeling fractionates complex motor behaviors into stereotyped, recurring sub-second modules (behavioral “syllables”) such as rearing, coming down from a rear (rear down), and locomotion. **(B)** Syllable usage in dPV^norm^ (grey) and dPV^abl^ (red) animals ordered by relative differential usages from the most upregulated (left, red arrow) to the most downregulated syllables on the (right, blue arrow) following dPV ablation. **(C)** Word cloud representation of syllable usages depicted by ethological descriptor and corresponding syllable number in parentheses. dPV ablated up- and downregulated syllables are colored in red and blue, respectively, and sized by differential syllable usage between groups. **(D)** Normalized classification matrix demonstrating the linear discriminant analysis (LDA) classifier performance in distinguishing between dPV^norm^ and dPV^abl^ mice. A higher rate indicates improved discrimination. **(E-F)** Logistic regression analysis unearthed a series of distinct syllables and transitions between syllables that carry a large weight in discriminating between dPV^norm^ and dPV^abl^ mice (see **Figure S8**, and **Tables S1 and S2** for full characterizations of syllables and transitions). **(E)** Four locomotor syllables were identified as demonstrating large contribution weight to, and high probability of inclusion in, the classifier model: syllable ID 18 (head down to head up locomotion, ∼9 cm/s), syllable ID 14 (head down locomotion, ∼10.5 cm/s), syllable ID 8 (locomotion to medium rear, ∼12 cm/s), and syllable ID 30 (head down locomotion, ∼21 cm/s). dPV^abl^ mice bias syllable 18 (slow) and dPV^norm^ mice bias syllable 8 (fast). **(F)** Left: Behavioral statemap illustrates syllables as nodes and probability of transitioning between syllables as arrows. Using bigram probabilities, the directionality and color of the arrows (upregulated transition probabilities are red, downregulated are blue) are weighted by the difference between statemaps of dPV^norm^ and dPV^abl^ mice. Right: Example syllable transitions demonstrate that dPV^abl^ mice (bottom) exhibit less complexity in motor behavior as compared to dPV^norm^ mice (top), preventing these animals from smoothly transitioning from one movement to another. For further details on genetic crosses and statistical tests see Methods.

The MoSeq pipeline involves several key features for high-throughput data analysis: (1) A depth camera is positioned above an open arena where the mouse can freely behave, assigning three dimensions (x, y, z - with z being the height of the mouse off the ground of the arena) to each “mouse” pixel. (2) A hyperparameter is selected to identify pervasive behavioral discontinuities in the recorded dataset at the sub-second to second timescale. This hyperparameter accurately identifies syllable boundaries and transitions from one syllable to the next. (3) Additional classification and statistical algorithms, such as multinomial logistic regression models, can be employed to categorically estimate the distribution of single syllables and transition probabilities to the group (experimental, control) that is exhibiting those behavioral motifs more frequently or stereotypically.

Here, we use MoSeq to test the specific hypothesis that the divergent action of dPVs onto motor and premotor networks facilitate movement switching, leading to the emergence of variable and adaptable behaviors. As such, using MoSeq we would not expect that individual module usage be affected, but rather how the animal transitions between the modules, and/or the variability in which these modules are emitted. Our analysis demonstrated that the overall usage of individual modules remained largely unchanged, with significant changes to the usage of only two modules out of 44 (**Figure 8B-C**). Despite this, both linear discriminant analysis (LDA) and a multinomial logistic regression classifier accurately distinguished between dPV^norm^ and dPV^abl^ mice (**Figure 8D**), indicating discernible differences in their behavioral patterns. Notably, we identified four locomotor syllables that contributed significantly to the classifier model (**Figure 8E**), displaying high inclusion probability and weight. Within these syllables, dPV^abl^ mice exhibited a preference for lower velocity movements compared to higher velocity movements (See **Table S1** for all syllables with large weights). This is in-line with our previous finding that dPV^abl^ mice prefer walking gaits (**Figure 7 G-K**), and we therefore propose that during naturalistic behavior dPV^abl^ mice will select speeds that accommodate a walking gait.

Additional analysis using the multinomial logistic regression classifier revealed transition probabilities (bigram probabilities) that have large contribution weight in discriminating between dPV^norm^ and dPV^abl^ groups (**Table S2**). We observed that following dPV ablation, transition probabilities are largely downregulated (**Figure 8F**), indicating a decrease in the complexity of motor behavior exhibited by dPV^abl^ mice. In the example shown in **Figure 8F**, dPV^norm^ mice transition from coming down from a rear (Syllable ID 4) to a scrunch and propel forward (Syllable ID 1), and finally to locomote (Syllable ID 30). In contrast, dPV^abl^ mice display a similar rear down to locomotion pattern, but without an intermediary syllable, suggesting a role for dPVs in defining particular features of complex motor behavior. These results support the hypothesis that dPVs are involved at the junction between distinct behavioral motifs, emphasizing their role in facilitating smooth transitions from one motor movement to the next.

Together these data suggest that dPVs play a role in enabling movement switching during naturalistic behaviors. These findings align with our kinematic and gait analysis, reinforcing the significance of dPVs as key mediators in ensuring seamless transitions between distinct movements. By modulating transition probabilities and complexity of motor behavior, dPVs may contribute to the generation of variable and adaptable behaviors, ultimately enhancing the versatility and adaptability of the motor repertoire.

## DISCUSSION

The medial deep dorsal horn (mDDH) of the spinal cord is ideally placed to integrate incoming sensory input and inform motor output. Several studies have identified this region as an important hub for sensorimotor control ^23–31^. However, the specific mechanisms by which sensory input is integrated to inform motor output have been unclear. In this study, we demonstrate that inhibitory interneurons within the mDDH receive convergent sensory input (cutaneous and proprioceptive) and exert tonic inhibitory control onto premotor and motor networks to regulate the timing and magnitude of motor neuron activity. By ablating dPVs, we observed a reduction in cutaneous evoked inhibition and a reduction in latency of cutaneous evoked motor neuron activity. These circuit changes result in altered step-cycle dynamics, hindlimb hyperflexion at the transition from stance to swing during locomotion, and a reduction in transition probabilities between distinct movements during naturalistic behavior.

### The medial deep dorsal horn is a node for multimodal sensorimotor processing

Integrating information from multiple sensory modalities is crucial for appropriate motor output. Convergence is a mechanism for the emergence of complex processing that represents more aspects of the sensory experience. For instance, the convergence of visual inputs within the cortex allows for movement and orientation selectivity^116–118^ and convergence within the inferior colliculus underlies our ability to attribute all aspects of sound to one entity, orient to the sound, and distinguish it from self-generated sounds^119^. In the spinal cord, proprioceptive and cutaneous inputs modulate the magnitude of muscle activity and the timing of the step cycle during locomotion^120–130^. However, it remains unclear whether these two modalities remain in parallel pathways or converge to adjust motor output. In support of an integrative model, previous studies have demonstrated facilitation or depression of proprioceptive reflexes when coupled with cutaneous afferent stimulation^17,18,131^. In this study, we demonstrate that the mDDH integrates convergent proprioceptive and cutaneous input (**Fig 1B-C & Fig 3F-K**), with individual excitatory and inhibitory neurons within the mDDH receiving convergent proprioceptive and cutaneous afferent input.

We propose that sensory convergence within the mDDH serves two primary functions: (1) to improve the efficiency and accuracy of locomotion modulation by sensory inputs, and (2) to enable the system to adapt to different contexts, with output signals dependent on the mix of sensory information present at any given time. Supporting this idea, we observed that sensory afferent stimulation ‘resets’ the timing of spontaneous action potential discharge in dPVs to increase the uniformity of output timing (**Fig 5K-L**). This response could play a role in the ‘reset’ in fictive locomotor rhythm that is observed following stimulation of proprioceptive/cutaneous afferents^126,127,132^. It’s important to note the limitations to our manipulations of sensory afferent activity; made *in-vitro,* without background activity or neuromodulation that can shape neural activity, and resulting in artificially synchronous input that is unlikely to occur *in-vivo*. Despite these limitations, our data suggest that sensory convergence increases the diversity of mechanisms by which sensory information can influence mDDH networks in a context-dependent manner, depending on the combination of inputs active at any given time (e.g., during stance or swing). Our findings highlight the mDDH as a key node for multimodal sensorimotor processing and provide new insights into the integration of sensory inputs for motor output control.

### Sensorimotor coding by inhibitory neurons within the mDDH

The mDDH contains a large population of premotor interneurons^23–28^ that participate in cutaneous withdrawal reflexes^29,30^ and coordinate motor response across multiple muscle groups^31^. Our study reveals that dPVs within the mDDH exhibit highly divergent outputs that spread throughout the deep dorsal horn, intermediate zone, and ventral horn to form synaptic contacts with premotor circuits and motoneurons (**Fig 4C-E**). This diffuse inhibition is consistent with previous anatomical studies showing that axonal projections of inhibitory interneurons relaying proprioceptive input are more extensive than their excitatory counterparts^53^. Despite the widespread output dPVs’, retrograde labeling suggests preferential targeting of extensor motor neurons supporting a medio-lateral spatial segregation of Lbx1-derived extensor and flexor premotor interneurons^81^ and provides further evidence for a disynaptic link between proprioceptors and extensor motor neurons^132–135^. dPVs postsynaptic circuitry suggests widespread inhibitory control over a variety of cell types involved in sensorimotor processing. Under this arrangement, inhibitory interneurons within the mDDH act as a source of broad inhibitory control, regulating the gain of network activity.

Despite a likely role for the mDDH in motor control, little is known of how the properties of excitatory and inhibitory neurons within this region encode sensory information to inform motor output. Similar to previous reports^136,137^, we observed that DDH neurons exhibit tonic firing, which produces reliable signal transmission^137,138^. They also provide a constant inhibitory/excitatory tone that can be differentially tuned by sensory input. Furthermore, we demonstrate the rhythmogenic capacity of dPV inhibitory neurons. Together, this arrangement supports a push-pull^139,140^ mechanism for gain control, where tonic inhibitory and excitatory inputs increase response gain to phasic sensory input^139^.

Similar to the superficial dorsal horn^44^, we show that inhibitory neurons within the DDH are more excitable than excitatory neurons, promoting their preferential recruitment during locomotion (**Fig 2**). These properties would promote earlier recruitment of inhibitory neurons, which means inhibitory regulation is in place before excitation provided by local interneurons transmits the sensory signal. Similarly, within the LII-III of the dorsal horn Aβ-evoked inhibition arrives roughly 40 ms before Aβ evoked EPSPs reach firing threshold^141^. This arrangement suggests inhibition provided by the mDDH must be overcome by excitatory pathways to achieve polysynaptic signal transduction, a model that would effectively reduce network excitability and promote selective muscle recruitment during locomotion.

### Regulation of motor output by mDDH inhibitory neurons

In shaping motor activity, inhibitory interneurons play a crucial role by regulating the transmission of sensory information and influencing the pattern and rhythm of motor neurons. The modulation of motor output through inhibition is achieved through several mechanisms, including presynaptic inhibition of sensory afferents^35,62,142^, postsynaptic inhibition of spinal neurons interposed in sensorimotor pathways^34,143,144^, as well as direct postsynaptic inhibition of motor neurons^31,34,145–149^. This study focuses on a population of ipsilaterally projecting inhibitory interneurons, dPVs, and demonstrates that their ablation leads to a reduction in the latency and inhibition of cutaneous-evoked motor activity (**Fig 6C, F**). These findings suggest that dPV-mediated inhibition plays an important role in gating the flow of sensory information, ensuring the appropriate timing and magnitude of motor unit recruitment.

Previous research investigated the role of two other inhibitory interneuron populations within the DDH. The removal of RORβ interneuron-mediated presynaptic inhibition of proprioceptive afferents results in a ‘duck-gait’ phenotype characterized by hip hyperflexion^35^, while disruption of Satb2 interneuron-mediated postsynaptic inhibition leads to hyperflexion of the ankle beginning at swing onset^34^. These studies highlight the importance of inhibitory gating of sensory input in promoting appropriate limb flexion. Building upon these findings, we show that ablation of dPVs, which reduces inhibitory gating of sensory input, causes hyperflexion of the knee and ankle during the onset of the swing phase (phase-dependent). We also demonstrate altered gait dynamics are speed dependent (state-dependent). Our data suggest that dPVs and inhibitory interneurons within the deep dorsal horn play a state- and phase-dependent role in modulating locomotor behavior.

The specific mechanisms by which loss of divergent inhibition following dPV ablation leads to a hyperflexion phenotype remains unclear. It is plausible that the deficits observed following dPV manipulation result from the loss of direct motor neuron inhibition as well as indirect inhibition of premotor circuits. Our findings align with previous literature highlighting the crucial role sensory input in regulating limb flexion timing and phase transitions from swing to stance^11,126,128,150^. It is possible that the hindlimb hyperflexion observed is due to defects in how sensory inputs control the timing of limb flexion. Collectively, these findings suggest that reduced sensory gating in sensorimotor circuits, whether by presynaptic or postsynaptic mechanisms, disrupts the balance between flexors and extensors to impair phase transition timing, ultimately promoting flexor activation during the transition from stance to swing.

Natural behaviors, such as rearing, turning, and crouching, are likely composed of simple movement modules assembled in sequence^151,152^. This modular organization, mirroring the proposed modular organization of the spinal cord^7,153–155^, is evident in sensorimotor circuits across various species: scratch in turtles^156^, kicking in frogs^152^, nociceptive withdrawal reflex of rodents, cats and humans^157^. Deep dorsal horn premotor networks represent an ideal system to explore this modular organization. First, this region represents the largest population of premotor networks^31^. Second, this region serves as a convergence point for inputs important for motor control: proprioceptive, cutaneous, corticospinal, rubrospinal input^29,30,158–162^. Third, extensive connectivity and functional features support DDH interneuron’s role in coordinating different muscles to shape complex movement patterns^31^. Lastly, activation of the deep dorsal horn is sufficient to induce complex motor behaviors^23–28,31,163–165^. Thus, deep dorsal horn premotor networks are uniquely poised to shape complex motor behavior by dictating distinct sub-second movements based on their unique sensory input and motor neuron output connectivity patterns^31,163^. Recent work suggests that spinal cord glycinergic pre-motor networks form part of the locomotor ‘switch’, enabling the smooth transition from one behavior to another^111^. Combining 3-D depth imaging with artificial intelligence enabled us to dissect naturalistic behaviors into sub-second behavioral syllables (Motion Sequencing: MoSeq, **Figure 8A**)^115^. In-line with a role for glycinergic inhibition in behavioral ‘switching’, we demonstrate a decrease in motor behavioral complexity, as indicated by a loss of syllable transitions following dPV ablation (**Figure 8F**). These findings reveal a new role for spinal inhibition in enabling the smooth transition between discrete behavioral modules during naturalistic behavior.

## ACKNOWLEDGEMENTS

We are grateful to Samuel Pfaff, Hanns Ulrich Zeilhoffer, and Mark H. Tuszynski for generously providing Satb2^CreER^;Ai9, GlyT2^GFP^, and monkey tissue, respectively. We would also like to thank David Ginty for providing PTGFR^CreER^, *Ret^CreER^*, *Split^Cre^*, *Advillin^FlpO^*, *RORβ^CreER^*, and *Robo3^iresCreER^* mice, Takeshi Kaneko for providing *GAD67^GFP^* mice, Martyn Goulding for providing *Lbx1^FlpO^* mice, and Carmen Birchmeier for providing *Lbx1^Cre^* mice. We would like to thank Sean O’Leary for providing animal care. Additionally, we express our appreciation to Brenda Ross for coordinating animal shipments between Rutgers and Dalhousie, as well as providing technical support for EMG recordings.

Financial support was provided by Whitehall Foundation (V.E.A.), Craig H. Neilsen Foundation (V.E.A.); Pew Charitable Trust (V.E.A.), NJ Commission on Spinal Cord Research (N.O-E., M.A.G., V.E.A.); NIH NINDS K01NS116224 (V.E.A.), NIH NINDS R01NS119268 (V.E.A.), NIH NINDS R01NS119268-S1 (M.Gonzalez), NIH NINDS R01NS119268-S2 (O.O.).

## AUTHOR CONTRIBUTIONS

M.A. Gradwell performed and analyzed electrophysiology experiments. N. Ozeri-Engelhard performed and analyzed anatomy experiments with help from A. Upadhyay., M.A. Gradwell, T. Shrier, M. Gandhi, A. Aoki., and G.A. Abbas-Zadeh. N. Ozeri-Engelhard performed and analyzed behavior experiments with help from M. Gonzalez, M.A. Gradwell, A. Upadhyay, J. Katz, J. Keating, N. Yusuf, and S.A. Alomary. O.D. Laflamme performed and analyzed EMG experiments. Moseq analysis was performed by J.T. Eisdorfer, J. Thackray, M. Ricci, and O. Oputa. Y. Hernandez performed all genotyping. M.A. Gradwell and N. Ozeri-Engelhard prepared the figures with input from J.T. Eisdorfer. M.A. Gradwell, N. Ozeri-Engelhard, O.D. Laflamme, T. Akay, and V.E. Abraira designed experiments. V.E. Abraira conceived and supervised the study. V.E. Abraira, wrote the paper with M.A. Gradwell, N. Ozeri-Engelhard, and J.T Eisdorfer all the authors contributed to its editing.

## DECLARATION OF INTERESTS

The authors declare no competing interest.

## DECLARATION of INCLUSION AND DIVERSITY

All the authors in this study actively support inclusion, diversity, and equality in science. The senior author, and several co-authors are from racial and ethnic groups underrepresented in science. Gender and sexual orientation diversity is also highlighted in the composition of this team, including representation of women as a first and last author. One or more of the authors of this paper received support from a program designed to increase minority representation in science, including the PDEP fellowship from Burroughs Wellcome Fund. While citing references scientifically relevant for this work, we also actively worked to promote gender balance in our reference list.

## STAR+METHODS

### Key Resource table

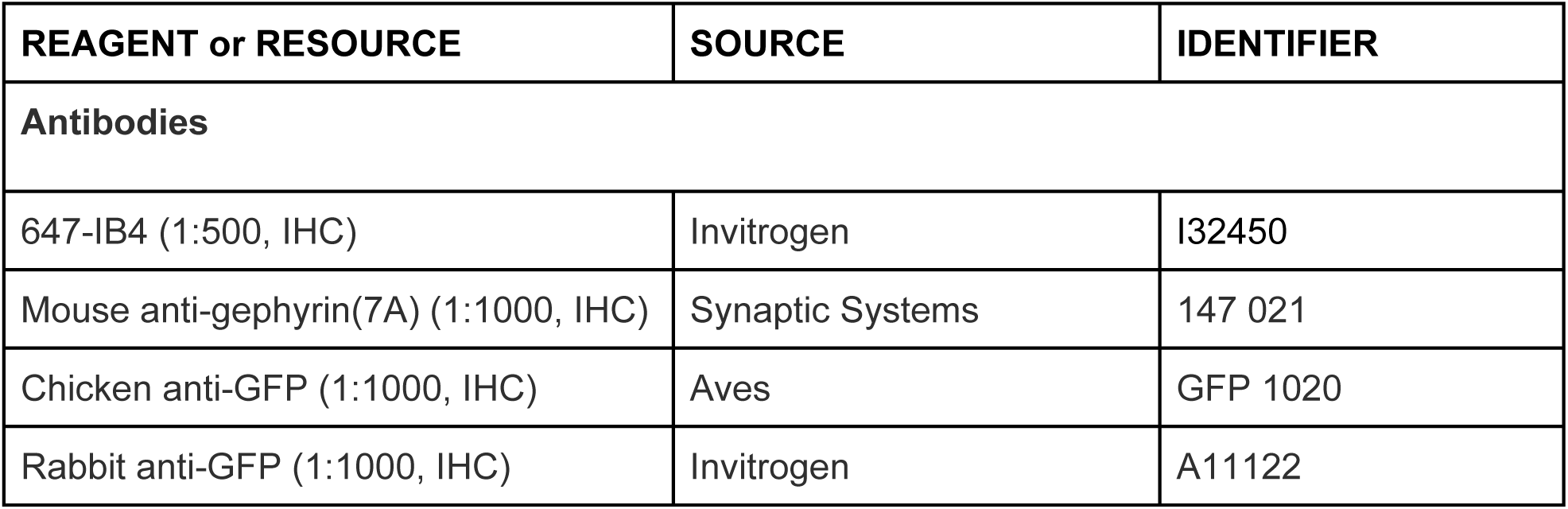
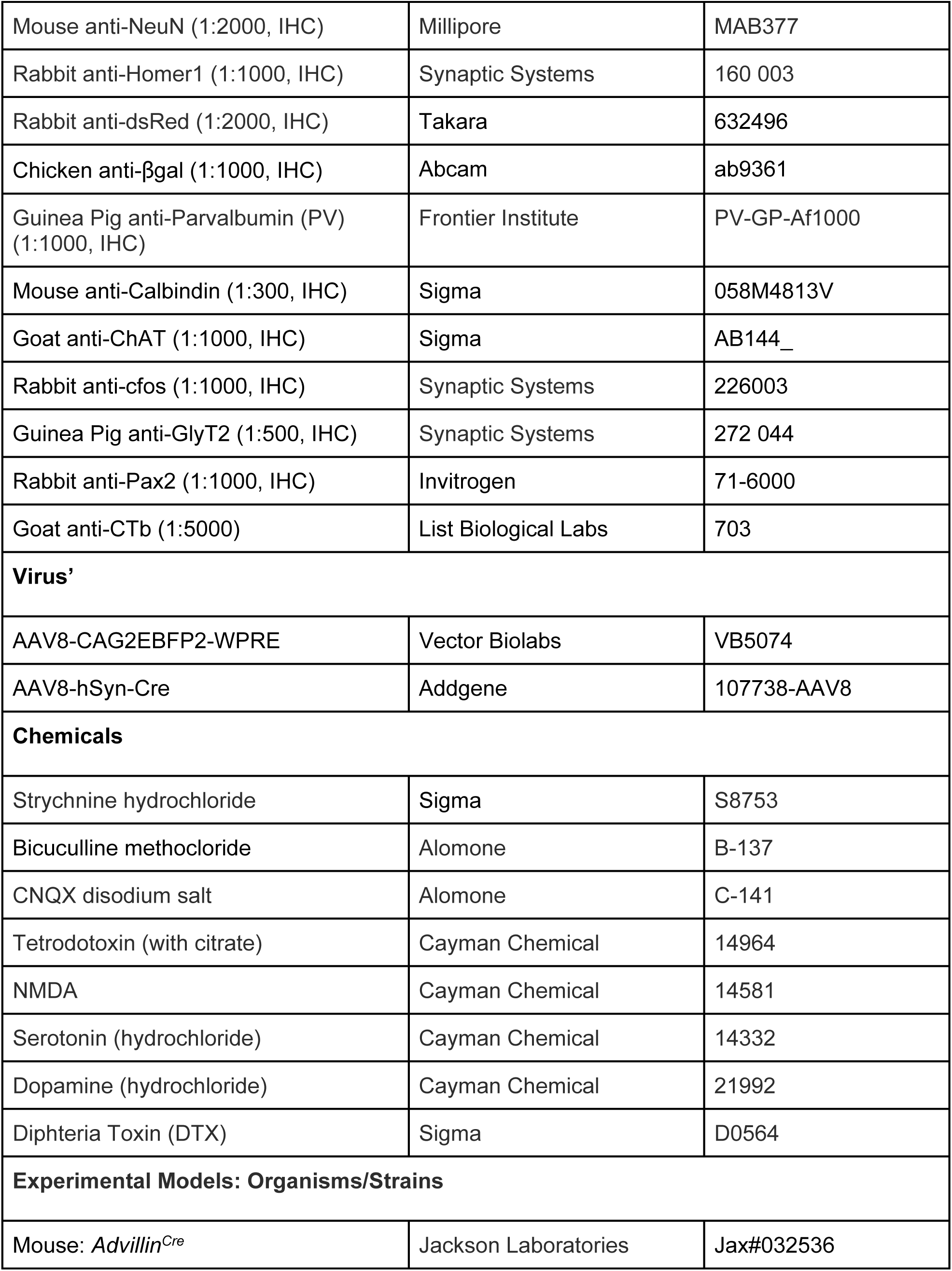
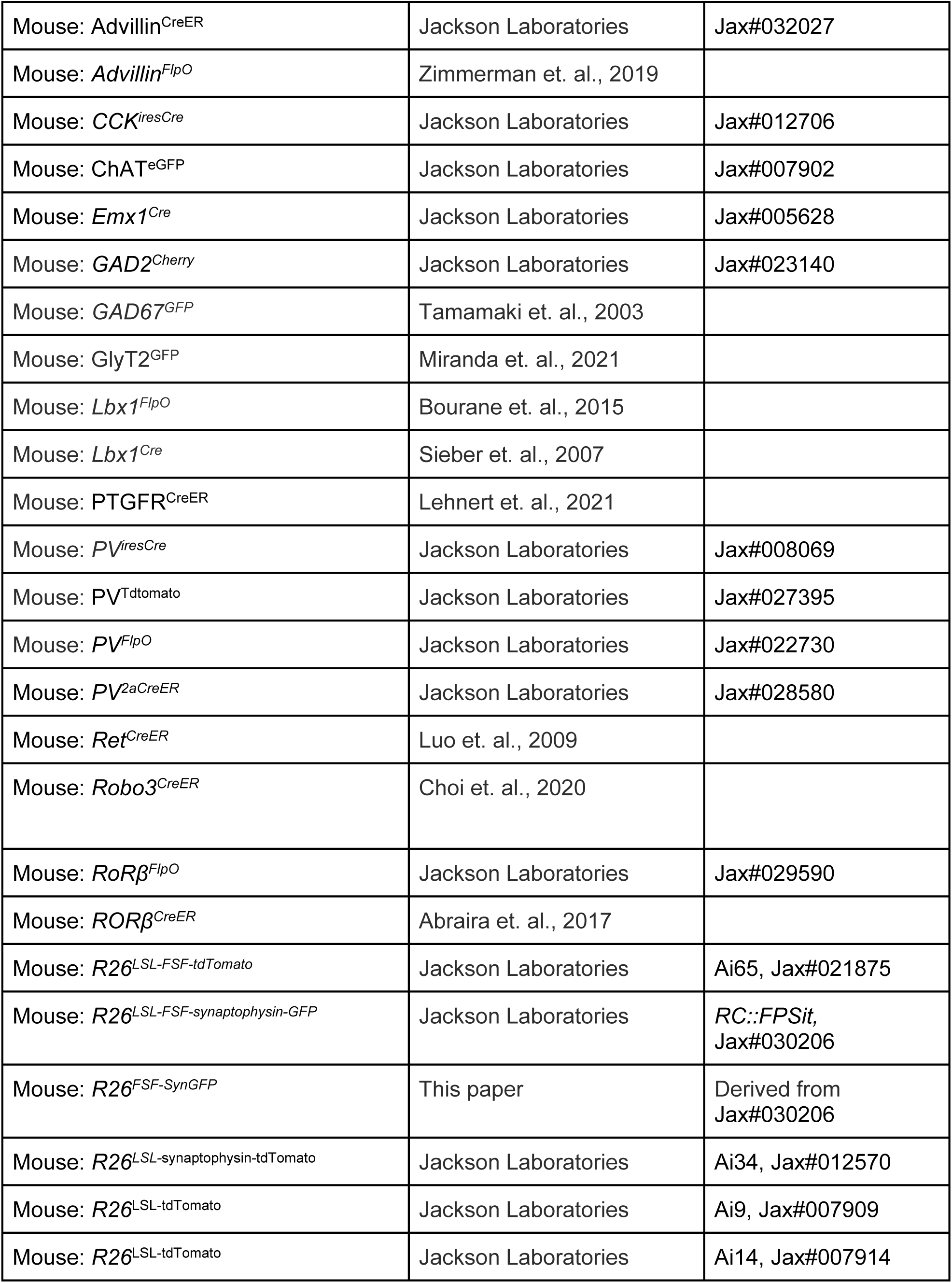
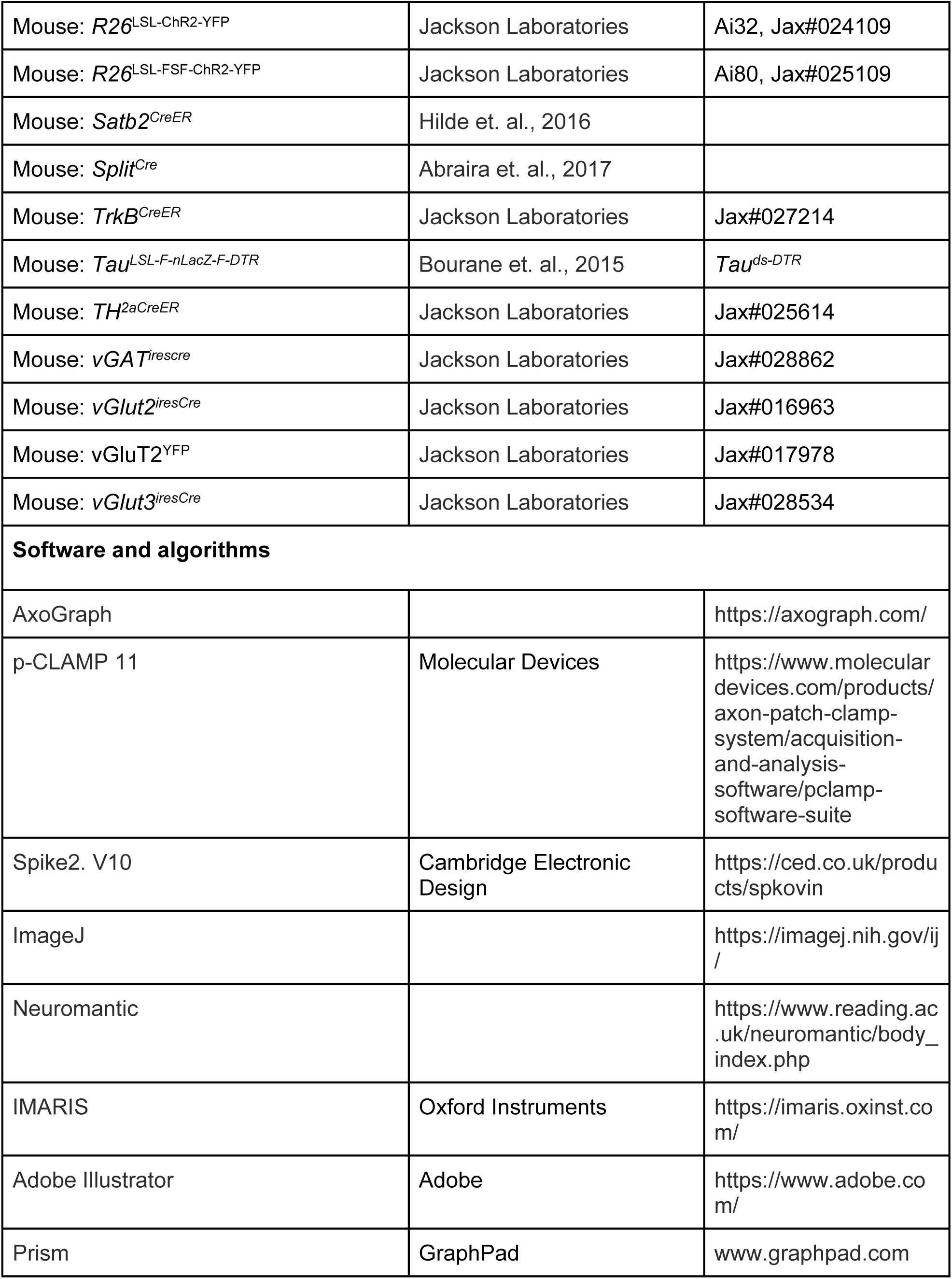

### LEAD CONTACT AND MATERIALS AVAILABILITY

Further information and requests for resources should be directed to and will be fulfilled by the Lead Contact, Victoria E. Abraira.

### EXPERIMENTAL MODEL AND SUBJECT DETAILS

Mouse lines used to target DRG neuron populations include *TH^2aCreER^* (Jax#025614)^49^, *vGlut3^iresCre^* (Jax#028534)^166^, *TrkB^CreER^* (Jax#027214)^57^, PTGFR^CreER 167^, *Ret^CreER^* ^58^, *Split^Cre^* ^57^, *Advillin^Cre^* (Jax#032536)^168^, *Advillin^CreER^* (Jax#032027)^169^, *AdvillinFlpO*^170^, and *PV^2aCreER^* (Jax#028580)^171^. Mouse lines used to target spinal cord interneurons include *vGAT^irescre^* (Jax#028862)^172^, *GAD2^Cherry^* (Jax#023140)^173^, *GAD67^GFP^* ^174^, GlyT2^GFP 175^, *RoRβ^FlpO^* (Jax#029590)^176^, *RORβ^CreER^* ^49^*, Satb2^CreER 3^*^4^, *vGlut2^iresCre^*(Jax#016963)^172^, *CCK^iresCre^* (Jax#012706)^177^, vGluT2^YFP^ (Jax#017978)^178^, *Robo3^CreER^* ^179^, *Lbx1^FlpO^* ^180^, and *Lbx1^Cre^* ^181^. To target parvalbumin+ interneurons we used PV^Tdtomato^ (Jax#027395)^182^, *PV^ireCre^* (Jax#008069)^183^, and *PV^FlpO^* (Jax#022730)^184^ mouselines. For visualization and manipulation of target populations the following mouse lines were used: *R26^LSL-FSF-tdTomato^* (Ai65, Jax#021875)^185^, *R26^LSL-FSF-synaptophysin-GFP^* (*RC::FPSit,* Jax#030206)^186^, R26^FSF-SynGFP^ (derived from *RC::FPSit*), *R26^LSL-^*^synaptophysin-tdTomato^ (Ai34, Jax#012570), *R26*^LSL-tdTomato^ (Ai9, Jax#007909)^184^, *R26*^LSL-tdTomato^ (Ai14, Jax#007914)^184^, *Tau^LSL-F-nLacZ-F-DTR^ (Tau^ds-DTR^)^7^*^0^, *R26*^LSL-ChR2-YFP^(Ai32, Jax#024109)^187^, *R26*^LSL-FSF-ChR2-YFP^(Ai80, Jax#025109)^188^. ChAT^eGFP^ (Jax#007902)^189^ was used to target cholinergic interneurons and motor neurons, *Emx1^Cre^* (Jax#005628)^190^ was used to target cortical neurons, and *Lbx1^Cre^* ^181^ was used to identify dPVs genetic lineage and restrict PV expression to the deep dorsal horn. Transgenic mouse strains were used and maintained on a mixed genetic background (129/C57BL/6). Experimental animals used were of both sexes. With the exception of EMG recordings, housing, surgery, behavioral experiments and euthanasia were performed in compliance with Rutgers University Institutional Animal Care and Use Committee (IACUC; protocol #: 201702589). All mice used in experiments were housed in a regular light cycle room (lights on from 08:00 to 20:00) with food and water available ad libitum. EMG recordings were performed at the lab of Dr. Turgay Akay (Dalhousie, Canada) according to the Canadian Council on Animal Care (CCAC) guidelines and approved by the local council on animal care of Dalhousie University. All mice used in experiments were housed in a regular light cycle room (lights on from 07:00 to 19:00) with food and water available ad libitum.

### METHOD DETAILS

#### Genetic crosses and statistical methods related to individual figures

##### Genetic crosses related to Figure 1

**(B)** Cutaneous and proprioceptive inputs labeled with *PV^2aCreER^;Advillin^FlpO^;Rosa^LSL-FSF-Tomato^/Rosa^FSF-SynGFP^* (2 mg tamoxifen treatment at P21-P23) mice. Quantification of cutaneous and proprioceptive inputs across laminae (3 images, 1 animal). **(C)** L4-L6 spinal cord segments of same quadruple transgenic mice were injected with AAV8-CAG2EBFP2-WPRE, 150 nL, 3 injections **(E)** C-fos analysis n= 4 ‘homecage’ mice and n = 5 ‘walk’ mice: Kruskal Wallis test with Dunn’s correction for multiple comparisons, walk *vs.* rest: LI-III, p = 0.5245; LIV-V_medial_,***p = 0.0005; LIV-V_lateral_,*p = 0.0177; and LVI-X, *p = 0.0386.

##### Genetic crosses and statistical methods related to Figure 2

**(A)** C-fos in excitatory and inhibitory interneurons was generated using *vGAT^iresCre^;Rosa^LSL-Tomato^* mice, n = 4 homecage’ n = 5 ‘walk’. Inhibitory neurons are preferentially recruited by walk: Mann-Whitney test, vGAT+ *vs.* vGAT-, **p = 0.0079. **(B)** oAP incidence, n = 31 GAD67GFP+ and n = 34 GAD67GFP-neurons in 4 *Advillin^CreER^*;*Rosa26^LSL-ChR2^*;*GAD67^GFP^* mice. **(D)** Characterization of sAP incidence: n = 85 from 5 *vGAT^iresCre^*;*R26^LSL-TdTomato^* mice, and n = 85 from 5 *vGluT2^iresCre^*;*R26^LSL-TdTomato^* mice. **(E, F)** Characterization of spontaneous action potential (sAP) discharge: n = 57 from 5 *vGAT^iresCre^*;*R26^LSL-TdTomato^* mice, and n = 32 from 5 *vGluT2^iresCre^*;*R26^LSL-TdTomato^* mice. Inhibitory neurons exhibit **(E)** higher sAP frequency: Mann-Whitney test, ***p = 0.0001 **(E)** and lower sAP coefficient of variation (CV): Mann-Whitney test, ***p = 0.0003. **(H-J)** Characterization of DDH neuron response to depolarizing current injection: n = 86 from 5 *vGAT^iresCre^*;*R26^LSL-TdTomato^* mice, and n = 65 from 5 *vGluT^iresCre^*;*R26^LSL-TdTomato^* mice. Inhibitory neurons exhibit **(H)** lower latency to 1st action potential: Mann-Whitney test, ***p = 0.0003, **(I)** increased AP number, 2-way ANOVA, ****p ≤ 0.0001, **(J)** and increased AP instantaneous frequency, 2-way ANOVA, ****p ≤ 0.0001. (**L)** Analysis of threshold APs showed inhibitory neurons exhibit more hyperpolarized AP threshold: Mann-Whitney test, **p = 0.0024, **(M)** faster AP rise time: Mann-Whitney test, ****p ≤ 0.0001, **(N)** faster AP decay time: Mann-Whitney test, **p = 0.0062, and **(O)** smaller AP half width: Mann-Whitney test, **p = 0.0050.

##### Genetic crosses and statistical methods related to Figure 3

**(A)** PV^TdTomato^ used to identify PV expression in the medial deep dorsal horn. **(B)** Colocalization analysis of inhibitory and excitatory markers with dPVs: n = 2 *vGAT^iresCre^*;*Tau^LSL-F-nLacZ-F-DTR^*;PV^TdTomato^, n = 3 *vGluT2^YFP^*;*PV^TdTomato^* mice. **(C)** Quantification of percent dPVs expressing GAD65, GAD67, or GlyT2: n = 4 *PV^FlpO^*;*R26^FSF-GFP^*;*GAD2^mcherry^* mice, n = 3 *GAD67^GFP^*;PV^TdTomato^ mice, and n = 3 GlyT2^GFP^ mice (stained with GP-PV). **(D)** sAP discharge: n = 316 from 40 PV^TdTomato^ mice. **(E)** AP discharge properties in response to depolarizing current injections: n = 51 dPVs from 9 PV^TdTomato^ mice. **(F)** Left - sensory and proprioceptive inputs labeled with *PV^FlpO^;Advillin^Cre^;R26^LSL-FSF-SynGFP^/R26^LSL-Tomato^* (2 mg tamoxifen treatment at P21-P23). Right - quantification of cutaneous and proprioceptive input onto dPVs: n = 21 from 2 *PV^2aCreER^;Advillin^FlpO^;R26^LSL-FSF-Tomato^/R26^FSF-SynGFP^* (2 mg tamoxifen treatment at P21-P23) mice. dPVs receive more input from cutaneous afferents: Mann-Whitney test, **p = 0.0038. **(G)** Input from Aβ field LTMR to dPVs shown using *PTGFR^CreER^*;*R26^LSL-SynTomato^* mice (2 mg tamoxifen treatment at p21-p22), **(H)** Input from Aβ RA LTMR to dPVs shown using *Ret^CreER^*;*R26^LSL-SynTomato^* mice (2 mg tamoxifen treatment at E10-E11). **(I-J)** Photostimulation of proprioceptive (n = 28, 4 mice), Aβ field (n = 31, 5 mice), Aβ RA (n = 30, 4 mice), Aẟ- (n = 25, 3 mice), and C-LTMRs (n = 18, 2 mice) using *PV^2aCreER^*;*Advillin^FlpO^*;*R26^LSL-FSF-ChR2)^*;PV^TdTomato^ (2 mg tamoxifen at P21-P23), *PTGFR^CreER^*;*Advillin^FlpO^*;*R26^LSL-FSF-ChR2^*;PV^TdTomato^ (2 mg tamoxifen at P21-P23), *Split^Cre^;R26^LSL-ChR2^*;PV^TdTomato^, *TrkB^CreER^*;*Advillin^FlpO^*;*R26^LSL-FSF-ChR2^*;PV^TdTomato^ (2 mg tamoxifen at P21-P23), and *vGlut3^iresCre^*;*Advillin^FlpO^*;R26^LSL-FSF-ChR2;^PV^TdTomato^ mice, respectively. **(K)** Proprioceptive (n = 24, 4 mice) and Aβ field-LTMR (n = 25, 5 mice) excitatory inputs to dPVs are significantly larger than Aβ RA-LTMR (n = 15, 4 mice) inputs: Kruskal-Wallis test with Dunn’s correction for multiple comparisons, Proprioceptive *vs.* Aβ field-LTMR, p = 0.5241; Proprioceptive *vs.* Aβ RA-LTMR, ***p = 0.0008; Aβ RA-LTMR *vs.* Aβ Field-LTMR, *p = 0.0399. **(L)** Polysynaptic oEPSCs recorded in a subset of proprioceptor (n= 15), Aβ field (n = 15), Aβ RA (n = 7), Aẟ-(n = 1), and C-LTMRs (n = 2) photostimulation experiments. **(M)** Recordings were performed in 29 dPV neurons from 3 *CCK^iresCre^*;Lbx1^FlpO^;*R26^LSL-FSF-ChR2^* (Ai80);PV^TdTomato^ mice. **(N)** Polysynaptic oIPSCs recorded in a subset of proprioceptor (11/18), Aβ field (7/15), Aβ RA (3/13), Aẟ-(0/9), and C-LTMRs (0/2) photostimulation experiments **(O)** Recordings performed in 25 dPV neurons from 2 *RORβ^CreER^*;Lbx1^FlpO^;*R26^LSL-FSF-ChR2 (Ai80)^*;PV^TdTomato^ (2 mg tamoxifen treatment at P18-P19, mostly labeling LTMR-recipient zone RORβ interneurons) mice.

##### Genetic crosses and statistical methods related to Figure 4

**(A)** Colocalization of dPVs (red) with Lbx1 shown using *Lbx1^Cre^;Tau^LSL-F-nLacZ-F-DTR^*;PV^TdTomato^ mouse, n = 3. **(B)** Images of lumbar spinal cord from PV^Tdtomato^ (left) and *PV^FlpO^*;*Lbx1^Cre^*;*Rosa26^LSL-FSF-TdTom^* (right) mice. Contour plot was generated using 1 PV^Tdtomato^ mouse (left) and 2 *PV^FlpO^*;*Lbx1^Cre^*;*Rosa26^LSL-FSF-TdTom^* mice. **(C)** Contour density plot generated using 2 *PV^FlpO^*;*Lbx1^Cre^*;*Rosa26^LSL-FSF-SynaptophysinGFP^* mice. **(D)** *PV^FlpO^*;*Lbx^1Cre^*;*Rosa^LSL-FSF-SynGFP^*;*Rosa^LSL-FSF-TdTomato^* was used to demonstrate dPVs’ mutual connections. *PV^FlpO^*;*Lbx^1Cre^*;*Rosa^LSL-FSF-SynGFP^* stained with mouse-anti calbindin was used to demonstrate dPVs’ connections to Renshaw cells. *PV^FlpO^*;*Lbx^1Cre^*;*Rosa^LSL-FSF-SynGFP^* stained with goat-anti ChAT was used to demonstrate dPVs’ connections to V0c and MNs. V0c neurons were identified in lamina X, around the central canal, while MNs were identified by their location in lamina IX and large soma size. **(E-G)** Responses in dPVs to photostimulation of dPVs: n = 23 in 3 *PV^FlpO^*;*Lbx1^Cr^*^e^;*Rosa26^LSL-FSF-ChR2^*;PV^TdTomato^ mice. Responses in V0c (n = 10, 4 mice), lateral MNs (n = 30, 4 mice), medial MNs (n = 8, 3 mice), and putative foot MNs (n = 14, 4 mice) to photostimulation of dPVs recorded in *PV^FlpO^*;*Lbx1^Cre^*;*Rosa26^LSL-FSF-ChR2^*;ChATe^GFP^ mice.

##### Genetic crosses and statistical methods related to Figure 5

**(A)** Percent c-fos+dPVs increased in walk *vs.* rest: Mann Whitney test, *p = 0.0286. **(D)** NMDA, 5HT, DA cocktail increased holding current recorded in 9 dPVs from 3 PV^TdTomato^ animals: paired t-test, ***p = 0.0004. **(E)** NMDA, 5HT, DA cocktail increased resting membrane potential (RMP) recorded in 19 dPVs from 4 PV^TdTomato^ animals: paired t-test, ****p ≤ 0.0001. **(H)** NMDA, 5HT, DA cocktail increased spontaneous AP instantaneous frequency (sAP-IF) recorded in 7 dPVs from 4 PV^TdTomato^ mice: paired t-test, * p = 0.0142. **(J)** Bicuculline, Strychnine, CNQX cocktail increased sAP recorded in 11 dPVs from 2 PV^TdTomato^ mice: Wilcoxon matched-pairs signed rank test, **p = 0.0098. **(O)** Sensory afferent input that evokes AP discharge (n = 29) is significantly larger than afferent input that does not (n = 22): Mann-Whitney test, ****p ≤ 0.0001.

##### Genetic crosses and statistical methods related to Figure 6

**(B-G)** EMG recordings were obtained from 6 dPV^norm^ (*PV^FlpO^*/*Lbx1^Cre^*;*Tau^LSL-F-nLacZ-F-DTR^*) and 9 dPV^abl^ (*PV^FlpO^*;*Lbx1^Cre^*;*Tau^LSL-F-nLacZ-F-DTR^*) mice intrathecally injected with 50 ng DTx. **(D)** We observed no difference between Sural *vs.* Saphenous nerve stimulation in paired muscles: dPV^norm^ ST, Unpaired t-test, p = 0.8800; dPV^abl^ ST, Unpaired t-test, p = 0.1601; dPV^norm^ GS, Unpaired t-test, p = 0.3756; dPV^abl^ GS, Unpaired t-test, p = 0.4185. Therefore, we combined sural and saphenous nerve stimulation. dPV^abl^ mice exhibit shorter cutaneous evoked latency in flexors (n = 17 dPV^norm^ and 21 dPV^abl^): Unpaired t-test, ****p ≤ 0.0001, and extensors (n = 11 dPV^norm^ and n = 11 dPV^abl^): Unpaired t-test, ***p = 0.004. **(G)** We observed no difference between Sural *vs.* Saphenous nerve stimulation in paired muscles: dPV^norm^ ST, Unpaired t-test, p = 0.7336; dPV^abl^ ST, Mann-Whitney test, p = 0.3968; dPV^norm^ GS, Unpaired t-test, p = 0.2812; dPV^abl^ GS, Mann-Whitney test, p = 0.9286. Therefore, we combined sural and saphenous nerve stimulation. dPV^abl^ mice exhibit reduced cutaneous evoked inhibition in flexors (n = 19 dPV^norm^ and 14 dPV^abl^): Mann-Whitney test, ****p ≤ 0.0001, and extensors (n = 9 dPV^norm^ and n = 10 dPV^abl^): Mann-Whitney test, ****p ≤ 0.0001.

##### Genetic crosses and statistical methods related to Figure 7

**(C-E)** Joint angles were generated using 93 steps from 3 dPV^norm^ (*PV^FlpO^/Lbx1^Cre^*;*Tau^LSL-F-nLacZ-F-DTR^*) and 81 steps from 4 dPV^abl^ (*PV^FlpO^*;*Lbx1^Cre^*;*Tau^LSL-F-nLacZ-F-DTR^*) mice. For statistical analysis we used a linear mixed effects regression model with mouse ID as a random effect and the Holm-Bonferroni method was used to correct for multiple comparisons. **(C)** dPV^abl^ *vs.* dPV^norm^ hip PEP, p = 0.1861 and AEP, p = 0.2709 **(D)** knee PEP, **p = 0.0035 and AEP, p = 0.8236; **(E)** ankle PEP, **p = 0.0069 and AEP, p = 0.2671. **(G-K)** Digigait data was obtained from 7 dPV^norm^ (*PV^FlpO^/Lbx1^Cre^*;*Tau^LSL-F-nLacZ-F-DTR^*) and 10 dPV^abl^ (*PV^FlpO^*;*Lbx1^Cre^*;*Tau^LSL-F-nLacZ-F-DTR^*) mice. For statistical analysis we used a linear mixed effects regression model with mouse ID as a random effect and the Holm-Bonferroni method was used to correct for multiple comparisons. The following number of steps were used for dPV^norm^ and dPV^abl^, respectively, at each speed: 20 cm/s -133 and 196; 40 cm/s - 108 and 209; 60 cm/s - 61 and 138; 80 cm/s - 26 and 61. **(G)** dPV^abl^ *vs.* dPV^norm^ stride frequency at: 20 cm/s, p = 0.3511; 40 cm/s, *p = 0.0185; 60 cm/s, ****p = 0.0001; 80 cm/s, **p = 0.0096. **(H)** dPV^abl^ *vs.* dPV^norm^ stride length at: 20 cm/s, p = 0.4513; 40 cm/s, *p = 0.0464; 60 cm/s, ****p ≤ 0.0001; 80 cm/s, p = 0.4513. **(I)** dPV^abl^ *vs.* dPV^norm^ stride frequency at: 20 cm/s, p = 0.2345; 40 cm/s, *p = 0.0168; 60 cm/s, ***p = 0.0005; 80 cm/s, *p = 0.0168.

##### Genetic crosses and statistical methods related to Figure 8

Motion sequencing was performed on 9 dPV^norm^ (*PV^FlpO^/Lbx1^Cre^*;*Tau^LSL-F-nLacZ-F-DTR^*) and 9 dPV^abl^ (*PV^FlpO^*;*Lbx1^Cre^*;*Tau^LSL-F-nLacZ-F-DTR^*) mice, all male. The Benjamini-Hochberg bootstrapping method (a = 0.05) was used to correct for multiple hypothesis testing and control for the false discovery rate. Statistical testing procedures were performed previously described^115^. In brief, group comparisons were performed for each syllable by taking 1,000 bootstrap samples of the syllable’s frequency for each group and a z-test was performed on these two distributions. **(B)** dPV^abl^ *vs.* dPV^norm^ syllable usages: syllable ID 0, upregulated, ***p = 0.00009; and syllable ID, downregulated, 1 ***p = 0.00004. **(D)** A multinomial logistic regression classifier was used to compare the expressive power of behavioral syllable usages (57 dimensions) and transition probabilities (most frequent transitions on average). Leave-one-out validation was performed on train and test datasets in a 70:30 split using scikit-learn in Python3. **See Figure S8** for the reported F-Statistic for the classifier model. The following scores were observed when comparing groups in the classifier confusion matrix: True label (TL) dPV^norm^ *vs.* Predicted Label (PL) dPV^norm^: 1; TL dPV^norm^ *vs.* PL dPV^abl^: 0; TL dPV^abl^ *vs.* PL dPV^norm^: 0.11; and TL dPV^abl^ *vs.* PL dPV^abl^: 0.89. **(E-F)** Logistic regression analysis unearthed a series of distinct syllables and behavioral transitions between syllables that carry a large weight in discriminating between dPV^abl^ and dPV^norm^ groups. The average testing F1 across all splits was 0.849 for usages, and 0.655 for transition probabilities. **(E)** Four locomotor syllables were identified as demonstrating large contribution weight to, and high probability of inclusion in, the classifier model: syllable ID 18 (head down to head up locomotion, ∼9 cm/s), 32.8 average weight, 0.47 probability of inclusion, upregulated in dPV^abl^ group; syllable ID 14 (head down locomotion, ∼10.5 cm/s), 39.1 average weight, 0.3 probability of inclusion, downregulated; syllable ID 8 (locomotion to medium rear, ∼12 cm/s), 13.5 average weight, 0.09 probability of inclusion, downregulated; and syllable ID 30 (head down locomotion, ∼21 cm/s), -29.5 average weight, 0.29 probability of inclusion, downregulated (See **Table S1** for all syllables with large weights). **(F)** Transition probabilities identified as demonstrating large contribution weight to, and high probability of inclusion in, the classifier model. Examples shown: syllable IDs 4 → 1, -376.5 average weight, 0.15 probability of inclusion, downregulated in the dPV^abl^ group; syllable IDs 4 → 30, 53.0 average weight, 0.19 probability of inclusion, upregulated; and syllable IDs 1 → 30, -292.7 average weight, 0.75 probability of inclusion, downregulated (See **Table S2** for all transition probabilities with large weights).

#### Tamoxifen treatment

Tamoxifen was dissolved in ethanol (20 mg/ml), mixed with an equal volume of sunflower seed oil (Sigma), vortexed for 5-10 min and centrifuged under vacuum for 20-30 min to remove the ethanol. The solution was kept at −80°C and delivered via oral gavage to pregnant females for embryonic or postnatal treatment. For all analyses, the morning after coitus was designated as E0.5 and the day of birth as P0.

#### Immunohistochemistry and free-floating sections

Male and female P30-P37 mice were anesthetized with isoflurane and perfused transcardially (using an in-house gravity driven-perfusion system) with heparinized-saline (∼30 sec) followed by 15-20 minutes of 4% paraformaldehyde (PFA) in PBS at room temperature (RT). Vertebral columns were dissected and post-fixed in 4% PFA at 4°C for 2-24 hr. Sections were collected using a vibrating microtome (Leica VT1200S) and processed for immunohistochemistry (IHC) as described previously^191^. Unless otherwise mentioned, transverse sections (50-60 µm thick) were taken from limb level lumbar spinal cord (L4-6). Free floating sections were rinsed in 50% ethanol (30 min) to increase antibody penetration, followed by three washes in a high salt PBS buffer (HS-PBS), each lasting 10 min. The tissue was then incubated in a cocktail of primary antibodies made in HS-PBS containing 0.3% Triton X-100 (HS-PBSt) for 48 at 4°C. Next, tissue was washed 3 times (10 minutes each) with HS-PBSt, then incubated in a secondary antibody solution made in HS-PBSt for 24 hr at 4°C. Immunostained tissue was mounted on positively charged glass slides (41351253, Worldwide Medical Products) and coverslipped (48393-195, VWR) with Fluoromount-G mounting medium (100241-874, VWR).

#### Image acquisition and analysis

Images were captured with a Zeiss LSM 800 confocal or a Zeiss axiovert 200M fluorescence microscope. Images for cell counts were taken with 10x or 20x air objectives, and images of synaptic contacts were taken using a 40x oil objective. ImageJ (cell count plug-in) was used for colocalization analysis of cell bodies. Imaris (spot detection plug-in) was used for synaptic analysis, quantification of synaptic terminals, and generation of contour density plots. Only puncta apposing a postsynaptic marker (Homer1 and Gephyrin for excitatory and inhibitory synapses, respectively) were included for analysis. Synaptic analysis was performed separately for somas and proximal dendrites (up to 50 µm from the cell body). For quantification of proprioceptive and cutaneous synaptic terminals across the dorsoventral axis, synaptophysin GFP (all sensory) was used to mask the tomato (proprioceptive fibers) channel to generate a separate channel of red and green proprioceptive puncta. To generate contour density plots, Imaris spot detection plug-in was used to detect the spatial coordinates of cell bodies/synaptic terminals. Following detection, coordinate values were corrected and expressed relative to common reference points (central canal, dorsal, lateral and ventral edges of the gray matter). Corrected coordinates were used to obtain two-dimensional kernel density estimation utilizing the MATLAB ‘ksdensity’ function. Kernel density estimations were graphically displayed as contour plots, using the MATLAB ‘contourf’ function.

#### C-fos induction

Mice were transferred to a holding area adjacent to the testing room at least 7 days prior to the experiment, to decrease stress during the task. 6-8 weeks old mice were randomly placed in one of two groups: homecage and walk. All mice were habituated to the testing room for 30 minutes. Homecage mice were kept in their homecage for 60 minutes. “Walk” group walked on a treadmill (Mouse Specifics, Inc, Boston, MA, USA) at 20 cm/s for 2 x 20 minutes with a 5-minute break. 60 minutes following the completion of the task, mice were anesthetized with isoflurane and perfused for immunohistochemistry. Sections stained with chicken-β-galactosidase (ab9361, Abcam), mouse-NeuN (MAB377, Millipore), and rabbit-c-fos (226003, Synaptic systems) were used to acquire 20x images of the entire hemisection, stitches with zen. Images were then used for quantification of c-fos expressing neurons (normalized to total NeuN) by lamina and cfos expressing inhibitory (vGAT+cfos+NeuN+) and excitatory (vGAT-cfos+NeuN-) neurons in the medial portion of laminas IV-V. Alternate sections from the same animals stained with Guinea pig-PV (PV-GP-Af1000, Frontier institute) and c-fos, were used to acquire 20x images of the medial deep dorsal horn to quantify the fraction of dPVs that express cfos in each of the conditions. All images were acquired with the Zeiss LSM 800 confocal microscope.

#### CTb injections

##### Muscle

*PV^FlpO^;Lbx1^Cre^;R26^LSL-FSF-SynaptophysinGFP^* mice were anesthetized (5% induction, 2-3% maintenance), and the hair above one of their hindlimbs was shaved. The skin was wiped with 70% ethanol and cut to expose the hindlimb muscles. Injections of 0.125% Alexa fluor 555-conjugated CTb (2-3 sites along each muscle) were made into the knee extensor vastus lateralis (VL), knee flexor biceps femoris (BFP), ankle extensor lateral gastrocnemius (Gs), and ankle flexor tibialis anterior (TA). Animals were perfused 5 days after CTb injection and tissue was processed for IHC as described above. Free floating sections were immunostained with chicken-GFP (GFP-1010, Aves) and mouse-Gephyrin (147 021, Synaptic systems) primary antibodies, and appropriate secondary antibodies. 40x images of CTb-labeled motor neurons were taken at the Zeiss LSM 800 confocal microscope and ImageJ was used for synaptic analysis.

##### Paw

Mice were anesthetized (5% induction, 2-3% maintenance). Injections of 1% unconjugated CTb (∼7µL across four sites) was injected into the ventral paw of Wild type mice to label paw sensory input. Animals were perfused 5 days after CTb injection and tissue was processed for IHC as described above. Free floating sections were immunostained with goat-CTb (list-labs). 40x images of CTb-labeled afferent inputs were taken at the Zeiss LSM 800 confocal microscope and ImageJ was used for synaptic analysis.

#### Spinal cord injections

Mice were anesthetized with isoflurane (5% induction, 1.5-2% maintenance). A small incision was made in the skin and underlying musculature and adipose tissue was teased apart to reveal the vertebral column. Vertebral surfaces were cleaned with fine forceps to reveal the dorsal surface of the spinal cord. AAV8-hSyn-Cre (107738-AAV8, Addgene) or AAV8-CAG2-EBFP2-WPRE (VB5074, Vector Biolabs), were injected into 3-4 sites in the L4-L6 spinal cord through a pulled glass needle, ∼150 nL per site. Following injections, the overlying muscle and the skin incision was sutured. Mice were allowed to recover from anesthesia and returned to their home cages. 3-4 weeks following the injection, mice were perfused, and their tissue was processed for immunohistochemistry.

#### Timing of dPV neurogenesis

Pregnant PV^TdTomato^ females were intraperitoneally (i.p) injected with 50 mg/kg EdU (Fisher Scientific, C10337) at E10.5, E11.5 and E12.5. PV^TdTomato^ F1 progeny (identified with PCR genotyping) were perfused at P30-P37 and L4-L6 spinal cord sections were collected using a vibrating microtome (Leica VT1200S). EdU was revealed as previously described using a click reaction^192^. After EdU staining, slices were additionally stained with rabbit-DsRed (632496, Takara) and appropriate secondary antibodies as described above, for amplification of PV fluorescence. Following, 20x confocal images of the medial deep dorsal horn were taken and used to quantify EdU expressing-dPVs at each embryonic day.

#### Neuronal reconstructions and morphometric analysis

Sagittal sections of *PV^FlpO^*;*Lbx1^Cre^*;*R26^LSL-FSF-TdTomato^* lumbar spinal cords were immunostained with Guinea-pig PV, and 20x, z stack images were taken on a Zeiss LSM 800 confocal microscope. Stacks were manually reconstructed using the Neuromantic software. Reconstructions were then opened with ImageJ to measure somatic area and neurite length. Spine counts were performed with the cell counter plug-in to calculate spine density for each neuron (number of spines/ total neurite length). Sholl regression plug-in was used to perform sholl regression, to characterize neuronal arbors, as described as previously described ^193^. Briefly, the plug-in creates a series of concentric spheres around the soma and counts the number of intersections with each sphere. A linear regression is fitted to the sampled data (number of intersections vs distance from the soma) and the sholl regression coefficient (the slope of the linear regression) is calculated. The Regression coefficient represents the decay in the number of branches with distance from the cell body where higher values denote faster decay.

#### Electrophysiology

Recordings were made from male (n = 33) and female (n = 32) mice (5.1 +/-0.2 wks). Mice were anesthetized with ketamine (100 mg/kg i.p), decapitated, and the lumbar enlargement of the spinal cord rapidly removed in ice-cold sucrose substituted artificial cerebrospinal fluid (sACSF) containing (in mM): 250 sucrose, 25 NaHCO3, 10 glucose, 2.5 KCl, 1 NaH2PO4, 6 MgCl2, and 1 CaCl2. Sagittal or transverse slices (200µm thick) were prepared using a vibrating microtome (Leica VT1200S). Slices were incubated for at least 1 hour at 22-24°C in an interface chamber holding oxygenated ACSF containing (in mM): 118 NaCl, 25 NaHCO3, 10 glucose, 2.5 KCl, 1 NaH2PO4, 1 MgCl2, and 2.5 CaCl2.

Following incubation, slices were transferred to a recording chamber and continually superfused with ACSF bubbled with Carbogen (95% O2 and 5% CO2) to achieve a pH of 7.3-7.4. All recordings were made at room temperature (22-24°C) and neurons visualized using a Zeiss Axiocam 506 color camera. Recordings were acquired in cell-attached (holding current = 0mV), voltage-clamp (holding potential -70mV), or current-clamp (maintained at −60mV) configuration. Patch pipettes (3-7 MΩ) were filled with a potassium gluconate-based internal solution containing (in mM): 135 C6H11KO7, 8 NaCl, 10 HEPES, 2 Mg2ATP, 0.3 Na3GTP, and 0.1 EGTA, (pH 7.3, adjusted with KOH, and 300 mOsm) to examine AP discharge and excitatory synaptic transmission. A cesium chloride-based internal solution containing (in mM): 130 CsCl, 10 HEPES, 10 EGTA, 1 MgCl2, 2 Na-ATP, 2 Na-GTP, and 5 QX-314 bromide (pH 7.3, adjusted with CsOH, and 280 mOsm) was used to record inhibitory synaptic transmission. No liquid junction potential correction was made, although this value was calculated at 14.7 mV (22°C). Slices were illuminated using an X-CITE 120LED Boost High-Power LED Illumination System that allowed visualization of Td-Tomato expressing dPVs with TRITC filters, and visualization of GFP fluorescence and ChR2 photostimulation using FITC filter set.

AP discharge patterns were assessed in the current clamp from a membrane potential of −60 mV by delivering a series of depolarizing current steps (1 s duration, 20 pA increments, 0.1 Hz), with rheobase defined as the first current step that induced AP discharge. Spontaneous action potential discharge was assessed using cell-attached voltage clamp recordings, with minimal holding current (< 20 mV). For optogenetic circuit mapping, photostimulation intensity was suprathreshold (24 mW), duration 1 ms (controlled by transistor-transistor logic pulses), 0.1 Hz. This ensured generation of action potential discharge in presynaptic populations and allowed confident assessment of postsynaptic currents in recorded neurons.

All data were amplified using a MultiClamp 700B amplifier, digitized online (sampled at 20 kHz, filtered at 5 kHz) using an Axon Digidata 1550B, acquired and stored using Clampex software. After obtaining the whole-cell recording configuration, series resistance, input resistance, and membrane capacitance were calculated (averaged response to -5mV step, 20 trials, holding potential -70mV).

#### Electrophysiology Analysis

All data were analyzed offline using AxoGraph X software.

AP discharge was classified according to previously published criteria^44^. Briefly, tonic firing (TF) neurons exhibited continuous repetitive AP discharge for the duration of the current injection; initial bursting (IB) neurons were characterized by a burst of AP discharge at the onset of current injection; delayed firing (DF) neurons exhibited a clear interval between current injection and the first AP; and single spiking (SS) neurons that discharge one or two APs at depolarization step onset. In addition, we also divided the TF population into two subpopulations; TF non-accommodating (TF_Na_) neurons maintained the TF profile with incremental current injection; TF accommodating (TF_Ac_) neurons could not maintain the TF profile with incremental current injection, and AP duration dropped below 900 ms or reduced by more than 10%. In the analysis of AP discharge, individual APs elicited by step current injection were captured using a derivative threshold method (dV/dt > 15 V/second) with the inflection point during spike initiation defined as the AP threshold. Individual AP properties for each neuron were assessed from the first spike generated at the rheobase. AP latency was measured as the time difference between the stimulus onset and AP threshold. The difference between the AP threshold and its maximum positive peak was defined as the AP peak. AP half-width was measured at 50% of AP peak. AP rise and decay time were measured from 10-90% of peak. AP afterhyperpolarization (AHP) amplitude was taken as the difference between the AP threshold and the maximum negative peak following the AP. AHP half-width was measured at 50% of AHP peak. AP threshold adaptation was determined by dividing the AP threshold of the last AP by the AP Threshold of the first AP evoked by a step current. AP number was the number of spikes discharged in the entire response. Instantaneous frequency is the average instantaneous frequency across a step current.

Optogenetically-evoked excitatory postsynaptic currents (oEPSCs) and inhibitory postsynaptic currents (oIPSCs) were measured from baseline, just before photostimulation. The peak amplitude of responses was calculated from the average of 10 successive trials. Several parameters were considered for determining whether a photostimulation-evoked synaptic input was monosynaptic or polysynaptic. The latency of oPSCs was measured as the time from photostimulation to the onset of the evoked current. The “jitter” in latency was measured as the standard deviation in latency of 10 successive trials. Importantly, the latency of monosynaptic inputs was much shorter, there was minimal jitter in the onset of responses between trials, and reliability (percentage of photostimulation trials to evoke a response) was higher than those deemed polysynaptic inputs. To assess the contribution of different neurotransmitter systems to photostimulation responses, various synaptic blockers were sequentially applied.

#### Diphtheria Toxin preparation and delivery

DTx was dissolved in 1xPBS to 100 ug/ml and kept at −20°C. On the day of injection, DTx was diluted 1:10 in a hyperbaric solution (filtered 0.9% NaCl, 8% dextrose) as previously described^89^. P21-P28 mice carrying the *PV^FlpO^*;*Lbx1^Cre^*;*Tau^LSL-F-nLacZ-F-DTR^* (dPVs^abl^) or *PV^FlpO^*/*Lbx1^Cre^*;*Tau^LSL-F-nLacZ-F-DTR^* (dPVs^norm^) alleles, received Diphtheria Toxin (DTx, D0564-1MG, Sigma) on day 1 and 3 (5 ul of 10 ug/ml DTx, overall 50 ng) through intrathecal (I.T.) injection to the L5-L6 intervertebral space. DTx was delivered with a 25 ul Hamilton syringe (72310-388) with a 30G needle and performed as previously described^194^. Briefly, mice were anesthetized with isoflurane and the hair above their lower back was removed. Under anesthesia, exposed skin was wiped with 70% ethanol. Next, mice were held firmly by the pelvic girdle with their skin taut in one hand, while the other hand was used to gently rotate the base of the tail to find the midline of the spine. Needle was inserted through the skin along the midline, between the pelvic girdles to the L5-L6 intervertebral space. An occurrence of a sudden tail flick indicated successful entry into the intradural space.

#### Quantification of DTx mediated ablation

Following behavioral testing, dPV^norm^ (*PV^FlpO^*/*Lbx1^Cre^*;*Tau^LSL-F-nLacZ-F-DTR^*) and dPV^abl^ (*PV^FlpO^*;*Lbx1^Cre^*;*Tau^LSL-F-nLacZ-F-DTR^*) mice were perfused, their spinal cord and cerebellum were dissected, and sectioned using a vibrating microtome (Leica VT1200S). For consistency, sections were collected at the levels of the mid cervical enlargement, mid thoracic, mid lumbar enlargement, and the midline of the cerebellum. Sections were immunostained with Guinea pig-PV and mouse-NeuN, and 20x images were taken using the Zeiss axiovert 200M fluorescence microscope. 3 spinal cord (at each level) and 2 cerebellum images from each animal were used for quantification. PV neurons were counted in the superficial dorsal horn (sPVs), deep dorsal horn (dPVs) and molecular layer of the cerebellum. sPVs were normalized to NeuN, dPVs were normalized to sPVs (to account for difference in staining success), and PV neurons in the molecular layer were normalized to area.

#### Construction of EMG electrodes

EMG electrodes were made using multi-stranded, Teflon coated annealed stainless-steel wire (A-M systems, 793200). The construction of the EMG electrode and nerve cuff was previously described in detail^61,195,196^. Two nerve cuff electrodes and six EMG recording electrodes were attached to the headpiece pin connector (female, SAM1153-12; DigiKey Electronics Thief River Falls, MN) and covered with 3D printed cap.

#### EMG electrode implantation surgeries

Experiments were conducted on 4-5 month-old males and females dPVs^norm^ (*PV^FlpO^*/*Lbx1^Cre^*;*Tau^LSL-F-nLacZ-F-DTR^*) and dPVs^abl^ (*PV^FlpO^*;*Lbx1^Cr^*^e^;*Tau^LSL-F-nLacZ-F-DTR^*) mice. All surgeries were performed in aseptic conditions and on a warm water-circulated heating pad, maintained at 42°C. Each mouse underwent an electrode implantation surgery as previously described^94,96^. Briefly, animals were anesthetized with isoflurane (5% for inductions, 2% for maintenance of anesthesia), ophthalmic eye ointment was applied to the eyes, and the skin of the mice was sterilized with three-part skin scrub using hibitane, alcohol and povidone-iodine. Prior to each surgery, buprenorphine (0.03 mg/kg) and meloxicam (5 mg/kg) were injected subcutaneously as analgesics while the animals were still under anesthesia. Additional buprenorphine injections were performed in 12-hour intervals for 48 hours.

A set of six bipolar EMG electrodes and two nerve stimulation cuffs were implanted into each mouse as the following: small incisions were made on the shaved areas (neck and both hind legs), and each bipolar EMG electrode and the nerve cuff electrodes were led under the skin from the neck incisions to the leg incisions, and the headpiece connector was stitched to the skin around the neck incision. EMG recording electrodes were implanted into the left knee flexor (semitendinosus, St_l_), ankle flexor (tibialis anterior, TA_l_) and extensor (gastrocnemius, Gs_l_) as well as the right knee flexor (semitendinosus, St_r_) and extensor (Vastus Lateralis, VL_r_) and the left ankle extensor (gastrocnemius, Gs_r_). Nerve stimulation electrodes were implanted in the left saphenous nerve and right sural nerve to evoke cutaneous feedback. After surgery, the anesthetic was discontinued, and mice were placed in a heated cage for three days before returning to a regular mouse rack. Food mash and hydrogel were provided for the first three days after the surgery. Any mouse handling was avoided until mice were fully recovered, and the first recording session started at least ten days after electrode implantation surgeries.

#### EMG recording sessions

After animals fully recovered (∼10 days) from electrode implantation surgeries, recording session went as follows: Under brief anesthesia with isoflurane, a wire to connect the headpiece connector with the amplifier and the stimulation insulation units (ISO-FLEX -AMPI, Jerusalem, Israel or the DS4 -Digitimer, Hertfordshire, United Kingdom) were attached to the mouse. Anesthesia was discontinued, and the mouse was placed on a mouse treadmill (model 802; custom built in the workshop of the Zoological Institute, University of Cologne, Germany). The electrodes were connected to an amplifier (model 102; custom built in the workshop of the Zoological Institute, University of Cologne, Germany) and a stimulus isolation unit. After the animal fully recovered from anesthesia (at least 5 minutes), the minimal (threshold) current necessary to elicit local reflex responses in the ipsilateral leg was determined by injecting a double impulse of 0.2 ms duration into the saphenous nerve (average ±SD threshold current: 436.4 µA ±396; range: 120-1300 µA) and the sural nerve (average ±SD threshold current: 117.6 µA ±83.4; range: 28-230 µA) of dPVs^norm^, and the saphenous nerve (average ±SD threshold current: 215 µA ±157.9; range: 64-450 µA) and the sural nerve (average ±SD threshold current: 253.3 µA ±127.5; range: 130-500 µA) of dPVs^abl^. Following the determination of threshold currents, the current injected into the sural and saphenous nerve was set as either 1.2 times the local reflex threshold current (1.2 X threshold) or five times the local reflex threshold current (5 X threshold).

EMG signals from six muscles of the right and left leg were simultaneously recorded (sampling rate: 10 kHz). Nerve stimulations (five 0.2ms impulses at 500 Hz) were applied using the ISO-FLEX (AMPI, Jerusalem, Israel) and DS4 (Digitimer, Hertfordshire, United Kingdom) stimulation insulation units. Stimulation protocols were recorded while the mice were resting on the treadmill. EMG signals were amplified (gain 100), band-pass filtered from 400 Hz to 5 kHz, and stored on the computer using Power1410 interface and Spike2 software (Cambridge Electronic Design, Cambridge, United Kingdom). For the single nerve assay, saphenous or sural nerve were electrically stimulated, while for the paired nerves assay, both nerves were stimulated at varying delays (0,5,15,15,20,25,30,35,40,45, or 50 ms). Paired stimulation of both legs at varying delays provided evidence of an inhibitory period if there was a suppression of the motor responses from one leg following stimulation of the other leg. Varying delays were used to account for differences in response latencies to the 2 nerves within muscles and within animals.

#### Analysis of EMG responses

EMG analysis response was done using Spike2 software (Cambridge Electronic Design, Cambridge, United Kingdom). For the single nerve assay, latency to response was measured for each mouse and each muscle. For the paired nerve assay, responses (single trials) were compared to the averaged response to stimulation of a single nerve (averaged over 20 trials), termed “expected response”. Responses (expected response and response to paired nerve assay) were aligned using the onset of the stimulation artifact, or the time when it was expected to occur, when the artifact was not visible. When “expected response” possessed higher amplitude than the paired stimulation response, the trial was considered to include an inhibitory component (originating from the nerve that was not used to evoke the expected response). Next, percent inhibition, the percentage of trials on which inhibition occurred was calculated for each muscle, and each mouse. For quantification of percent inhibition, a single delay (used in the paired nerve assay) was chosen, which could vary between muscle and mice. Delay at which the peak amplitude of the expected response was located 10-20 ms after the second nerve stimulation was chosen for further analysis, as this latency was shown to involve an inhibitory pathway^94,96^.

#### Behavioral testing

7-12 weeks old male and female dPV^norm^ (PV^FlpO^/*Lbx1^Cr^*^e^;*Tau^LSL-F-nLacZ-F-DTR^*) and dPV^abl^ (PV^FlpO^;*Lbx1^Cre^*;*Tau^LSL-F-nLacZ-F-DTR^*) mice injected with DTx (at P21-P28) were used for behavior testing .The experimenter was blind to the genotype of the animals. For all behavior assays, at least 7 days prior testing mice were transferred to the holding area adjacent to the behavior rooms. On the day of training/testing, mice were transferred to the behavior room for a 30-minute habituation.

#### Balance beam

Mice were trained for 2 days to cross an elevated, 1m long, 10 mm wide rectangular beam. On test day, mice were allowed to cross wide and narrow (10 mm and 5 mm, respectively) rectangular and circular beams. Individual trials were video recorded at 30 frames per second (fps) using GoPro camera, capturing mouse performance from one side. A slightly angled mirror, positioned on the other side of the mouse, was used to capture the performance of the limbs that did not face the camera. Each mouse performed 3 consecutive trials on each beam and the total number of slips and time to cross the beam were measured. All beams were custom made.

#### Horizontal ladder

Mice were trained for 2 days to cross a 60 cm long, horizontal ladder (Maze Engineers, Skokie, USA). The ladder was elevated from the ground with two cages, a neutral cage at the start and homecage at the end, to motivate mice to cross the ladder. The ladder consisted of 2 transparent side walls, with drilled holes in regularly spaced, 1 cm intervals, allowing for the insertion of metal bars (0.3 cm wide) at different spacing. During training days, ladder rungs were spaced 1 cm apart (regular pattern ladder) and mice performed 3-5 consecutive trials of ladder crossing. On the test day, mice performed 3-5 trials on a regular pattern ladder. Mice were allowed to rest for 2-3 hours, then performed 3 trials on an irregular pattern ladder where some rungs were spaced 2 cm apart. 3 different irregular patterns were used for each mouse which were consistent across mice. A camera (PROMON U1000, AOS Technologies, AG, Switzerland)) located to the side of the mouse and positioned in an angle to capture the performance of all 4 paws was used to videotape mouse performance at 30 fps. Videos were analyzed frame by frame to quantify the number of slips.

#### Rotarod test

The rotarod assay lasted for 5 days. On the first day, mice were habituated to the apparatus and placed on the rotarod (rotating at 4 rpm) for a maximum of 3 minutes. On following days, mice were put on the accelerating rotarod (4-40 rpm, maximum of 3 minutes), and their latency to fall and speed at fall were measured. Mice performed 4 trials each day, with a break of 15-20 minutes between trials.

#### Analysis of step cycle parameters

Mice were allowed to walk on a motorized treadmill (Mouse specifics, inc.) with a transparent belt at speeds 10-100 cm/s for 2 days. Each mouse had to exhibit a minimum of 3 second walk at a certain speed, for it to be included in the analysis. The second day was used to allow mice to walk at speeds they failed at on the first day. A high-speed camera (165 fps) located underneath the belt, recorded mice’ paw placement. A custom-developed software was used to extract the area of the paw that was in contact with the belt for each paw, and in each video frame, as described in^197^. These data were analyzed in MATLAB, to calculate stance duration (when paw area > 0), swing duration (paw area = 0), stride duration (stance duration + swing duration) and stride frequency (number of strides per second) of each limb. To calculate stance-to-swing phase transition, or interlimb coordination, each step cycle was normalized to values of 0 (beginning of stance) to 1 (end of swing). Phase transition between stance to swing was considered as the time in which the paw area (normalized to maximal paw area) changed from values greater than 0 to 0. Limb coupling phase values were calculated as the time difference between stance onset of the tested limb and the reference limb (left hindlimb for ipsilateral, diagonal, and hindlimb coupling and left forelimb for forelimb coupling) divided by the duration of the step cycle of the reference limb. Phase values of 0/1 (+0.25), 0.5 (+0.25), and 0.25/0.75 (+0.25), indicated in phase (synchronization), anti-phase (alternation), or out-of-phase limb coupling, respectively. Gaits were classified as walk, trot, gallop or bound and their occurrence, persistence, and attractiveness were calculated as described previously^198^. For a given gait, occurrence was calculated as its percentage out of all gaits. Persistence was calculated as the percentage of two consecutive steps using that gait out of all consecutive steps. Attractiveness was the likelihood of other gaits transferring to the gait in question.

#### Estimation of joint angles

One day before the test, the hair above the right hindlimb of mice was removed. On test day, mice were habituated to the test room for 30 minutes, and under brief anesthesia 5 marker dots (oil-based white marker, 208423, Office Depot) were placed on the following joints of their right hindlimb: iliac crest, hip, ankle, metatarsophalangeal (MTP) and toe. Due to excess skin above the knee, knee joint location was estimated post-hoc using measurements of tibia and femur length. 6 calibration points with a known distance (5 mm apart) were put on the upper right side of the transparent wall of a motorized treadmill (Mouse specifics, inc.). Mice were placed on the treadmill to recover from anesthesia (∼5 minutes), and then let to walk at 20 cm/s. A high speed camera (PROMON U1000, AOS Technologies, AG, Switzerland) recorded mice performance from the side, at 415 fps. Videos were analyzed with DeepLabCut for 2D estimation of joint marker location in each video frame as previously described^106^. Briefly, 2 separate models were created for the calibration points and joint markers. Each model was trained with a random subset of videos. An average of 125 frames were extracted and labeled by clustering based on similarities in visual appearance (k-means). The models were trained with a ResNet-50-based neural network for an average of 300,000 training iterations. Using a p-cutoff of 0.9, the training datasets were shuffled with the default 95/5 split for training and testing and evaluated with an average train error: 2.16 pixels, test: 2.38 pixels. Incorrectly labeled or missing labels were detected and corrected via extraction of outlier frames, refinement of label errors, and subsequent retraining of the model. With minimal retraining, models were able to estimate and yield 2D pixel coordinates. We used the estimated pixel coordinates of the calibration points to calculate a pixel-to-mm conversion value, then estimated the location of the knee joint in each frame using the lengths of the femur and tibia. A custom semi-automatic matlab script was used to identify the beginning of stance (the first frame on which the toe touches the ground) and swing (the first frame of paw lift-off) in each step cycle (defined from the beginning of stance to the end of swing). The complete data set of the estimated location of all 6 joints was used to generate stick diagrams of hindlimb pose and calculate the instantaneous joint angle between every three consecutive markers, throughout the step cycle.

#### Motion sequencing (MoSeq) Modeling

Moseq analysis was performed as previously described^115^. The MoSeq analysis was performed using the following programs: kinect2-nidaq (v0.2.4alpha), moseq2-extract (v1.1.2), moseq2-pca (v1.1.3), moseq2-model (v1.1.2), and moseq2-viz (v1.2.0). Data acquisition consisted of recording individual mice during 20-minute bouts of naturalistic behavior in an open arena using a Microsoft Kinect v2 camera pointed downwards at the arena. Captured depth video frames were each 512 x 424 px, collected at 30 Hz, with each pixel containing a 16-bit unsigned integer indicating the distance of that pixel from the Kinect camera sensor in mm. Raw data in each frame was then processed using a flip model supplied by the Datta lab and customized as required. In brief, the pixels corresponding to the mouse were identified, and the centroid and orientation were computed. Next, we centered and rotated the mouse within an 80 x 80 pixel image such that the mouse is always horizontally oriented with its head pointing towards the right and tail towards the left. A flip classifier (flip_classifier_k2_c57_10to13weeks.pkl), provided by the Datta Lab, was used to detect and correct any flips. Quality control consisted of human intervention ensuring that the mouse was correctly enclosed and oriented appropriately. We reduced the dimensionality of the extracted depth data via principal component analysis (PCA) by projecting the extracted depth data onto the first 10 learned PCs, resulting in a 10-dimensional time series that describes the mouse’s 3D pose dynamics. Additionally, we performed a model-free changepoint analysis, which identifies abrupt discontinuities in mouse pose dynamics and provides an empirical estimate of timescale behaviors without any model constraints. Results were assessed for quality by visual examination of the pixel weights that were assigned by each PC and the cumulative distribution of the percent variance explained by each PC. Next, we trained an autoregressive hidden Markov model (AR-HMM, aka the “MoSeq model”) consisting of a vector autoregressive process which captured the evolution of the first 10 PCs over time for each state (i.e., behavioral syllable) and a hidden Markov model describing the switching dynamics between states. A maximum of 100 behavioral states was allowed, though typically the model only discovers 40-60 states.

#### Identifying behavioral syllable usage and transition probabilities

Syllable usage (percentage) was calculated by counting the number of emissions for each syllable and dividing by the total sum of syllable emissions in each recording session. Transition matrices were calculated by counting the total number of occurrences of each syllable transitioning to any other syllable and dividing by the sum of the matrix (bigram normalization). Statistical analysis was performed as previously described^115^. In brief, group comparisons were performed for each syllable by taking 1,000 bootstrap samples of the syllable’s frequency for each group and a z-test was performed on these two distributions. The Benjamini-Hochberg procedure was used to correct for multiple hypothesis testing and control for the false discovery rate.

We performed linear discriminant analysis (LDA) using the scikit-learn implementation with the eigen solver and two components. Data was split into train and test subsets in a 70:30 ratio, and LDA models were trained using this train subset. Group labels, along with normalized syllable usages or bigram transition probabilities were inputted into the LDA model, and performance was assessed on the test subset. A permutation test was also performed (sklearn.model_selection.permutation_test_score). Here, a family of models was trained against 100 sets of randomly permuted labels. The distribution of model scores from the shuffled data was compared to the final model score to calculate a p-value. Plots were generated using seaborn and matplotlib.

#### Classifier model analysis

A multinominal logistic regression classifier was used to compare the expressive power of syllable usages (57 dimensions) or transition probabilities (3,249 dimensions). These features were randomly split into stratified training and test sets 100 times. Each training set was further split into a training and validation set using 1-fold cross validation. For each training set, a separate L1-regularized logistic regression classifier was trained to predict the labels, control or experimental, of the data. The L1 regularizer was chosen as that which had the best performance on the validation sets. The average testing F1 across all splits was 0.849 for usages, and 0.655 for transition probabilities. For each split, we tabulated the transitions which were given nonzero weight by the sparse classifier, and we took the full set of such features as an estimate of those transitions which, in general, explain the differences between control and experimental classes. We kept track of (1), the probability that a feature was given nonzero weight by a classifier and (2), if it was given nonzero weight, what that weight was. These two values, probability of inclusion in the classifier’s set of nonzero weights and the average assigned weight, were taken as numerical markers of that feature’s importance.

#### Statistical analysis

All data are reported as mean values ± the standard error of the mean (SEM), unless otherwise stated. Statistical analysis was performed in GraphPad Prism (USA) or Matlab. All data was tested for normality using the Shapiro-Wilk test. For all tests *p < 0.05, **p < 0.01, ***p < 0.001, and ****≤ 0.0001.

The means of different data distributions were compared using an unpaired Student’s t test (Figures 6D, S1G,S3B, S5B, S5C, S6A, S7A), paired Student’s t test (Figures 5D, 5E, 5H, S3H), Mann-Whitney test (Figures 2E, 2F, 2H, 2L, 2M, 2N, 2O, 3F, 5O, 6G, S1B, S1F, S1H, S1I, S1J, S1K, S2E, S2F, S4E, S4F, S6B, S7A, S7B, S7C), Wilcoxon matched-pairs signed rank test (Figure 5J), Friedmans test with Dunn’s multiple comparisons (Figures 1E, S3F), 2-way ANOVA (Figures 2I, 2J, S1E, S2H, S2I), Kruskal-Wallis test with Dunn’s correction for multiple comparisons (Figure 3K, S2L, S2M, S2N), Watson’s non-parametric two-sample U test (Figure S7 D), or linear mixed effects regression model with mouse ID as a random effect and Holm-Bonferroni method was used to correct for multiple comparisons (Figures 7C, 7D, 7E, 7G, 7H, and 7I).

## DATA AND CODE AVAILABILITY

Data are available upon request from the Lead Contact, Victoria E. Abraira.

---

**Figure S1 - related to Figure 2.**
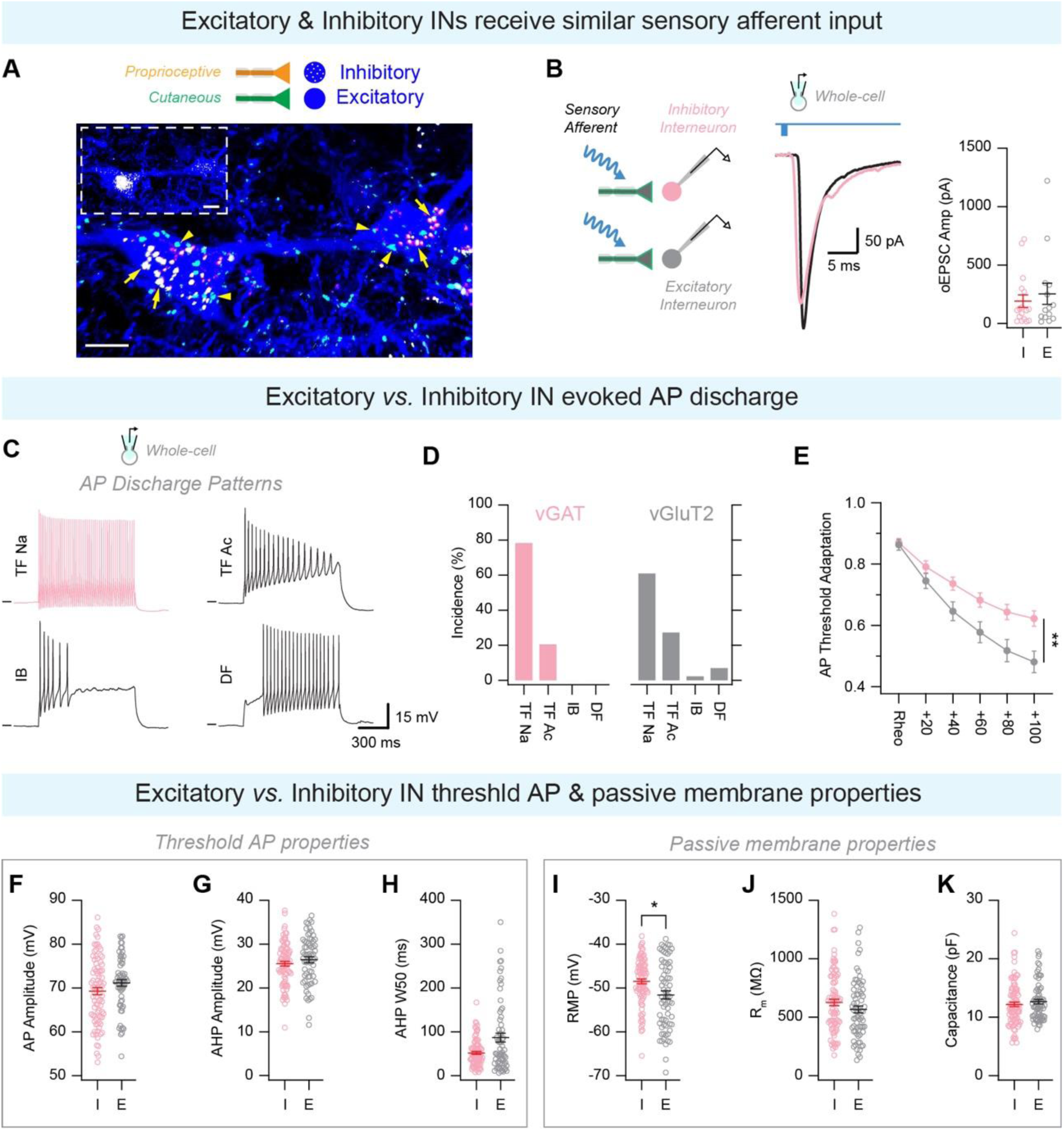
Electrophysiological characterization of deep dorsal horn inhibitory and excitatory neurons. **(A)** BFP labeled neurons within the mDDH of *PV^2aCreER^;Advillin^FlpO^;Rosa^LSL-FSF-Tomato^/Rosa^FSF-SynGFP^* mice. Inhibitory (PAX2+, white) and excitatory (PAX2-) neurons receive convergent proprioceptive (arrows) and cutaneous (arrowheads) afferent input. Scale bars: 10 µm. **(B)** Whole-cell voltage clamp recording from putative DDH inhibitory (GAD67+, pink) and excitatory (GAD67-, black) neurons following optic stimulation of sensory input (*AdvillinCre*;*R26*^LSL-ChR2-YFP^; *GAD67GFP*) showing optically evoked EPSCs (left) and oEPSC amplitude in inhibitory (I) and excitatory (E) neurons right). **(C-K)** Recordings made from vGAT^iresCre^/VGluT2^iresCre^;*R26*^LSL-tdTomato^ mice. **(C)** Representative traces of the four discharge patterns observed within the deep dorsal horn: Tonic firing non-accommodating (TF_Na_), TF accommodating (TF_Ac_), Initial bursting (IB), and Delayed firing (DF). **(D)** Incidence of each discharge pattern in inhibitory (vGAT^+^, pink) and excitatory (vGluT2^+^, gray) neurons. **(E)** Inhibitory neurons show less AP threshold adaptation. Two-way ANOVA, **p = 0052. **(F-K)** Quantification of **(F)** AP amplitude; **(G)** AHP amplitude; **(H)** AHP half width; **(I)** Resting membrane potential (RMP), Mann Whitney test; *p = 0.0102; **(J)** Membrane resistance (Rm); and **(K)** Capacitance.

**Figure S2 - related to Figure 3.**
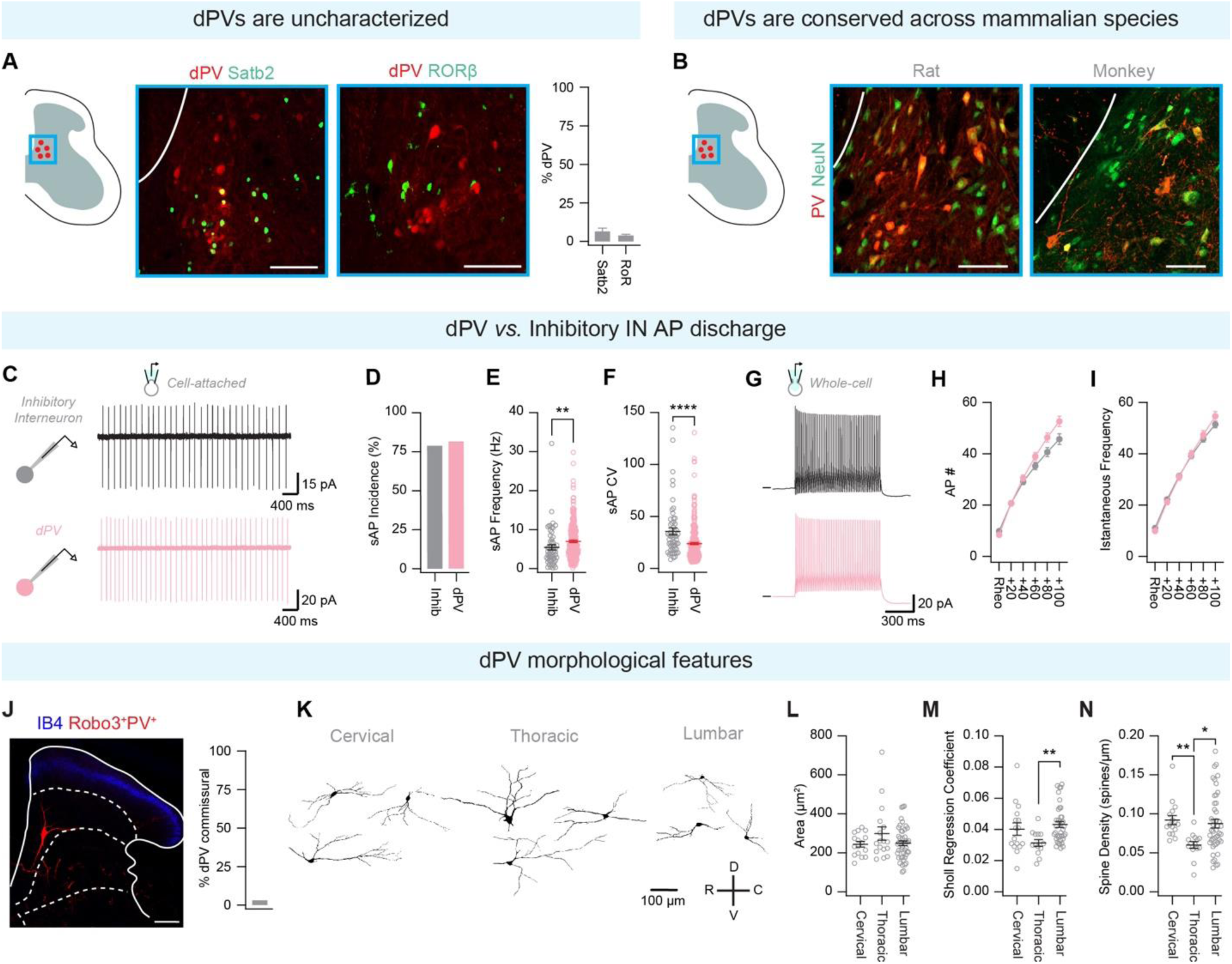
Previously uncharacterized, dPVs electrophysiology and morphology suggest a role in input integration. **(A)** Images showing colocalization of dPVs (red) and Satb2+ neurons (Green, left, intersection:*Satb2^CreER^*; *Tau^LSL-F-LacZ-F-DTR^*; PV^TdTomato^, n = 2) and RoRβ+ neurons (Green, middle, intersection; *RoRβ^FlpO^;R26*^LSL-tdTomato^ (stained with GP-PV), n = 2). Scale bars: 100 µm. Right: Percent of dPVs colocalizing with Satb2 and RoRβ. **(B)** Images showing dPVs in rat (left) and monkey (right) stained with GP-PV (red) and MS-NeuN (green). Scale bars: 100 µm. **(C-I)** Recordings made from vGAT^iresCre^;*R26*^LSL-tdTomato^ and PV^TdTomato^ mice. **(C)** Cell-attached voltage clamp recording showing spontaneous AP discharge in inhibitory (black) and dPV (pink) neurons. **(D)** Percentage of neurons exhibiting spontaneous AP discharge. **(E)** Quantification of sAP frequency; Mann-Whitney test, **p = 0.0033. **(F)** Quantification of sAP coefficient of variation (CV); Mann-Whitney test, ****p < 0.0001. **(G)** Whole-cell current clamp recording from inhibitory (black) and dPV (pink) neurons in response to depolarizing current injection. **(H)** Quantification of AP number. **(I)** Quantification of AP instantaneous frequency. **(J)** Image from *PV^FlpO^;Robo3^iresCreER^*;*R26^LSL-FSF-Tomato^* showing few dPVs express Robo3. Stained for IB4 (blue). Scale bar: 200 µm. Quantification was performed on *PV^iresCre^*;*Robo3^iresFlpO^*;*R26^LSL-FSF-Tomato^* **(K)** Representative reconstructions of dPVs at cervical, thoracic, and lumbar segments. **(L)** Soma size. **(M)** Sholl regression coefficient, Krusal Wallis test with Dunn’s correction for multiple comparisons, lumbar dPVs *vs.* thoracic dPVs, ** p = 0.0073). **(N)** Spine density, Krusal Wallis test with Dunn’s correction for multiple comparisons, cervical dPVs *vs.* thoracic dPVs, ** p = 0.0012; lumbar dPVs vs thoracic dPVs, * p = 0.0105) between cervical, thoracic, and lumbar dPVs.

**Figure S3 - related to figure 3.**
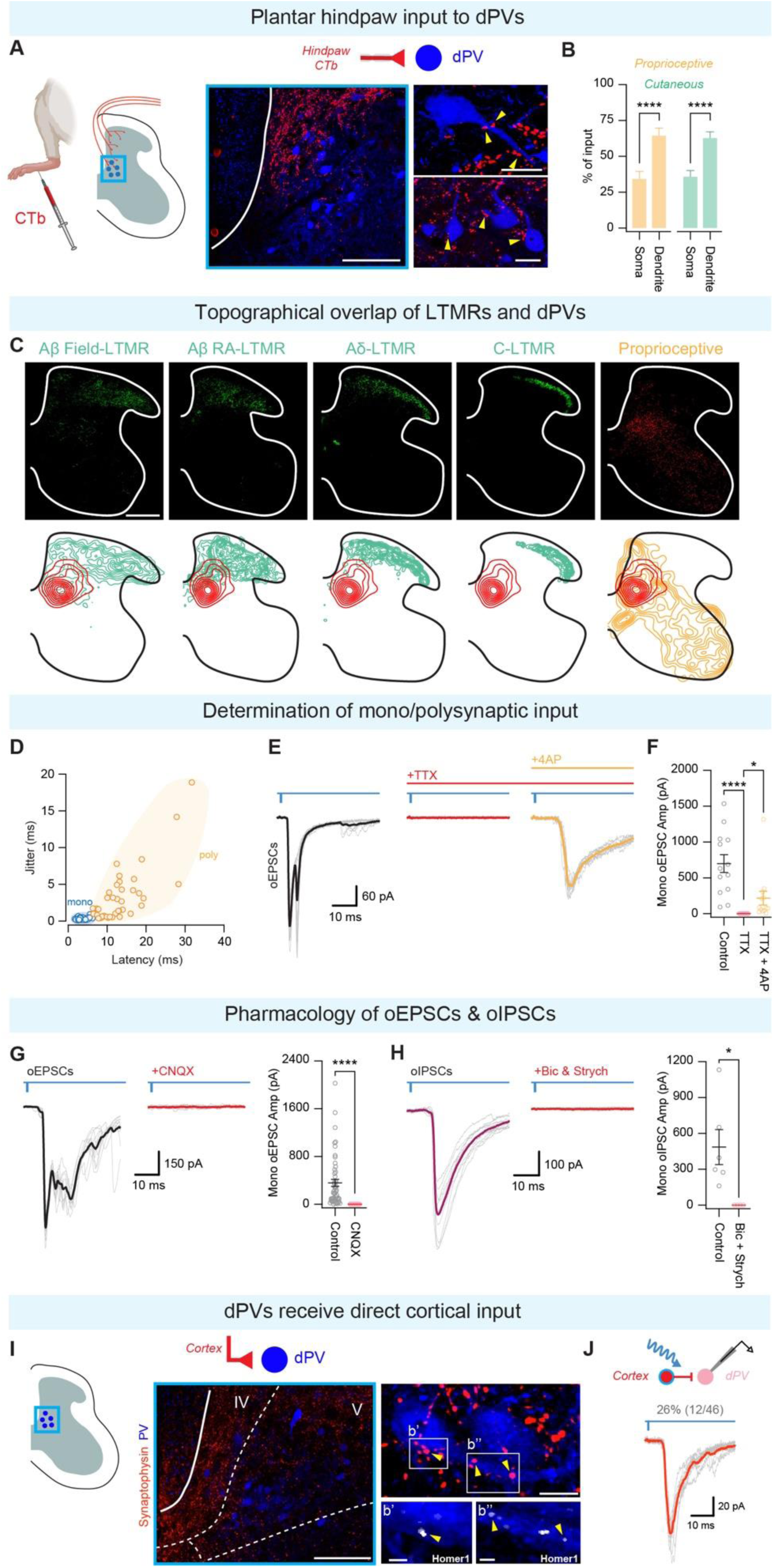
dPVs integrate diverse inputs through direct paw sensory afferents, local interneurons, and corticospinal neurons. **(A)** CTb was injected to the ventral paw of a WT animal to label paw sensory input. Tissue was immunostained with Goat-CTb (red) and Guinea pig -PV (blue). Low (left, scale bar: 100 µm) and high (right, scale bar: 20 µm) resolution images show that CTb labeled-paw afferents contact dPVs (arrowheads). **(B)** Distribution of proprioceptive and cutaneous input onto dPVs soma and dendrites (n = 2 *PV^2aCreER^;Advillin^FlpO^;Rosa^LSL-FSF-Tomato^/Rosa^FSF-SynGFP^* mice; Mann Withney test with Holm-Sidak correction for multiple comparisons,****p < 0.0001. Cutaneous input; Mann Withney test with Holm-Sidak correction for multiple comparisons, ****p < 0.0001. **(C)** Left to right - confocal (top) and contour density plots (bottom) showing the termination pattern of axon terminals of Aβ-Field LTMRs (PTGFR^CreER^;*R26^FSF-LSL-SynGFP^*, 2 mg tamoxifen treatment at P21-P22), and Aβ-RA LTMRs (*Ret^CreER^*;*R26^FSF-LSL-SynGFP^*, 2 mg tamoxifen treatment at E10-E11), Aδ-LTMRs (*TrKB^CreER^*;*R26^FSF-LSL-SynGFP^*, 2 mg tamoxifen treatment at P21), C-LTMRs (*TH^2aCreER^*;*Advillin^FlpO^*;*R26^FSF-LSL-SynGFP^*, 2 mg tamoxifen treatment at P21-P23), and proprioceptors (*PV^2ACreER^*;*Advillin^FlpO^*;*R26^FSF-LSL-SynGFP^*, 2 mg tamoxifen treatment at P21-P23), relative to the location of dPVs (Red: *PV^FlpO^*;*Lbx1^Cre^*;R26^LSL-FSF-TdTomato^) in the lumbar spinal cord. Scale bars: 200 µm. **(D)** Quantification of light induced currents with latency and jitter used to determine mono- or poly-synaptic transmission. **(E)** Whole-cell voltage clamp recordings showing oEPSCs with subsequent addition of TTX (red) and 4-AP (orange). **(F)** Quantification of pharmacological effect on monosynaptic oEPSC. Addition of 4-AP in the presence of TTX isolates monosynaptic oEPSCs. Control *vs*. TTX, ****p < 0.0001 and TTX *vs*. 4AP, *p = 0.0181; Friedmans test with Dunn’s multiple comparisons. **(G)** Left: Whole-cell voltage clamp recordings showing oEPSCs with subsequent pharmacological block with CNQX. Right: Quantification of pharmacological effect on monosynaptic oEPSCs; Mann-Whitney test, ****p < 0.0001. **(H)** Left: Whole-cell voltage clamp recordings showing oIPSCs with subsequent pharmacological block with Bicuculline and Strychnine. Right: Quantification of pharmacological effect on monosynaptic oIPSCs; paired t-test, *p = 0.0488. **(I)** Low (left, scale bar: 100 µm) and high (right, scale bar: 10 µm top, and 2 µm bottom) resolution images from *Emx1^Cre^*;*R26^LSL-syn-tdTomato^* mice stained with GP-PV and Rb-Homer1. **(J)** Whole-cell voltage clamp recording from dPV showing oEPSCs following activation of Emx1^+^ neurons (*Emx1^Cre^*;*R26^LSL-ChR2-YFP^)*.

**Figure S4 - related to Figure 4.**
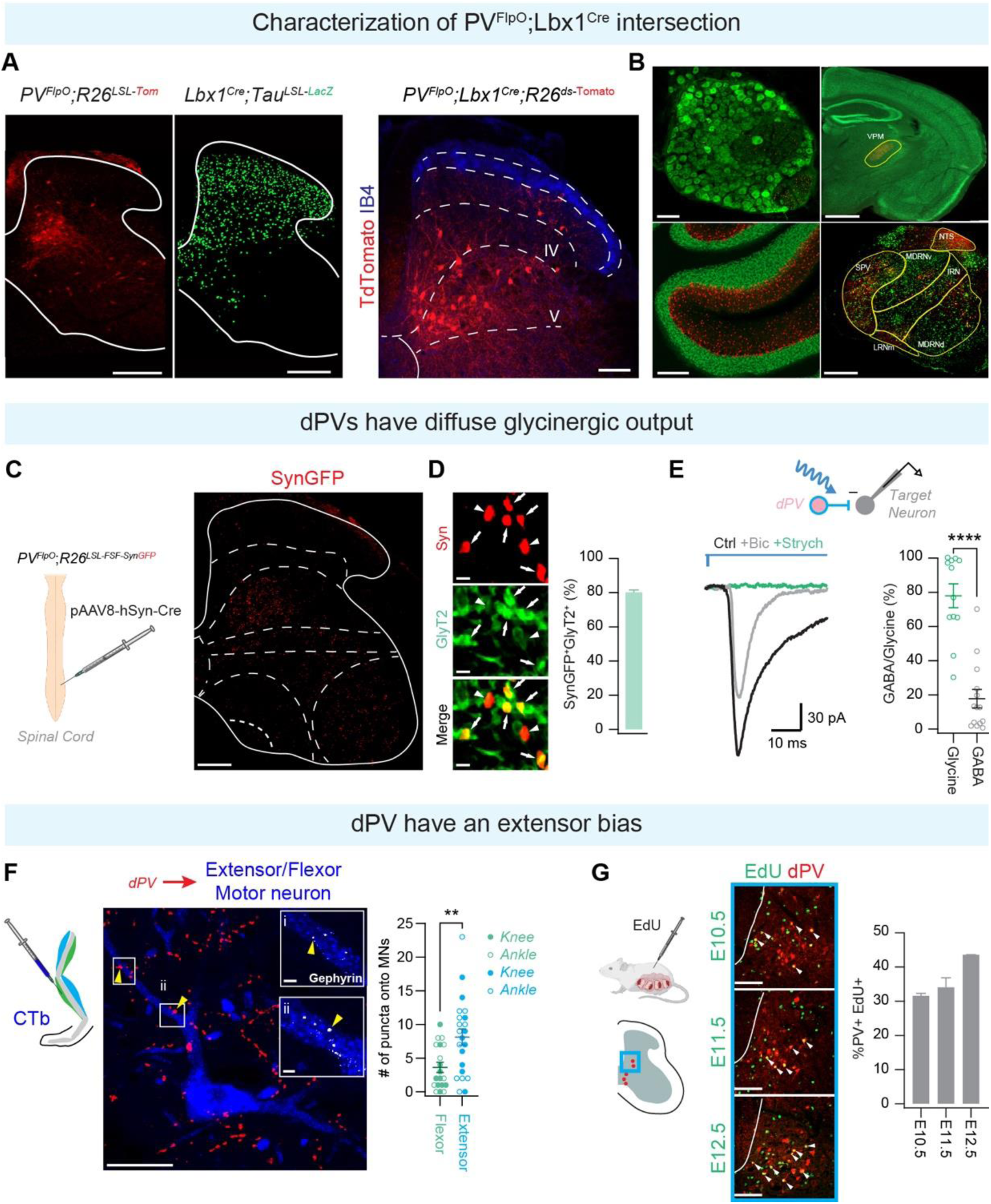
*PV^FlpO^;Lbx1^Cre^* restricts expression to spinal cord dPVs. **(A)** Transverse sections of *PV^FlpO^*;*R26^LSL-tdTom^* (left), *Lbx1^Cre^*;*Tau^LSL-F-LacZ-F-DTR^* (middle), and *PV^FlpO^*;*Lbx1^Cre^*;*R26^LSL-FSF-tdTomato^* (right). Scale bars: left and middle 200 µm, right 100 µm. **(B)** Images from *PV^FlpO^*;*Lbx1^Cre^*;*R26^LSL-FSF-tdTomato^* intersection in the DRG (top left, stained for NFH, scale bar 100 µm), ventral posteromedial nucleus (VPM, top right, stained for NeuN. scale bar 1000 µm), molecular layer of the cerebellum (bottom left, stained for NeuN, scale bar 200 µm), and the brainstem (bottom right, spinal nucleus of the trigeminal (SPV), nucleus of solitary tracts (NTS), ventral medullary reticular nucleus (MDRNv), dorsal medullary reticular nucleus (MDRNd), and intermediate reticular nucleus (IRN), stained for NeuN, scale bar 500 µm). **(C)** Transverse section of lumbar spinal cord from a *PV^FlpO^*;*R26^LSL-FSF-SynGFP^* mouse injected with pAAV8-hSyn-Cre. Scale bar 200 µm. **(D)** Left: Images from *PV^FlpO^;Lbx1^Cre^;R26^LSL-FSF-SynGFP^* mice show dPV terminals (red) colocalize with GlyT2+ (green, stained with GP-GlyT2). White arrows show dPV terminal colocalization with GlT2, arrowheads show no colocalization. Scale bar: 2 µm. Right: Percentage of dPV terminals that express GlyT2. **(E)** Whole-cell voltage clamp recording showing oIPSCs following dPV photostimulation (*PV^FlpO^;Lbx1^Cre^;R26^LSL-FSF-ChR2-YFP^)* and following bath application of bicuculline (gray) and strychnine (green). Right: Quantification of GABA/glycine contribution to dPV-evoked oIPSCs; Mann Whitney test. ****p < 0.0001. **(F)** Injections of the retrograde tracer Alexa fluor 555-conjugated CTb were made into extensor and flexor muscles of *PV^FlpO^*;*Lbx^1Cre^*;*Rosa^LSL-FSF-SynGFP^* mice. n = 10 motor neurons in 2 vastus lateralis (knee extensor) injected mice, n = 10 motor neurons in 2 biceps femoris (knee flexor) injected mice, n = 12 motor neurons in 1 lateral gastrocnemius (ankle extensor) injected mouse, and n = 9 motor neurons in 1 tibialis anterior (ankle flexor) injected mouse. Scale bars: 40 µm. Inset: Apposition to Gephyrin (white, arrowheads). Scale bar: 2µm. Right: Quantification of dPV inputs onto flexor *vs.* extensor motor neurons; Mann-Whitney test, **p = 0.0044. **(G)** Left: Images of dPVs (red) from PV^TdTomato^ mice injected with EdU (green) at E10.5 (top), E11.5 (middle) and E12.5 (bottom), arrowheads show colocalization. Right: Percentage of dPVs colocalized with EdU (n = 2 animals for each day).

**Figure S5 - related to Figure 6.**
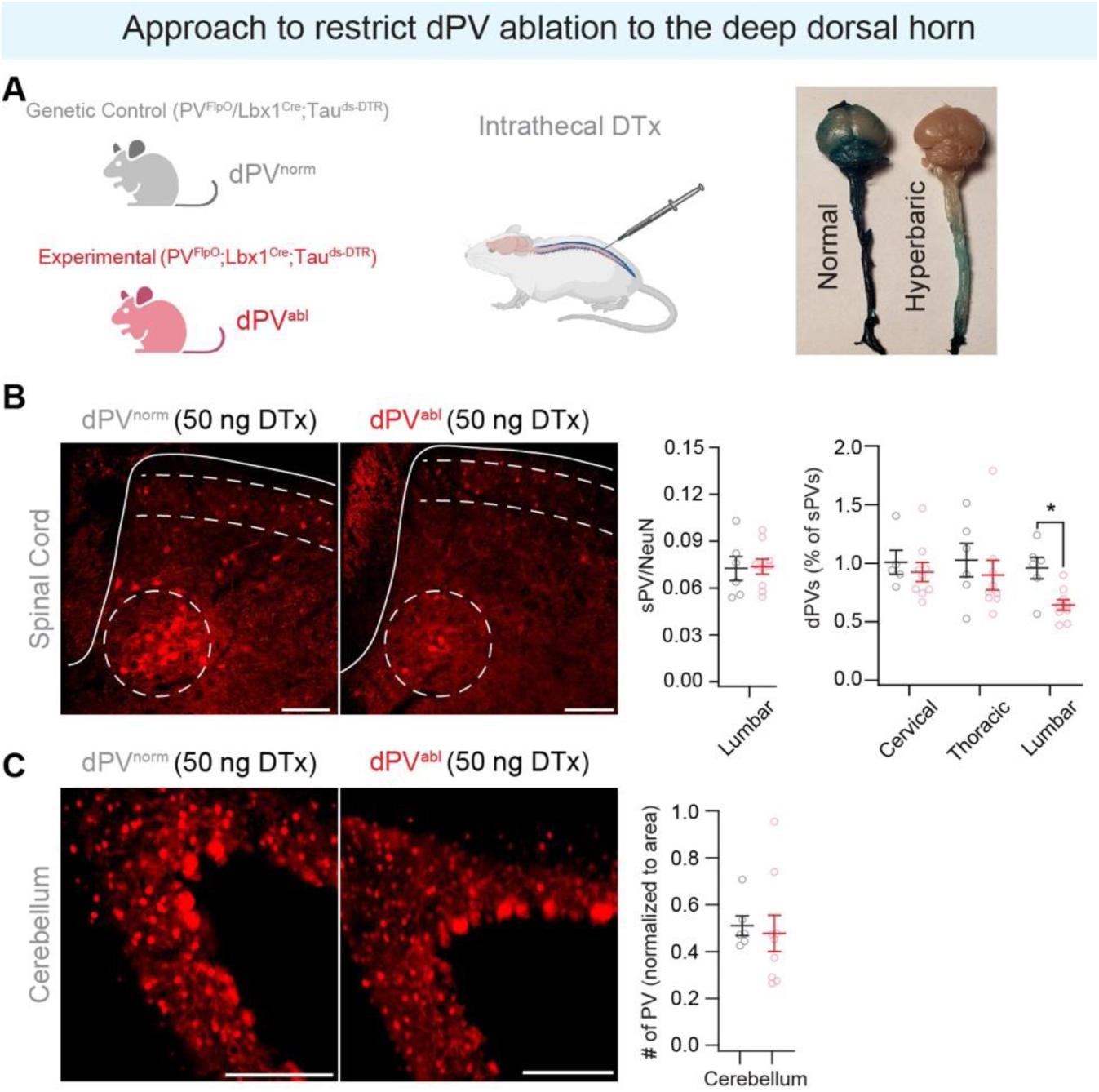
Approach for dPV ablation. **(A)** Strategy for dPVs ablation. Left: *PV^FlpO^*;Lbx1^Cre^;*Tau^LSL-F-LacZ-F-DTR^* and *PV^FlpO^*;*Tau^LSL-F-LacZ-F-DTR^* or *Lbx1^Cre^*;*Tau^LSL-F-LacZ-F-DTR^* mice were injected with 50 ng Diphtheria toxin (DTx) to generate animals with ablated dPVs (dPV^abl^) and non-ablated dPVs (dPV^norm^). Right: Intrathecal injection of 20% fast green made in PBS (left) or hyperbaric solution (right) to a WT mouse, demonstrates that the latter restricts spread to the spinal cord. **(B)** Left: Images showing lumbar spinal cord of dPV^norm^ (left) and dPV^abl^ (right). Scale bars 200 µm. Right: Quantification of dPV (circle) and sPV neurons (dotted lines) in dPV^norm^ and dPV^abl^ mice following I.T injection of 50 ng DTx. **(C)** Left: Images showing molecular layer of cerebellum of dPV^norm^ (left) and dPV^abl^ (right). Scale bars 200 µm. Right: Quantification of PV neurons in dPV^norm^ and dPV^abl^ mice following I.T injection of 50 ng DTx. n = 6 dPV^norm^ and n = 9 dPV^abl^. Mann Whitney test, *p = 0.02557.

**Figure S6 - related to Figure 6.**
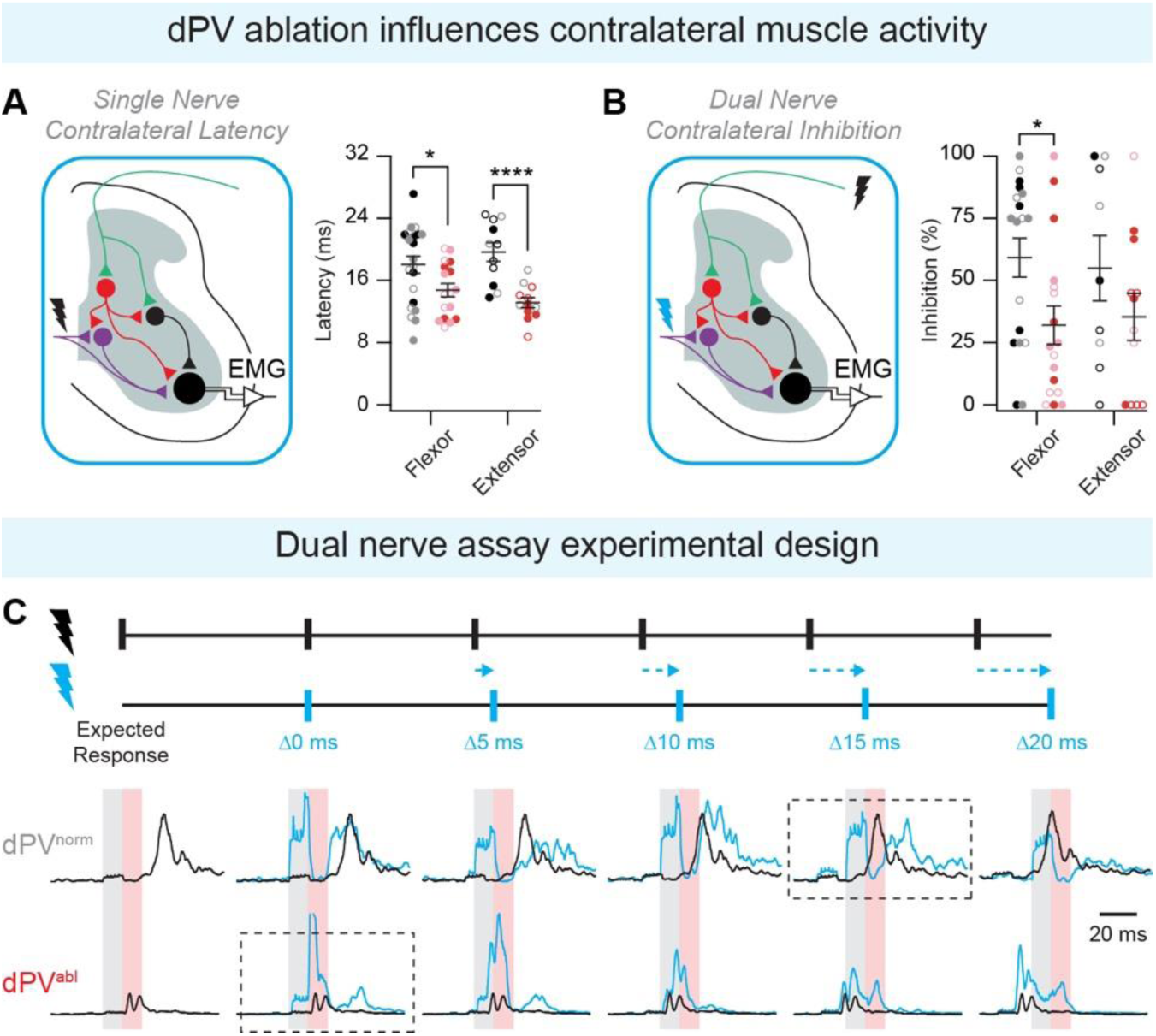
dPV ablation alters muscle response to contralateral cutaneous stimulation. **(A)** Left: Experimental setup designed to reveal the contribution of dPVs to flexor/extensor muscle activity following contralateral cutaneous nerve stimulation. Right: Quantification of latency to EMG responses following contralateral cutaneous nerve stimulation. EMG recordings were obtained from 6 dPV^norm^ (*PV^FlpO^*/*Lbx1^Cre^*;*Tau^LSL-F-LacZ-F-DTR^*) and 9 dPV^abl^ (*PV^FlpO^*;*Lbx1^Cre^*;*Tau^LSL-F-LacZ-F-DTR^*) mice intrathecally injected with 50 ng DTx. We observed no difference between Sural *vs.* Saphenous nerve stimulation in paired muscles: dPV^norm^ ST, Unpaired t-test, p = 0.1866; dPV^abl^ ST, Unpaired t-test, p = 0.7207; dPV^norm^ GS, Unpaired t-test, p = 0.6714; dPV^abl^ GS, Unpaired t-test, p = 0.3977. Therefore, we combined sural and saphenous nerve stimulation. dPV^abl^ mice exhibit shorter cutaneous evoked latency in flexors (n = 17 dPV^norm^ and 21 dPV^abl^): Unpaired t-test, **p < 0.0299, and extensors (n = 11 dPV^norm^ and n = 12 dPV^abl^): Unpaired t-test, ****p = <0.0001. **(B)** Left: Experimental setup designed to reveal the contribution of dPVs to inhibition of flexor/extensor muscle activity following contralateral cutaneous nerve stimulation. Right: Percentage of trials exhibited cutaneous-evoked inhibition (Single nerve response > Paired nerve response) following contralateral cutaneous nerve stimulation. We observed no difference between Sural *vs.* Saphenous nerve stimulation in paired muscles: dPV^norm^ ST, Unpaired t-test, p = 0.6410; dPV^abl^ ST, Unpaired t-test, p = 0.7942; dPV^norm^ GS, Unpaired t-test, p = 0.3322; dPV^abl^ GS, Mann-Whitney test, p = 0.2679. Therefore, we combined sural and saphenous nerve stimulation. dPV^abl^ mice exhibit reduced contralateral cutaneous evoked inhibition in flexors (n = 18 dPV^norm^ and 17 dPV^abl^): Mann-Whitney test, *p < 0.0308, and no change in extensors (n = 9 dPV^norm^ and n = 8 dPV^abl^): Mann-Whitney test, p = 0.1282. **(C)** Dual nerve assay experimental design. Stimulation of a single nerve is performed (black) to calculate the “expected response”. Both cutaneous nerves are stimulated at varying delays (blue). Traces show example EMG responses recorded in the right gastrocnemius (Gs) of dPV^norm^ (left) and dPV^abl^ (right). Top row shows the “expected response” to stimulation of the left saphenous nerve (black), averaged over 20 trials. Rows 2-6 show responses to paired stimulation of the left saphenous nerve and right sural nerves (cyan), at delay of 0, 5, 10, 15, and 20 ms, respectively, overlaid on the expected response. Grey shading highlights second nerve stimulus artifact, red shading highlights analysis window, dotted boxes show stimulus delay chosen for analysis (aligned response).

**Figure S7 - related to Figure 7.**
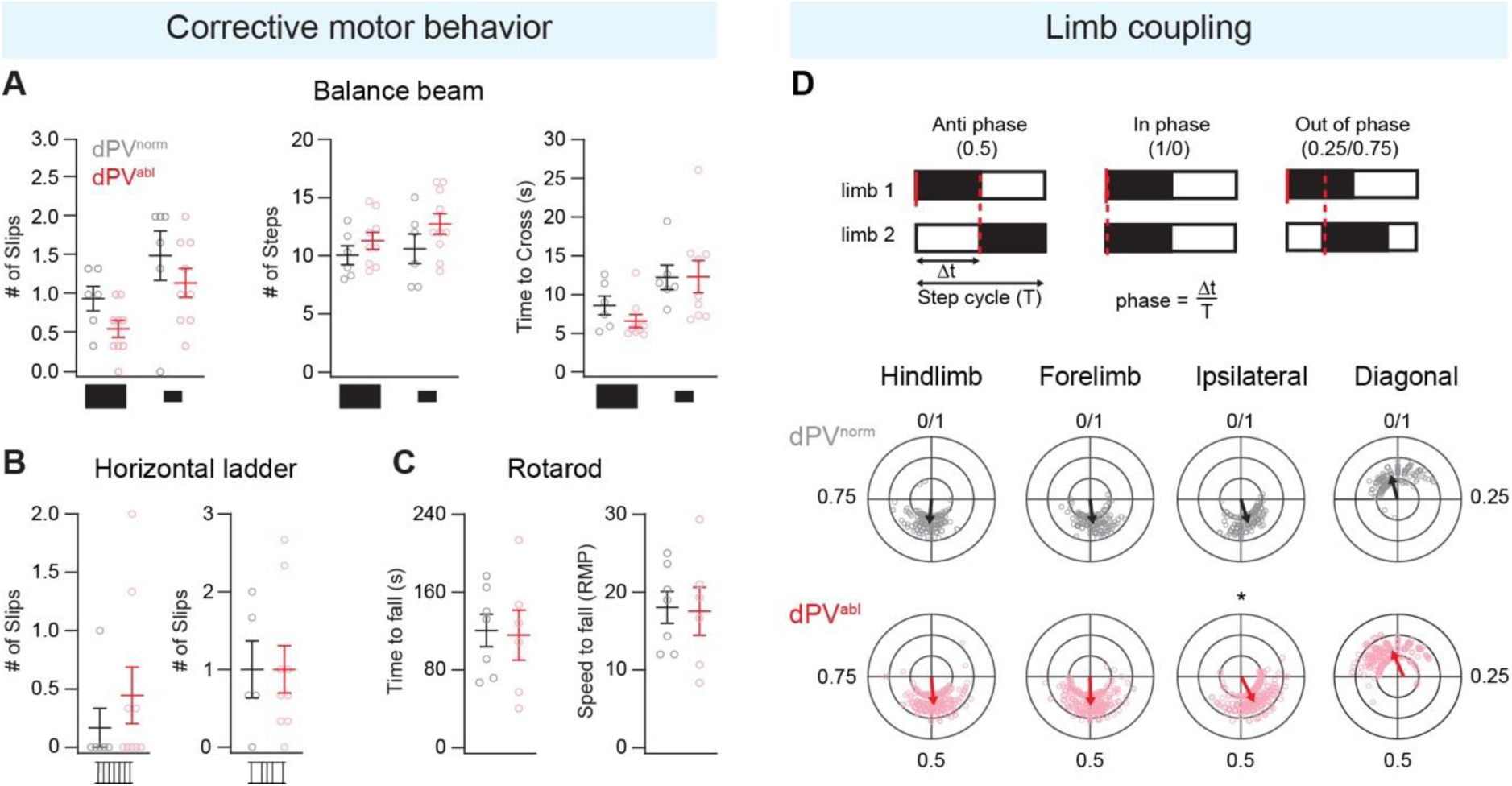
Ablation of dPVs does not alter corrective motor reflexes or limb coupling. **(A)** Quantification of number of slips (left) and number of steps (right) on rectangular 10 mm (n = 6 dPV^norm^ and n = 9 dPV^abl^), rectangular 5 mm (n = 6 dPV^norm^ and n = 9 dPV^abl^), circular 10 mm (n = 6 dPV^norm^ and n = 10 dPV^abl^), and circular 5 mm balance beams (n = 6 dPV^norm^ and n = 10 dPV^abl^). **(B)** Quantification of the number of slips on a horizontal ladder with rungs in a regular pattern (left: n = 6 dPV^norm^ and n = 9 dPV^abl^) or irregular pattern (right: n = 5 dPV^norm^ and n = 9 dPV^abl^). **(C)** Quantification of time to fall and speed at fall from the rotarod (4-40 rpm, maximum of 3 minutes: n = 7 dPV^norm^ and n = 6 dPV^abl^). **(D)** Top: diagram depicting phase coupling calculation. Δt, the time between the initial contact of a limb (limb 2) and a reference limb (limb 1), is divided by the duration of the step cycle (T). Step cycle duration is calculated for the reference limb from initial contact (beginning of stance) to lift-off (end of swing). Phase values of 0.5 (+0.25), 1/0(+0.25), and 0.25/0.75 (+0.25) indicate that limbs are in anti-phase, in-phase or out-of-phase, respectively. Bottom: Comparison of phase-frequency plot of 7 dPV^norm^ and 10 dPV^abl^ mice for (from left to right) hindlimb coupling, forelimb coupling, ipsilateral coupling, and diagonal coupling. Watson’s U^2^ for circular data, hindlimb: *p = 0.0390.

**Figure S8 - related to Figure 7.**
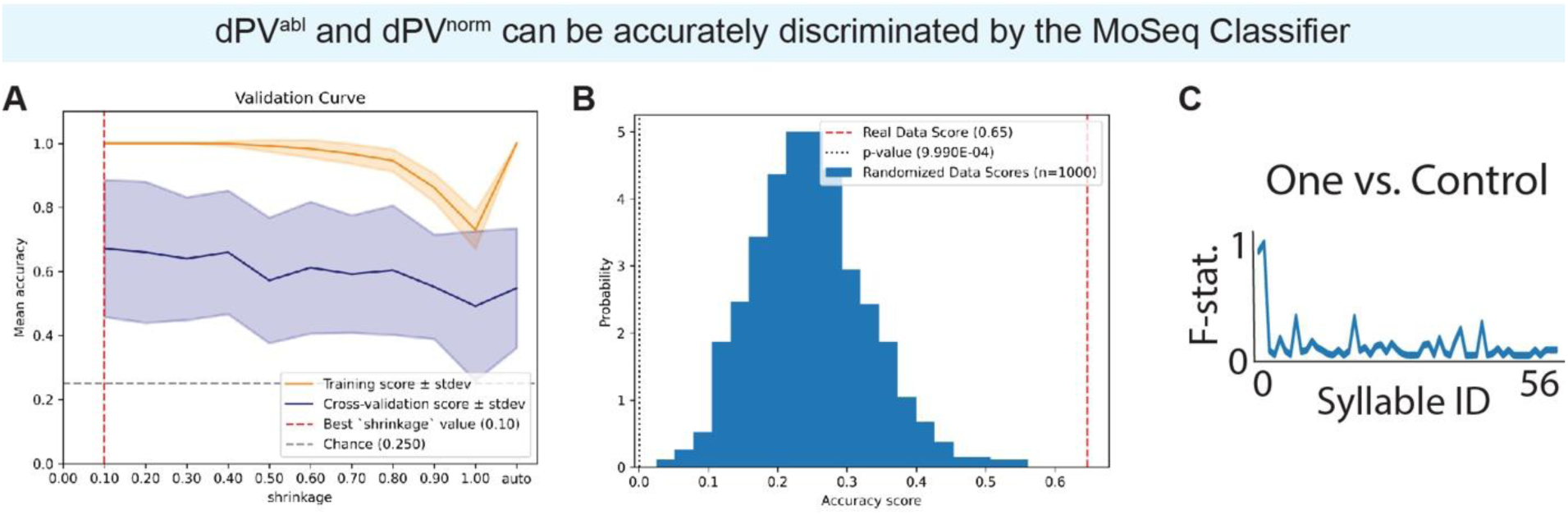
Behavioral modeling reveals distinct phenotypes in animals with dPV ablation. **(A)** Cross validation curve for selection of the shrinkage hyperparameter for LDA models using 5-fold cross validation. Orange line shows the training set performance (i.e., model score on training data). Blue line is the score on held-out data. Grey dotted line represents the level of chance (probability of success if a class is chosen at random). **(B)** Results of a permutation test, generating a family of 100 models trained against randomly permutated labels (histogram bars). The p-value is calculated using the distribution of model scores for shuffled data *vs.* the final model. **(C)** Normalized F-statistic reveals the relevance of each identified syllable in discriminating control and dPV ablated (“one”) groups. Results were plotted with seaborn and matplotlib.

**Table S1 – related to Figure 8.** Logistic regression reveals select behavioral syllables that carry large weights in discriminating groups. Logistic regression identified select syllables that can be used to distinguish dPV ablated animals from controls.

| Syllable | Avg weight to classifier | Probability of inclusion in the classifier |
| --- | --- | --- |
| 1 | -131.81 | 1.00 |
| 9 | 51.68 | 0.89 |
| 0 | 103.85 | 0.78 |
| 18 | 32.75 | 0.47 |
| 12 | 46.96 | 0.35 |
| 14 | 39.14 | 0.30 |
| 30 | -29.50 | 0.29 |
| 33 | -17.35 | 0.15 |
| 21 | -126.76 | 0.10 |
| 8 | 13.51 | 0.09 |
| 19 | -65.89 | 0.08 |
| 20 | 41.11 | 0.06 |

**Table S2 – related to Figure 7.** dPV ablation results in a loss of motor behavior complexity. Logistic regression reveals select behavioral syllable transitions that carry large weights in discriminating dPV ablated animals from controls. Nearly all the computationally identified syllable transitions are downregulated in the dPV ablation group.

| Syllable transitions | Up or down regulated | Avg weight to classifier | Probability of inclusion in the classifier | Adj p (bootstrap) |
| --- | --- | --- | --- | --- |
| 1 → 30 | Down | -292.66 | 0.75 | 0.41 |
| 1 → 17 | Down | -241.75 | 0.67 | 0.12 |
| 17 → 1 | Down | -412.72 | 0.53 | 0.00 |
| 21 → 19 | Down | -397.32 | 0.48 | 0.25 |
| 18 → 22 | Up | 42.60 | 0.34 | 0.40 |
| 39 → 40 | Down | -62.46 | 0.32 | 0.97 |
| 8 → 20 | Down | -77.54 | 0.23 | 0.26 |
| 1 → 14 | Down | -117.93 | 0.21 | 0.16 |
| 20 → 12 | Down | 164.48 | 0.19 | 0.78 |
| 4 → 30 | Up | 53.05 | 0.19 | 0.64 |
| 4 → 1 | Down | -376.51 | 0.15 | 0.05 |
| 3 → 21 | Down | -252.18 | 0.13 | 0.48 |
| 10 → 29 | Down | -131.95 | 0.12 | 0.50 |
| 43 → 39 | Down | -159.63 | 0.11 | 0.92 |
| 15 → 1 | Down | -217.76 | 0.1 | 0.12 |
| 8 → 10 | Down | -91.58 | 0.08 | 0.48 |
| 1 → 15 | Down | -206.05 | 0.08 | 0.123 |
| 10 → 4 | Down | 47.87 | 0.08 | 0.43 |
| 21 → 0 | Up | 40.33 | 0.07 | 0.35 |
| 4 → 26 | Down | 123.12 | 0.06 | 0.70 |
| 3 → 27 | Up | -52.96 | 0.05 | 0.69 |
| 10 → 3 | Down | -90.83 | 0.05 | 0.29 |

## Notes

### Competing Interest Statement

The authors have declared no competing interest.

### Summary of Updates

adding author

